# The amniote-conserved DNA-binding domain of CGGBP1 restricts cytosine methylation of transcription factor binding sites in proximal promoters to regulate gene expression

**DOI:** 10.1101/2024.06.20.599840

**Authors:** Ishani Morbia, Praveen Kumar, Aditi Lakshmi Satish, Akanksha Mudgal, Subhamoy Datta, Umashankar Singh

## Abstract

CGGBP1 is a GC-rich DNA-binding protein which is important for genomic integrity, gene expression and epigenome maintenance through regulation of CTCF occupancy and cytosine methylation. It has remained unclear how CGGBP1 integrates multiple diverse functions with its simple architecture of only a DNA-binding domain tethered to a C-terminal tail with low structural rigidity. We have used truncated forms of CGGBP1 with or without the DNA-binding domain (DBD) to assay cytosine methylation and global gene expression. Proximal promoters of CGGBP1-repressed genes, although significantly GC-poor, contain GC-rich transcription factor binding motifs and exhibit base compositions indicative of low C-T transition rates due to prevention of cytosine methylation. Genome-wide analyses of cytosine methylation and binding of CGGBP1 DBD show that CGGBP1 restricts cytosine methylation in a manner that depends on its DBD and its DNA-binding. The CGGBP1-repressed genes show an increase in promoter cytosine methylation alongside a decrease in transcript abundance when the DBD-deficient CGGBP1 is expressed. Our findings suggest that CGGBP1 protects transcription factor binding sites (TFBS) from cytosine methylation-associated loss and thereby regulates gene expression. By analysing orthologous promoter sequences we show that restriction of cytosine methylation is a function of CGGBP1 progressively acquired during vertebrate evolution. A superimposition of our results and evolution of CGGBP1 suggests that mitigation of cytosine methylation is majorly achieved by its N-terminal DBD. Our results position CGGBP1 DNA-binding as a major evolutionarily acquired mechanism through which it keeps cytosine methylation under check and regulates TFBS retention and gene activity.

## Introduction

Human CGG triplet repeat-binding protein CGGBP1 was identified as a protein bound to GC-rich DNA (Singh and Westermark, 2015). The human CGGBP1 gene, located on the p arm of chromosome 3, is transcribed from multiple transcription start sites with at least four well-characterised splice variants. However, all the CGGBP1 transcripts are known to encode the same 20 kD protein. Interestingly, the ORF for this 20 kD protein is restricted within one exon only. The emergence of diversity in its transcription start sites (Singh and Westermark, 2015) and transcript structures without giving rise to any alternative coding sequence suggests some evolutionary constraints on CGGBP1. It is well conserved in amniotes and highly conserved in mammals (Ensembl GeneTree). Although the tetrapod CGGBP1 seems to be derived from one of the multiple Coelacanth CGGBPs (Singh and Westermark, 2015; Yellan, Yang and Hughes, 2021), there is no evidence of any CGGBP ortholog in amphibians that is expressed as a protein. The importance of this emergence of CGGBP1 at the cusp of evolution of tetrapods remains unknown.

A comparison of CGGBPs from various tetrapod taxa suggests that the protein is composed of two roughly equally-sized parts each with a different domain (UniProt Q9UFW8). The N-terminal part contains two cysteine and histidine (C2H2) residues which seem to be well conserved and functionally selected UniProt Q9UFW8. Mutation in this part of CGGBP1 has been shown to abrogate DNA binding (Richards *et al*., 1993; Deissler *et al*., 1997; Müller-Hartmann *et al*., 2000; Agarwal *et al*., 2016). No known domains have been assigned to CGGBP1 in prominent databases but this N-terminal half has been attributed similarity to the Zinc finger BED domain (Singh and Westermark, 2015; Yellan, Yang and Hughes, 2021). The predicted structure of CGGBP1 in AlphaFold database (embedded in UniProt Q9UFW8) shows that this 10kD part of the protein is well-structured containing amphipathic helices compatible with formation of Zn fingers required for DNA binding.

The C-terminal part of CGGBP1 is not as well understood. It has no known domains, does not contain conserved residues indicative of any function and has low confidence structure prediction in AlphaFold. Mutations in this part of CGGBP1 affect nuclear localisation with variable and mild effects on DNA-binding (Deissler *et al*., 1997; Müller-Hartmann *et al*., 2000; Singh *et al*., 2014).

Sequence analyses suggest that this part of CGGBP1 is derived from hAT transposase (Singh and Westermark, 2015; Yellan, Yang and Hughes, 2021). All the available information on CGGBP1 suggest its origin out of functional exaptation of a Hermes DNA transposon in the common tetrapod ancestor.

Given that CGGBP1 is a rather recent addition to the vertebrate genomes, it is surprising that it is involved in several multiple cellular processes which are conducted by extremely well conserved proteins. It is required for stress response (Singh, Bongcam-Rudloff and Westermark, 2009; Datta *et al*., 2022), cell cycle progression (Singh *et al*., 2011) and cytokinetic abscission (Singh and Westermark, 2011), telomeric integrity and prevention of endogenous DNA damage (Singh *et al*., 2014; Datta *et al*., 2022).

A series of reports describing CGGBP1 function indicate that interactions of CGGBP1 with specific target regions in the genome are important for its functions. Some of these interactions could be direct binding of CGGBP1 to DNA. However, an absence of a DNA sequence motif signature in high-throughput assays (Agarwal *et al*., 2016; Snyder, 2021) suggests that a significant fraction of CGGBP1-DNA interactions could be indirect. The well established features of DNA with which CGGBP1 could directly interact and regulate the functions of the DNA include GC-richness and repetitive base compositions giving rise to G/C-skew (Deissler *et al*., 1997; Müller-Hartmann *et al*., 2000; Patel *et al*., 2018; Datta *et al*., 2024), such as the CGG repeats after which the protein has been named.

Direct binding of CGGBP1 to specific subsequences of Alu repeats has been shown to suppress Alu transcription and thus affect the global transcription activity by RNA Polymerase 2 (Agarwal *et al*., 2016). Phosphorylation of a tyrosine residue (Y20) flanking the N-terminal DNA-binding domain seems essential for direct binding of CGGBP1 to Alu A- and B-box oligonucleotides *in vitro*. A distinct phosphorylation of serine residues at the C-terminus, especially S164, seems to be required for collaboration between CGGBP1 and POT1 (Singh *et al*., 2014). This in turn is required for POT1 loading on the telomeres and protection of telomeric DNA from endogenous DNA damage. These findings indicate that the N-terminal part of CGGBP1 is indeed required for its direct DNA-binding while the C-terminal part functions without necessarily binding to the DNA but affects the outcomes of CGGBP1-DNA interactions. Experimental evidence for the distinct contributions of the N- and C-terminal parts of CGGBP1 to DNA-binding genome-wide and their functional outcomes remain unreported.

CGGBP1 depletion leads to a change in gene expression (Agarwal *et al*., 2016) which has hitherto been understood to be potentially due to a combination of two different mechanisms: a small set of genes getting deregulated due to loss of direct DNA-CGGBP1 interactions (Singh, Bongcam-Rudloff and Westermark, 2009; Agarwal *et al*., 2016) and a larger set of genes which are deregulated by higher order changes in RNA Polymerase activity (Agarwal *et al*., 2016), changes in histone marks and altered barriers between active and silent chromatin (Patel *et al*., 2019).

CGGBP1 is required for CTCF occupancy at LINE-1 repeats and thereby separation of active and silent chromatin marked by different levels of H3K9me3 and cytosine methylation (Patel *et al*., 2019). There is another level of gene regulation by CGGBP1 that involves aberrant transcription start and termination. This effect is observed most prominently around CGGBP1-dependent CTCF-binding sites (Patel *et al*., 2021). Proximity ligation assays and co-immunoprecipitations coupled with DNase digestion have shown that even if CTCF and CGGBP1 do not show prominent physical interaction with each other, they bind to DNA in enough proximity to exert regulatory influence on each other (Patel *et al*., 2019).

Early studies on human CGGBP1 showed that its binding to DNA in vitro was sensitive to cytosine methylation (Müller-Hartmann *et al*., 2000). Follow up investigations have shown that CGGBP1 levels affect cytosine methylation genomewide in CpG as well as non-CpG contexts (Agarwal *et al*., 2015; Patel *et al*., 2018). Cytosine methylation patterns at genomic interspersed repeats are specifically regulated by CGGBP1. Although the cytosine methylation level changes caused by CGGBP1 depletion are bidirectional, the net change is an increase. It remains unclear whether binding of CGGBP1 to target DNA sequences directly interferes with cytosine methylation or if the cytosine methylation levels change due to altered expression of genes involved in regulation of cytosine methylation (Agarwal *et al*., 2015). Evidence so far gives rise to a possibility that CGGBP1 binding to GC-rich DNA might confer steric protection against cytosine methylation at some sites.

Thus, there are multiple indications that the many functions of CGGBP1 are somehow associated with its interaction with the DNA or its chromatin occupancy. However, a systematic analysis of direct/indirect DNA-binding and its importance for CGGBP1 functions has never been reported. *In vitro* DNA-protein interaction assays have shown that CGGBPs from different taxa have different DNA-sequence motif preferences such that the binding site preferences seem to have evolved independently in different lineages (Yellan, Yang and Hughes, 2021). The preferred binding sites of CGGBPs from different species are distinct. These distinct binding site preferences could be derived out of two evolutionary phenomena. First, changes in the sequence and abundance of target sequences in the genome; but apart from GC-richness, there is no common feature that identifies the various CGGBP-binding sequences. Second, changes in CGGBPs which would lead to distinct DNA sequences as preferred binding sites. Most strikingly, the DNA-binding domain at the N-terminus of CGGBP1 is the most conserved part of the protein and it is unlikely that distinct DNA sequence preferences arose without concomitant changes in either the DNA-binding domain itself or a cooperating part of CGGBP1 that could affect the properties of the DNA-binding domain.

The Znf BED domain, proposed to be in CGGBP1 N-terminus, has been associated with gene repression, chromatin insulation and yet exhibits no known DNA sequence preference. The presumed transposase-derived C-terminal part of CGGBP1 has neither any known binding sequence preference nor any associations with cellular processes.

Here we report that dissociated N- and C-terminal truncated parts of CGGBP1 have distinct effects on global cytosine methylation and gene expression. A net pro-cytosine methylation and gene repressive is exerted by the C-terminal part of CGGBP1 devoid of the N-terminal DBD. We show that CGGBP1 C-terminus increases cytosine methylation at most regions genome-wide, including at transcription factor binding sites depending on GC content, GC-skew and evolutionary status of the cognate transcription factors. The CGGBP1-repressed genes commonly have GC-rich DNA sequence motifs embedded in their proximal promoters, which are overall GC- and CpG poor. By analysing the orthologous promoter sequences in different tetrapods we find that the restriction of cytosine methylation by CGGBP1 has minimised C to T transitions leading to retention of GC rich motifs in GC poor promoters. These motifs correspond to binding sites of known transcription repressor proteins, including CTCF. Chromatin immunoprecipitation results and DNA sequence analysis suggest that CGGBP1-binding to these GC-rich motifs sterically prevents their cytosine methylation and protects them against C-T transitions. Our results present a mechanism of gene expression regulation in which a DNA-binding protein CGGBP1 has evolved alongside its binding sites on target genes. Our interpretation of the evolution of the base composition of CGGBP1-repressed genes alongside the nature of genomic DNA occupied by CGGBP1 N-terminus provides evidence that CGGBP1 binding restricts C-T transitions, conserves transcription factor binding sites and thus regulates gene expression.

## Results

### 1. Loss of CGGBP1 DBD leads to decreased nuclear localization and global increase in cytosine methylation

Regulation of cytosine methylation seems to be a function of CGGBP1 that is fundamental to most of its other effects including maintenance of genomic integrity and structure as well as regulation of chromatin barrier activity and gene expression. In order to understand how cytosine methylation and possibly other downstream functions of CGGBP1 depend on its DNA-binding domain, we used truncated versions of CGGBP1.

We generated N-terminal (N-term, aa1-aa90) and C-terminal (C-term, aa79-aa167) truncated forms of human CGGBP1 such that they both contained the NLS (Fig 1A). To ensure the stability and subcellular visualisation of the truncated proteins, they were tagged with GFP (Fig 1A). Interestingly, even though the N-term and C-term CGGBP1 forms both contained the NLS, the C-term showed a higher extra-nuclear localisation (Fig 1B). This higher extra-nuclear presence of C-term was verified by nucleo-cytoplasmic fractionation assays (Figs S1A and 1C). The abundance of N-term in the nuclear fraction thus recapitulated the subcellular distribution of GFP-tagged full-length (F-len) CGGBP1. The non-retention of C-term in the nucleus despite having the same NLS as N-term suggested that DNA-binding or getting incorporated into the chromatin is needed for nuclear concentration and retention as observed for the N-term and F-len. That the N-term and F-len both have the DBD intact strengthened the possibility that direct DNA binding by CGGBP1 through this domain may also be required for its nuclear retention. We also observed no co-immunoprecipitation of N-term and C-term with each other or F-len (Fig S1B) ruling out any direct interference with N-term and C-term nuclear localization patterns by endogenous full-length CGGBP1.

**Figure 1.**
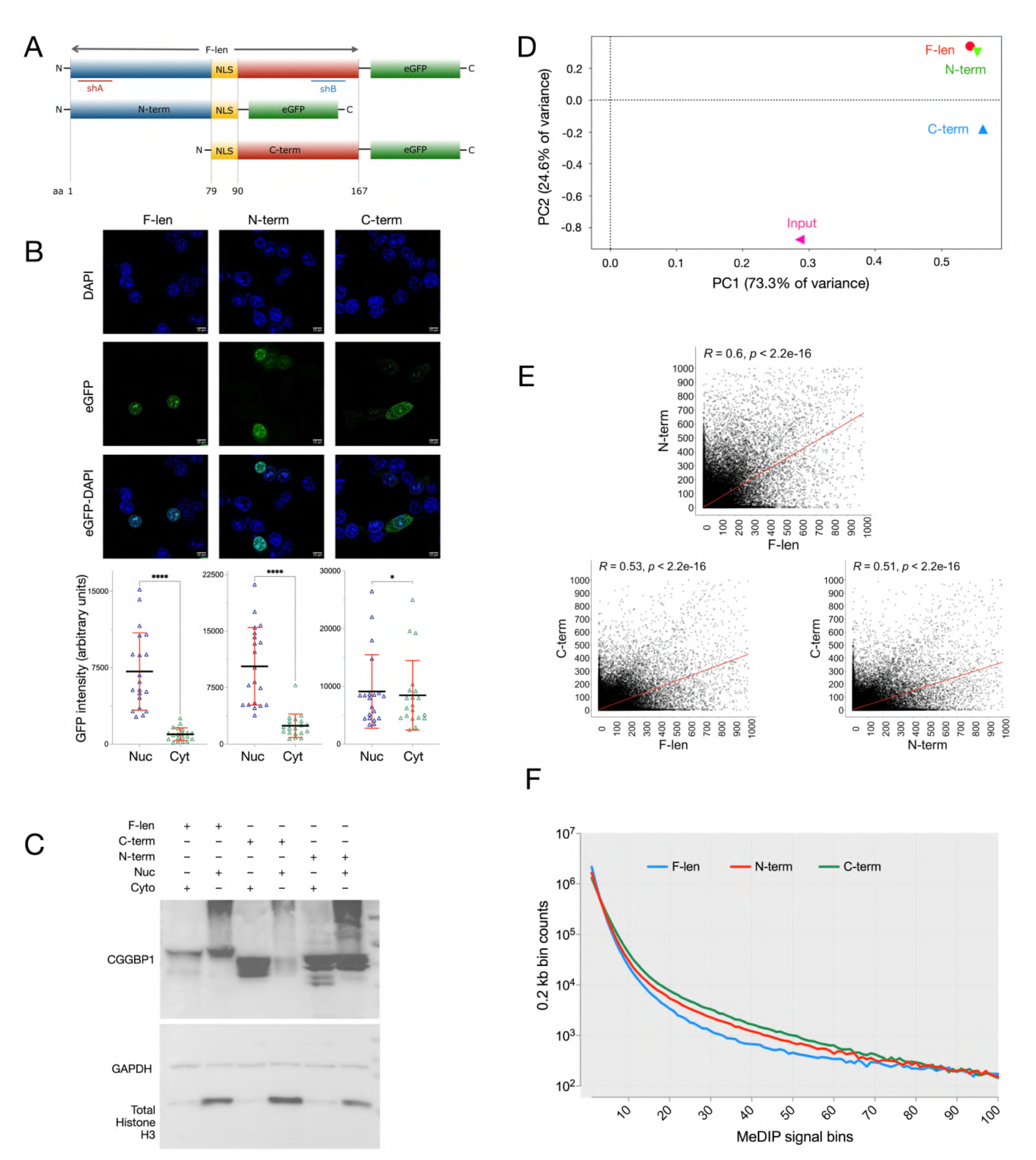
N- or C-terminal truncated forms of CGGBP1 affect global cytosine methylation differently: (A) The Full-length (F-len) CGGBP1 corresponds to UniProt ID Q9UFW8, to which a C-terminal spacer and a eGFP-tag have been appended. The amino acid locations are numbered and a nuclear localization signal-containing region (NLS) is highlighted in yellow. An N-terminal part (N-term; amino acid positions 1-90) was generated with the same C-terminal NLS, spacer and eGFP tag. A C-terminal part (C-term; amino acid locations 79-167 was generated with the same tagging with eGFP except that the NLS was positioned at the N-terminal end. The eGFP tagging was used to establish expression and localization in live as well as fixed cells. Expected molecular weights of the proteins coded by the constructs are mentioned on the right of each construct in kDa. The locations of shRNA used to target only the N-term (shA) or only the C-term (shB) are indicated. (B) Subcellular localisation of eGFP-tagged CGGBP1 shows strong nuclear presence of F-len and N-term. The presence of C-term however is prominently enhanced in the cytoplasm. A quantification of nuclear (nuc) and cytoplasmic (cyt) signals for each sample is shown in the lowermost panel (N-nuc vs N-cyt: n = 20, p value <0.0001; C-nuc vs C-cyt: n = 20, p value = 0.0637; F-nuc vs F-cyt: n = 20, p value < 0.0001). A paired t-test between nuc/cyt ratios shows that the nuc/cyt ratio is significantly decreased in C-term only. (C) Western blot analysis of nuc and cyt fractions shows the enhanced presence of C-term in the cytoplasmic fraction. The nuc and cyt fractions are marked by the detection of total histone H3 and GAPDH respectively. The expected band locations as per the calculated molecular weights of the eGFP-tagged forms are indicated by a yellow rectangle in each lane. Ladder bands are labelled with molecular weights in kilodaltons. (D) PCA plot depicting variance between global cytosine methylation patterns in F-len (red), N-term (green) and C-term (blue) determined by MeDIP-seq. As compared to the Input (pink), the variance in MeDIP has two components: the largest component (PC1 on the X-axis) accounting for MeDIP enrichment which segregates Input from a tight cluster of F-len, N-term and C-term and the second largest component (PC2 on the Y-axis) accounting for differential MeDIP enrichment between the samples. In a combined 97.9% of total variance the N-term and F-len show strong resemblance of cytosine methylation patterns distinct from that of the C-term. (E) A genome-wide scatter of MeDIP signals shows a stronger correlation between F-len and N-term MeDIP signals as compared to C-term MeDIP signals which show similar lower correlations with F-len or N-term. (F) The MeDIP signals are consistently higher in C-term between signal bins 5 and 50. The X-axis bins show the number of MeDIP reads per 0.2kb non-overlapping genomic segments. Y-axis values are counts of these 0.2kb genomic segments. The cytosine methylation distribution shows that F-len maintains low cytosine methylation in the signal range 5-50 MeDIP reads per 0.2kb (blue). Truncation mutations lead to an increase in cytosine methylation in this signal range with the highest increase observed in C-term. The higher area under the curve for C-term is also reflected in the scatter plots (E) in which C-term values show a prominent concentration of data points below 300 on the C-term axis.

To find out the molecular effects of expression of truncated forms of CGGBP1, we assayed global cytosine methylation using MeDIP-seq. In the cells expressing these truncated forms of CGGBP1, the endogenous full-length CGGBP1 was knocked down by using shRNA which could discriminate between N-term and C-term and selectively spare the truncated form being overexpressed (Figs 1A and S1C).

Since CGGBP1 has no known DNA cytosine methyltransferase, methylcytosine deaminase or methylcytosine oxidase activities, the underlying presumption was that CGGBP1 either interferes with or facilitates the activities of methylcytosine regulatory proteins through its interactions with these proteins or the target DNA. We performed MeDIP (details in methods) for N-term, C-term and F-len expressing cells (sequencing statistics in table S1). As compared to F-len, the largest change (an overall increase) in cytosine methylation was observed in C-term overexpressing cells (Figs 1D and S2; peak calling statistics in table S2; Figs 1E and S3). Interestingly, the disruption in cytosine methylation due to the loss of C-term (in N-term expressing cells) was less deviant from F-len as evident from a higher correlation between N-term and F-len MeDIP signals genome-wide (Pearson r = 0.50; Fig 1E) as compared to that between F-len and C-term MeDIP (Pearson r = 0.44; Fig 1E). For a quantitative overview of cytosine methylation changes caused by the loss of function of CGGBP1, we calculated MeDIP signals on 0.2 kb bins genome-wide as described earlier (Patel *et al*., 2020) and compared the distribution of cytosine methylation signal density between the samples. The frequency of a wide range of MeDIP signals (1-100) was consistently high in C-term with comparatively minor differences between N-term and F-len (Fig 1F). These regions with consistently high representation in C-term accounted for >99% of all the MeDIP signal captured. Such differences were not observed for some extremely high methylated regions with MeDIP signals ranging between 100-1000 (Fig S4). These results indicated that a function encoded by the N-term of CGGBP1 mitigates cytosine methylation globally. The N-term of CGGBP1 has only one known functional domain–the DNA-binding domain–raising the possibility that DNA-binding of CGGBP1 keeps cytosine methylation under check through steric hindrance and its loss unleashes cytosine methylation.

These regions with runaway cytosine methylation gain in C-term (MeDIP signal range 1-100 wherein 99% of the MeDIP reads are included; Fig 1F) were analysed for GC-content and G/C-skew, which are two main features associated with cytosine methylation regulation by CGGBP1. High frequency of low-moderate cytosine methylation was concentrated at GC-rich regions. In F-len this range of cytosine methylation was observed at regions with 44-45% GC-content (Fig S5A). The loss of C-term mildly increased the GC-content preference for cytosine methylation (Fig S5A). Conversely, the loss of N-term DBD lowered the GC-content preference slightly (Fig S5A). Similarly, the association between high G/C-skew and low levels of cytosine methylation, as observed in F-len, was remarkably lost in C-term (Fig S5B). Thus, the runaway cytosine methylation increase upon the loss of the DNA-binding domain of CGGBP1 occurs at relatively less GC-rich regions and loses its preference for G/C-skew. A small fraction of regions had extremely high cytosine methylation (MeDIP signal range 100-1000, which includes less than 1% of the MeDIP reads) at which C-term failed to induce the runaway cytosine methylation.

Regions with such an extremely high cytosine methylation by the C-term showed the highest GC-content and least GC-skew, whereas F-len maintained its preference for a relatively lower GC-content and the highest GC-skew (Fig S5 C and D). GC-content and G/C-skew are sequence features associated with regulatory elements. These sequence features present a rich template for cytosine methylation and are commonly exhibited by genome regulatory elements. At the same time such sequences face the evolutionary pressure of preventing the loss of cytosines which can be accelerated by cytosine methylation.

Mitigation of runaway cytosine methylation of cytosines by proteins such as CGGBP1 could aid the retention of cytosines at critical regulatory elements in the genome process such as promoters and transcription start sites (TSSs), enhancer elements and chromatin insulators. Varied patterns of cytosine methylation were observed at conserved regulatory elements (UCSC), enhancers (FANTOM) and TSSs (UCSC) (Fig S6, A-C). It was observed that the lowest cytosine methylation levels at these regions were observed in F-len. At a large subset of these sequences the highest cytosine methylation was observed in C-term. Chromatin insulators and organiser regions are identified by occupancy of CTCF. CTCF occupancy at some regions depends on the levels of CGGBP1 (Patel *et al*., 2019). CGGBP1-regulated CTCF binding sites are also regulated by cytosine methylation (Patel *et al*., 2020). CGGBP1 depletion increases cytosine methylation at canonically known CTCF-binding motifs as well as experimentally determined CTCF binding sites (CTCF-bs) which are devoid of the known motifs (GSE129548). We used the N-term, C-term and F-len MeDIP data to query cytosine methylation changes at previously described CGGBP1-regulated CTCF-bs. A comparison of signals of CTCF occupancy and cytosine methylation in N-term, C-term and F-len showed expectedly that cytosine methylation and CTCF occupancy correlate poorly (Fig S7). For F-len the Pearson coefficient ranged between 0.01 and 0.04 with the highest correlation observed for CTCF-OE (sites where CTCF occupancy depends on CGGBP1). In C-term, although the CTCF-OE sites showed only a slight increase in cytosine methylation (Pearson coefficient 0.06), a noticeable subset of regions distinctly emerged as gaining methylation (Fig S7, regions highlighted in blue insets). Interestingly, this increase in correlation between cytosine methylation in C-term and CTCF binding sites was observed for those CTCF binding sites at which CTCF occupancy is favoured by levels of CGGBP1 (CTCF-OE) but not at CTCF binding sites where CTCF occupancy is hindered by levels of CGGBP1 (CTCF-KD).

We concluded that CGGBP1 keeps cytosine methylation under check and loss of N-terminal DBD leads to a dysregulated increase in cytosine methylation. The restriction of cytosine methylation at flanking regions of TSSs and CTCF-bs raised the possibility that CGGBP1 could regulate transcription factor binding sites near TSSs with a direct effect on the expression of the associated genes.

## 2. Cytosine methylation restriction by CGGBP1 is robust at GC-rich binding sites of amniote-conserved transcription factors

Previous studies have shown that specific transcription factor binding sites undergo cytosine methylation change upon CGGBP1 depletion (Patel *et al*., 2020). So we next surveyed the changes in cytosine methylation brought about by CGGBP1 truncation mutations at known and predicted transcription factor binding sites (JASPAR vertebrates). JASPAR motifs were called using FIMO in MeDIP reads to infer cytosine methylation changes at them caused by CGGBP1 truncation mutations. TFBS abundance, calculated on 0.1 million randomly sampled reads, were consistently higher for 685 TFs in C-term and N-term as compared to F-len (F-len vs N-term paired t test p value <0.0001; F-len vs C-term paired t test p value <0.0001) (Fig S8). These results showed that full-length CGGBP1 maintains a lower level of cytosine methylation at TFBS motifs and any loss of its function due to truncation mutations renders TFBSs more prone to cytosine methylation. This restriction of cytosine methylation by CGGBP1 depends on its N-terminus, which contains the DBD and the expression of the DBD-deficient form of CGGBP1 (C-term) increases cytosine methylation at TFBSs.

GC-rich TFBSs can be regulated by cytosine methylation and thus we asked to what extent the cytosine methylation restriction by CGGBP1 correlates with GC-richness of TFBSs. The TFBSs were ranked according to GC content deviation from 50% (considering an expected value of A/T to G/C ratio of 1) and analysed for cytosine methylation changes on them due to loss of DBD (ratio of MeDIP signals in F-len and C-term). As compared to C-term and N-term both, F-len maintained a lower level of cytosine methylation at GC-rich motifs such that cytosine methylation and motif GC-contents showed inverse correlation (Fig 2 A and B) (F-len vs C-term: Spearman r = -0.5602, n=836, p=<0.0001; F-len vs N-term: Spearman r = -0.6704, n=836, p=<0.0001). The GC-rich TFBSs retained higher cytosine methylation in C-term as compared to F-len. The only motifs at which there was an increase in cytosine methylation by F-len were GC-poor. It follows from these observations that GC-rich motifs are primary targets for cytosine methylation restriction by CGGBP1 through a mechanism that depends on its N-terminal DBD. If cytosine methylation restriction by CGGBP1 at GC-rich TFBSs is functionally relevant then this phenomenon would be concentrated at binding sites of transcription factors with similar evolutionary conservation patterns as that of CGGBP1.

**Figure 2.**
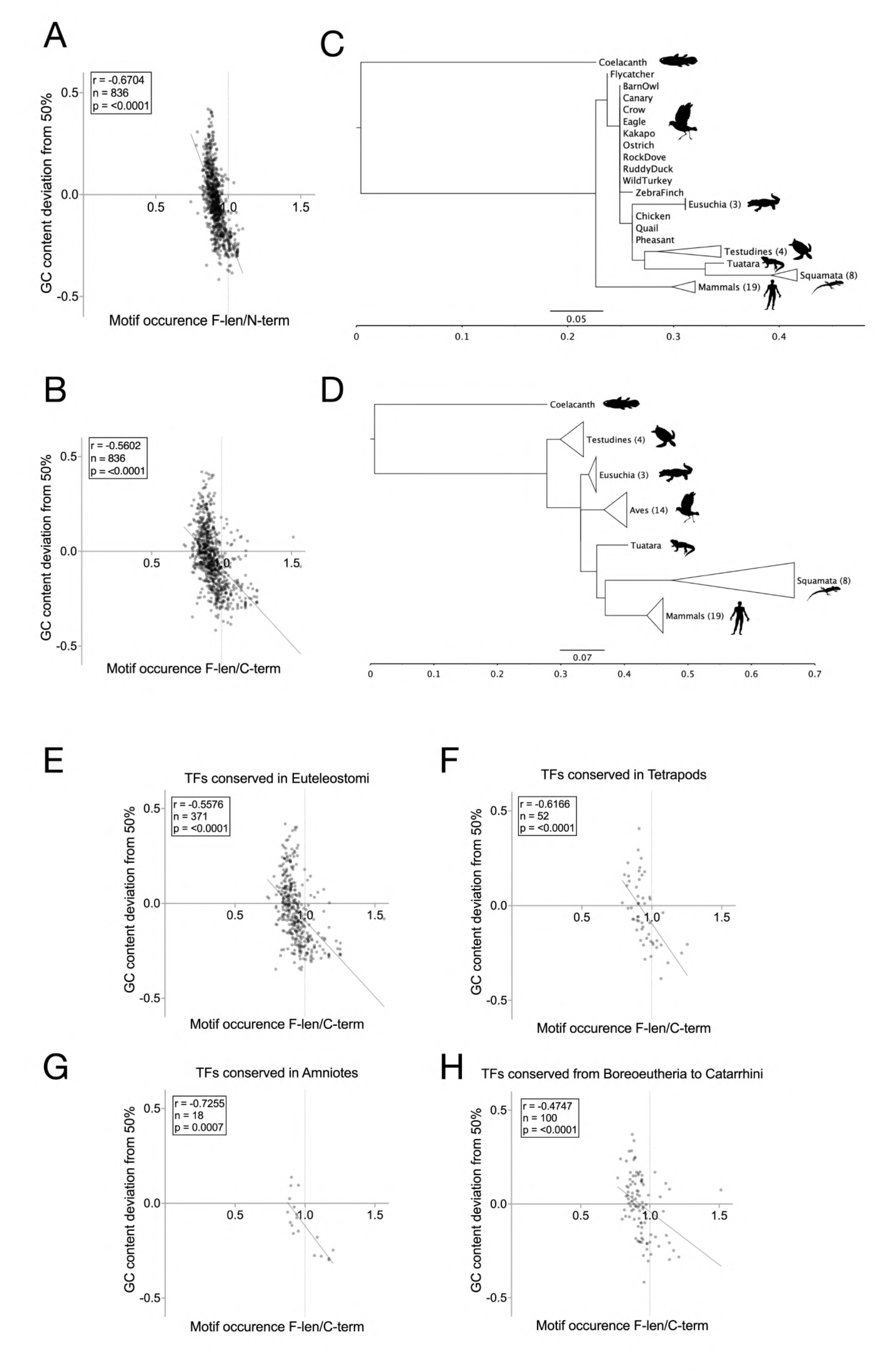
cytosine methylation restriction at TFBSs is an evolutionary property of CGGBP1: (A) JASPAR motifs with low GC content occur similarly in F-len and N-term MeDIP data while most of the motifs with higher GC contents are marginally less abundant in the N-term MeDIP. (B) JASPAR motifs, especially the low GC-content ones, show a higher abundance in C-term MeDIP as compared to F-len. (C and D) Phylogenetic analysis of N-term and C-term show that the C-term (D) is more divergent in vertebrates than the N-term (C). The horizontal axis projects an approximately 1.5x larger distance for all common ancestor nodes. The trees are rooted at "Coelacanth" and representative species for which annotated CGGBPs are available (none available for amphibia) have been used in the analysis. (E-H) The evolutionary conservation status of TFBSs affects their cytosine methylation upon expression of C-term. Of the motifs conserved in euteleostomi (E), tetrapods (F), amniotes (G) or boreoeutheria (H), the highest association of GC-poor motifs with cytosine methylation in C-term is observed for those conserved in amniotes. The correlation coefficients, n counts and p values are marked in top left insets in A, B, and E-H.

We first established the evolutionary conservation status of the N-term and C-term halves of CGGBP1 separately and then analysed cytosine methylation restriction by CGGBP1 at binding sites of TFs conserved in various taxon groups. CGGBP1 has evolved rapidly in tetrapods and become conserved in amniotes (data derived from legacy NCBI HomoloGene in table S4). A comparison of amino acid sequences of CGGBPs from vertebrates and invertebrates suggests multiple hAT domestication events of which the extant amniotes appear to have inherited just a single CGGBP through the common tetrapod ancestor with a loss of CGGBP in the amphibian lineage (Yellan, Yang and Hughes, 2021). Using full-length cDNA and protein amino acid sequences from representative vertebrate taxa we constructed a phylogenetic tree for CGGBP1. It showed that in the mammalian and avian lineages there has been a strong purifying selection in CGGBP1 (Fig S9 A and B). In reptiles, especially squamates, this purifying selection was weaker and amino acid sequence divergence was higher (Fig S9 A and B). To check if the purifying selection operates similarly on N-term and C-term we performed separate sequence analysis and constructed phylogenetic trees for N-term and C-term parts from representative CGGBPs (Table S3). By comparing the N- and C-terminal fragments of CGGBPs from representative taxa we could establish that the N-terminal DNA-binding domain has remained more conserved in the amniotes than the C-terminal part. Amino acid sequence analysis shows that N-term alone recapitulates the evolutionary pattern of F-len CGGBP1 such that the mammalian CGGBP1 derives from the common amniote ancestor (Fig 2C) whereas the C-term, most likely due to sequence divergence, attributable to low purifying selection, relates with the squamate CGGBPs (Fig 2D). We also generated phylogenetic trees using the cDNA sequences of the representative C-term and N-term fragments. Interestingly we found that the N-term, even with its low levels of amino acid divergences, has a larger cDNA sequence divergence as compared to the C-term (Fig S10 A and B). A comparison of the distance values on the X-axis (Fig 2 C and D; Fig S10 A and B) shows that for comparable amounts of evolutionary changes in their cDNA sequence, the C-terminal part has accumulated about 3x more amino acid changes than the N-terminus. For all the major monophyletic groups (mammals, crocodilian lineage and aves) except testudines, the N-terminus containing DBD shows evidence of less evolutionary changes than the C-terminus at amino acid level. This evolutionary difference between N-term and C-term of CGGBP1 appears to be functionally relevant as the structure prediction for human CGGBP1 by AlphaFold also shows a flexible C-terminal part suggesting a lesser evolutionary constraint. Our MeDIP-seq data allowed us to query if cytosine methylation mitigation at TFBSs is one such function. We asked whether cytosine methylation mitigation by the amniote-conserved N-term is functionally relevant. If so, then it is expected that CGGBP1 will target cytosine methylation at binding sites of co-evolving TFs, which are conserved in amniotes.

We categorically analysed cytosine methylation changes caused by loss of DBD at binding sites of TFs conserved in various taxon groups: euteleostomi, tetrapoda, amniota, euarchontoglires, boreoeutheria and catarrhini. JASPAR TFBSs were classified into groups based on their conservation statuses as described in table S4 (data derived from legacy NCBI HomoloGene). Changes in cytosine methylation signals at these TFBSs (from 0.1 million randomly fetched regions positive for MeDIP data) were calculated as cytosine methylation signal ratios F-len/C-term. GC-contents of TFBSs were transformed into deviation from the expected 50% GC content. We observed that the inverse correlation between TFBS GC-content and cytosine methylation signal ratios for F-len/C-term progressively increased from euteleostomi (n = 371, Spearman r = -0.5576, p = <0.0001; Fig 2E) through tetrapoda (n = 52, Spearman r = -0.6166, p = <0.0001; Fig 2F) and peaked in Amniota (n = 18, Spearman r = -0.7255, p = 0.0007; Fig 2G). Further on, in euarchontoglires, boreoeutheria and catarrhini no such strong relationship was observed between TFBS GC content and cytosine methylation changes caused by loss of N-term (n = 100, Spearman r = -0.4747, p = <0.0001; Fig 2H).

Based on these results we concluded that GC-rich motifs are maintained in low cytosine methylation states by F-len CGGBP1 through a mechanism that requires its N-term. Low TFBS cytosine methylation allows facultative binding of cognate TFs as cytosine methylation often constitutively abrogates TF-binding. We next studied if TFBS cytosine methylation restriction by CGGBP1 is important functionally for gene expression regulation by CGGBP1.

## 3. C-repressed gene promoters have GC-rich TFBSs embedded in GC-poor promoters with signs of restriction of cytosine methylation

Global gene expression profiling of N-term, C-term and F-len expressing cells were performed using Agilent microarrays. All samples were assayed in triplicates and after background subtraction the signal values were subjected to intra-sample group quantile normalisation (Zhao, Wong and Goh, 2020) (Fig S11A). Significantly differentially expressed genes were extracted by performing T-tests on the quantile-normalised signal values of each probeset (n=3) for the following comparisons: N-term/F-len, C-term/F-len, N-term/C-term (Fig S11B). The array-wide differential expression is shown as M values of mean signals plotted against -log10 p-values (Fig S12). Without applying any M value cutoff the differentially expressed genes were fetched solely based on a p value threshold of <0.01 (differentially expressed genes list in table S5 ).

We could observe that a distinct set of probes exhibited a higher expression in N-term as compared to C-term or F-len (Fig S12). Interestingly, an unexpectedly large fraction of these genes with high expression in N-term also showed a lower expression in C-term as compared to F-len (Fig 3A). However, the M value deviation from zero was higher for increased expression in N-term than that for decreased expression in C-term. Of the 5706 genes showing significantly higher expression in N-term/F-len, 3309 genes showed higher expression in N-tem/F-len as well as N-term/C-term (Fig 3B; coordinates of 2223 genes available from GENCODE (v43) out of 3309 in table S6). These results showed that CGGBP1 represses a set of genes and this repression depends on the C-term of CGGBP1. The loss of the C-terminal repressive part of CGGBP1 leads to gene de-repression in N-term.

**Figure 3.**
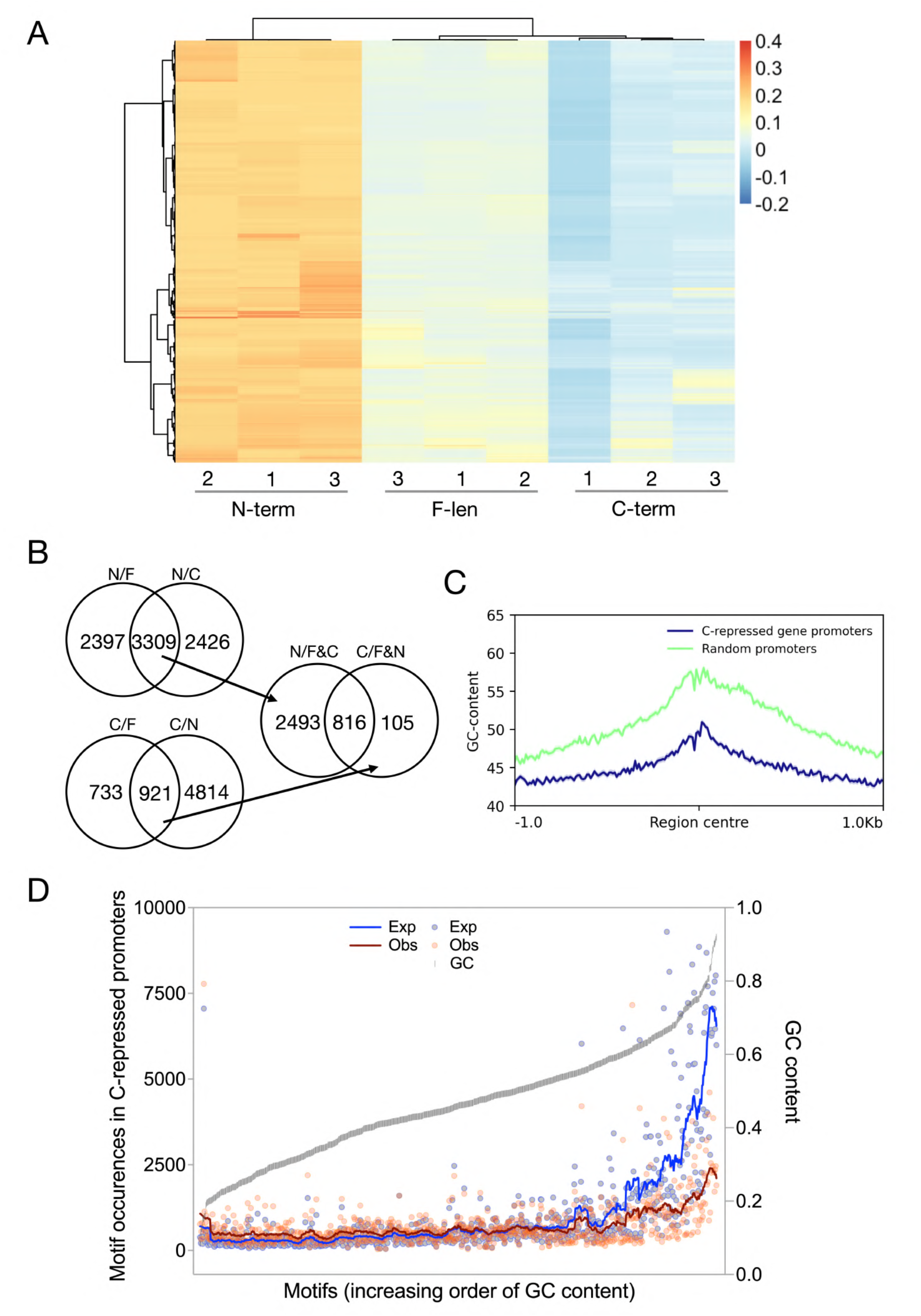
Truncated forms of CGGBP1 target genes with GC-poor promoters: (A) An unsupervised clustering of expression values from three independent replicate experiments for genes significantly differentially expressed between N-term, C-term and F-len classifies the three samples as expected. Although there are variations between the replicates, the arbitrary expression values show a general repression of expression by C-term and a derepression in N-term as compared to F-len. (B) 3309 genes are commonly upregulated by N-term when compared to FL (N/F) and C-term (N/C); 921 genes are commonly downregulated by C-term when compared to FL (C/F) and N-term (C/N); When these 3309 and 921 genes are compared, a considerably large number of 816 genes are commonly upregulated by N-term (N/F&C) and downregulated by C-term (C/F&N). (C) The C-repressed genes have a GC-poor promoter profile as compared to the reporters assayed on the microarrays. (D) GC-rich motif abundance in C-repressed promoters (red) is lower than expected (blue). However, even if the C-repressed promoters are GC-poor, the most abundant motifs in them are GC-rich.

Intriguingly, the genes deregulated by expression of truncated forms of CGGBP1 did not show any functional category enrichment (data not shown). We focussed on the genes de-repressed by N-term (These genes will henceforth be referred to as ‘C-repressed genes’). For further analysis we focussed on 2223 genes out of the 3309 genes for which reliable GENCODE annotations were available. The 1kb proximal promoter regions of these genes were poor in Alu and GC contents (Figs 3C and S13) whereas high Alu and GC content are two main features of DNA binding to CGGBP1.

The C-term, bearing similarity to the Hermes transposase DBD, seems capable of interacting with different proteins. If the absence of C-term interferes with the transcription regulation by CGGBP1 at specific transcription factor binding motifs, then the proximal promoter regions of the N-term de-repressed genes would show the presence of such TFBSs. We pursued this possibility and tested it rigorously from three different perspectives: (i) What are these TFBSs and do they have any sequence properties known to be regulated by CGGBP1? (ii) Is there an evolutionary justification for a gene regulatory cooperation between these TFBSs and CGGBP1; the latter being conserved in amniotes with an evolutionarily divergent C-terminal? (iii) Do the promoter sequences containing these TFBSs display sequence features of cytosine methylation restriction by CGGBP1 as observed in MeDIP experiments?

Interactions between human CGGBP1 and specific transcription factors could explain why a set of genes are repressed by C-term. The binding sites of such transcription factors could be enriched in the proximal promoter regions of the C-repressed genes. A *de novo* motif search in the 1 kb upstream sequences of the C-repressed genes returned some GC-poor motifs (Data not shown). The promoter regions of C-repressed genes are GC-poor (Fig 3C), and hence a preponderance of some GC-poor motifs is expected. These GC-poor motifs did not bear similarities with known TFBSs and we could not ascribe the regulation of C-repressed promoters to these TFs confidently.

To circumvent the masking of TFBSs by GC-poor motifs as a consequence of GC-poorness of C-repressed promoters, we performed a search for the pan-vertebrate TFBS set from JASPAR with a p-value threshold of 1e-4. This targeted search for known TFBSs revealed that despite their GC-poorness, the most abundant TFBSs in C-repressed promoters were those with high GC-contents. Such a presence of GC-rich TFBSs in GC-poor sequences suggested that the occurrence of TFBSs were not merely a consequence of the base compositions of the C-repressed promoters. To resolve this further we performed a comparison of the TFBS abundance and GC-contents of the C-repressed promoters with those of a randomly-derived null set of promoters. The GC-rich TFBS abundance in C-repressed promoters was lower than expected when compared using a random set of promoters which were n-matched but not controlled for GC-content (and hence having higher GC-content) (Fig 3D). These findings pointed to the possibility that in the C-repressed promoters the GC-rich TFBSs have been selectively maintained even when the promoter sequences are GC-poor. We concluded that some known GC-rich TFBSs, embedded in GC-poor proximal proximal promoters, are associated with gene repression by C-term. To understand how cytosine methylation and its restriction by CGGBP1 impacts these TFBSs, we investigated the abundance of these TFBSs across different tetrapod taxa in orthologous promoter sequences.

## 4. Localised low C-T transition in C-repressed gene promoters indicate TFBS retention due to low cytosine methylation

If the GC-rich TFBSs, which CGGBP1 appeared to protect from cytosine methylation, are indeed important for gene repression by CGGBP1 C-term, then they would have been selectively retained against an overall loss of GC in these regions. Such a selective retention of motifs can be expected to show some evidence for co-evolution with CGGBP1 C-term.

To test these possibilities we analysed the JASPAR-wide TFBS abundance in the promoters of orthologs of C-repressed genes across different tetrapod taxa. The 1kb upstream sequences from transcription start sites for the orthologs of C-repressed genes were analysed from 36 different species for which reliable orthology information was available. For the number of C-repressed orthologs identified (observed) in each species, an equal number of randomly sampled TSS upstream 1 kb sequences were used as null-set (expected). The observed-expected motif occurrence count differentials across these different taxa would represent if the TFBSs have become progressively enriched or lost in the course of evolution at these promoters. This analysis also allowed determining parallels between evolutionary changes in CGGBP1 C-term and the TFBS concentrations in the target gene promoters. We used Coelacanth orthologs as a starting point since the extant CGGBP1 in higher vertebrates is understood to have evolved from one of the many CGGBPs present in Coelacanthini. Most TFBSs which were highly abundant with high obs-exp differentials did not show any pattern in occurrence across the 36 species (Fig 4A). Interestingly, a cluster of 45 TFBSs (Table S7) showed consistently low obs-exp differentials in mammals only (Fig 4B). In aves and reptiles there were notable exceptions and some avian species had extremely large obs-exp differentials for all the 45 TFBSs. The highest obs-exp differentials were observed for non-amniotes used in the analysis. We could confirm that these 45 motifs have high GC-content (Fig S14), show lower than expected abundance in C-repressed promoters and yet are amongst the most highly occurring motifs in these regions (Fig 3D). These data suggested that either a set of GC-rich C-repressed TFBSs, which were highly abundant in pre-amniote common ancestors, have been increasingly lost during the evolution of mammals, or that the non-amniotes have progressively concentrated these motifs throughout their evolution with some lineage-specific differences. The low obs values for these TFBSs combined with low GC content of the promoter sequences indicated a loss of C/G, which is known to be accelerated by cytosine methylation. The restriction of cytosine methylation by CGGBP1 seems to retain these GC-rich motifs against a wave of GC-loss in these promoter regions. We examined if these 45 TFBSs in the human C-repressed promoters actually exhibit signatures of cytosine methylation associated GC-loss.

**Figure 4.**
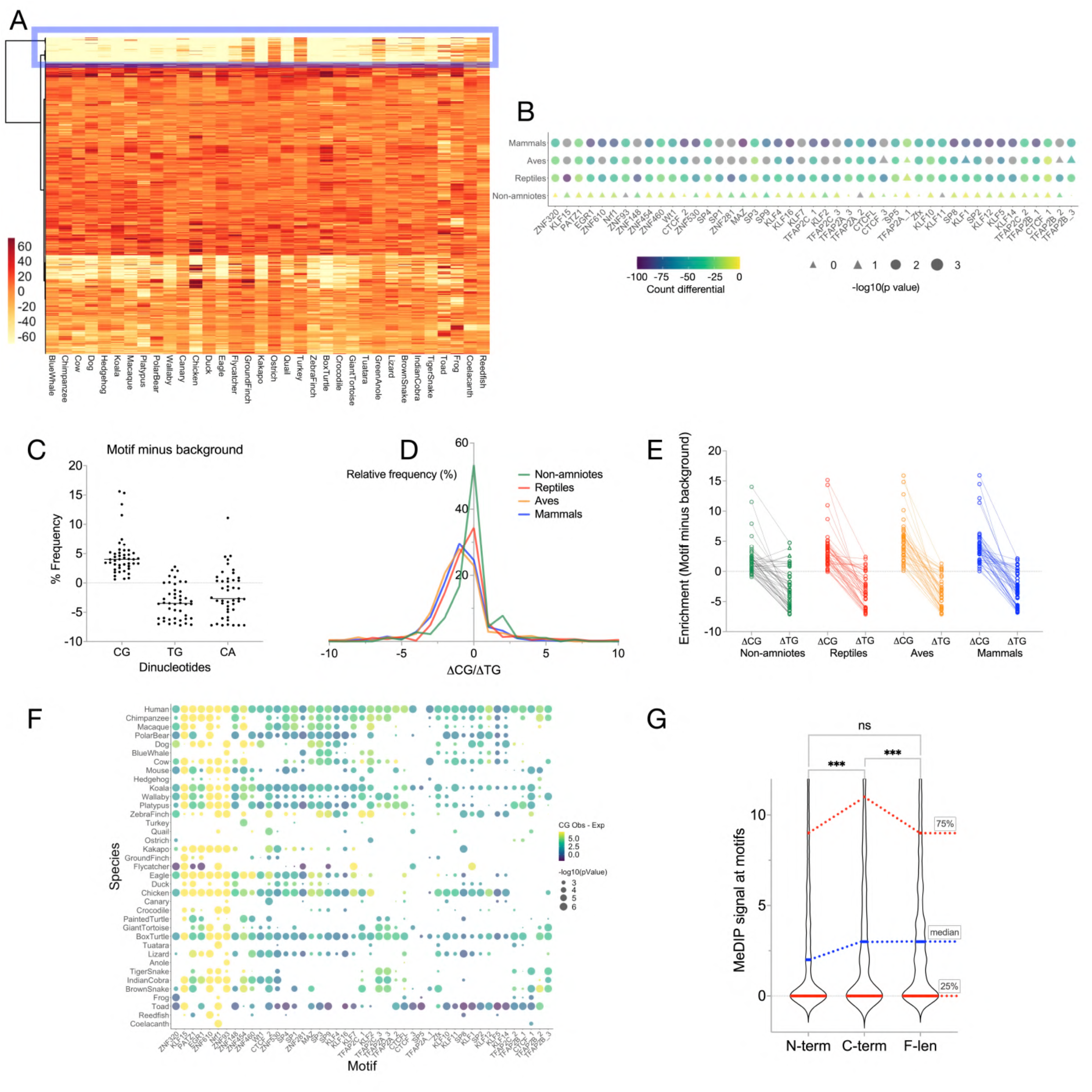
GC-rich motifs in GC-poor promoters of C-repressed genes show signs of cytosine methylation restriction by CGGBP1: (A) JASPAR-wide motif abundance in 1kb promoters of orthologs of C-repressed genes. The species from which orthologous sequences were derived are indicated below the heatmap. Clustering of motifs (rows) yields a set of 45 TFBSs (highly GC-rich, Fig S14) depleted in mammalian orthologs, divergently present in non-mammalian amniotes and present highly in non-amniotic representatives. (B) An observed-expected comparison of these 45 motifs using a randomly derived null-set from each species shows significant differences in all amniotes (mammals, aves and reptiles) with no significance observed in non-amniotes (pooled information from Toad, Frog, Reedfish and Coelacanth). The observed-expected differentials show lowest and consistently significant values in mammals for all the 45 motifs. The highest differentials were observed in reptiles. In aves the differential values were intermediary with insignificant differences for some TFBSs. The insignificant differences are depicted as triangles, significant differences are depicted as circles, circle sizes denote significance (scale at the bottom right) and observed-expected differentials are indicated by the colour scale (bottom left). All p values are derived from paired Mann-Whitney test. (C) The 45 GC-rich motifs depleted in mammals yet most abundant in C-repressed GC-poor promoters show dinucleotide content in human which indicates a protection of these motifs against cytosine methylation. The count differentials of CG/CG and its derivatives, arising through a C-T transition, TG/CA in the motifs and in adjacent non-motif regions show that the motifs are CG/CG-rich and TG/CA poor as compared to local background. (D) At different sets of orthologous promoters from different taxon groups the CG-TG count differentials show that mammals have the lowest CG-TG tradeoff between motifs and background followed by aves and reptiles. The non-amniotes show the highest CG-TG tradeoffs. (E) The CG and TG values underlying the distribution plots in D are depicted in paired CG-TG differentials for each of the 45 TFBSs (all CG and CT differentials are significantly different with p<0.01). (F) A bubble plot matrix showing the CG differentials derived from motifs and local background for each of the 45 TFBSs (horizontal axis) and species (vertical axis). It is appreciable that mammals form the group with the most consistent positive CG differentials for almost all the motifs; some notable exceptions include CTCF (highlighted) which are GC-rich but inherently CG-poor. (G) MeDIP signals at these 45 motifs were compared for differences between N-term, C-term and F-len. As compared to F-len, the N-term showed a net insignificant but noticeable reduction in cytosine methylation at motifs with low overall cytosine methylation (between median and 25% of the MeDIP signals). However, loss of N-terminal DBD in C-term led to an increase in cytosine methylation at these motifs such that MeDIP signals in C-term were significantly higher than those in N-term as well as C-term (N-term vs F-len: n = 1468, p value = 0.3692; C-term vs F-len: n = 1468, p value = 0.0081; N-term vs C-term: n = 1468, p value = 0.0002).

Cytosine methylation facilitates C-T transitions due to spontaneous deamination of methylcytosine that escapes base excision repair (BER) at a background rate. A difference in C-T transition rates between the GC-rich motifs and the neighbouring 1 kb promoter regions could explain how these motifs have been retained. We hypothesised that CGGBP1-binding to these motifs could exert two independently manifested but linked effects: (i) repressing the associated gene and (ii) minimising cytosine methylation and thus attenuating C-T transitions.

CGGBP1 is known to antagonise cytosine methylation at CpG dinucleotide context and this is calculable as inverse changes in CG and TG dinucleotide frequencies upon CGGBP1 knockdown. We first analysed the base composition of the motifs and associated 1 kb regions for signs of resistance to C-T transitions by comparing the frequency of CG and TG(+CA) dinucleotides in either the entire 1 kb upstream regions (background) or specifically in the 45 motifs for all the 2223 C-repressed genes. This calculation was performed for the human genes and their orthologs in 36 different species. Pooled data for each species showed a strong inverse correlation between the CG and TG(+CA) dinucleotide frequencies calculated by subtracting the background dinucleotide frequencies from those of the motifs (Fig 4C). It became clear that in the motifs there is a positive retention of CG and a deficit of TG as compared to the respective backgrounds. However, given the differences in base compositions between different species and the different motifs, the significance of these findings depended on a segregated comparison of each species-motif combination against a null set of sequences and motifs called in the genome of the same species. FIMO searches for the same 45 motifs were performed on a randomly drawn set of 2223 gene-upstream sequences or corresponding orthologs (used for calculating expected values). The following four parameters were used for a Fisher exact test for each species-motif combination: CpG% observed in motifs, CpG% observed in the background, CpG% expected in motifs and CpG% expected in background. We observed that the positive CG contents in motifs as compared to background were significantly different from those identified in the null set of sequences and motifs with the highest consistency of difference observed amongst 12 mammalian species including humans analysed (Fig 4, D-F). The reptilian and avian representatives showed a high species-specific variation. These results showed that the C-repressed genes are associated with motifs which are GC-rich in a way that CpG and TG dinucleotide frequencies suggest a low rate of C-T transition in humans and mammalian orthologs of the C-repressed genes. It is possible that the inconsistency of this mitigation of C-T transition in non-mammalian orthologs is due to a combination of two major components: the differences in various evolutionary forms of CGGBP1 and the genome compositional differences between different taxa.

In the flanks of the TSSs of C-repressed genes we could see that the cytosine methylation in C-term was consistent with lower variation along the promoter. On the other hand cytosine methylation patterns in N-term and F-len showed higher variations within the promoter (data not shown). The differential methylation of TFBSs in N-term, C-term and F-len could explain the cytosine methylation variability within promoters. If the GC-rich motifs exhibiting low C-T transition rates were indeed retained by cytosine methylation mitigation, we expected to observe cytosine methylation signal differences on and around these motifs supporting this proposition. To test this we compared cytosine methylation signals from N-term, C-term and F-len in the 1 kb promoters of the C-repressed genes distinctly at these 45 motifs. Overall, this comparison showed that cytosine methylation by N-term and F-len was indeed significantly low at the motifs (Fig 4G). These findings supported the possibility that cytosine methylation mitigation by CGGBP1 maintains low C-T transition rates thereby retaining GC-rich TFBSs. The C-term alone was not as potent in restricting cytosine methylation at motifs (Fig 4G). Such differences between N-term and C-term indicated that the targeted cytosine methylation mitigation distinctly at the TFBSs is dependent on the CGGBP1 DBD.

## 5. TFBSs under cytosine methylation restriction by CGGBP1 show CGGBP1 occupancy

The runaway cytosine methylation due to a loss of N-term suggests that the protection of GC-rich motifs from cytosine methylation could be due to binding of CGGBP1. A series of reports suggest an association between mitigation of cytosine methylation and genomic occupancy of CGGBP1 (Deissler *et al*., 1997; Müller-Hartmann *et al*., 2000; Agarwal *et al*., 2015, 2016). So next we asked if CGGBP1 DBD occupies these motifs and if these 45 motifs exhibit sequence properties previously reported to be regulated by CGGBP1.

We randomly sampled 1 million reads and analysed them for the presence of these 45 motifs in CGGBP1-ChIP-seq (GSE187851) and the corresponding input. All the 45 motifs exhibited a significant enrichment in CGGBP1 ChIP-seq over the input (Fig 5A). The C-repressed set of genes were identified by overexpression of the N-term and hence we argued that if these motifs are indeed regulated by occupancy of CGGBP1 then these motifs will show an association in N-term ChIP-seq as well. Through a ChIP-seq experiment (sequencing statistics in table S9 ) we found that the characteristic preference of CGGBP1 for binding to GC-rich DNA was retained by the N-term. All the 45 motifs were significantly over-represented in N-term ChIP-seq over the corresponding input in randomly sampled 1 million reads (Fig 5B). Interestingly, this similarity between CGGBP1 full-length ChIP-seq and N-term ChIP-seq data was retained even when there were other evident differences. However, apparently the occupancy of CGGBP1 at these motifs was enhanced upon the loss of C-term. Notably, the N-term occupancy did not show any strong preference for Alu-SINEs as reported earlier. It was also devoid of any significantly scoring peaks and binding was detected genome-wide with no clear sequence preference. Collectively, these results reinforced that CGGBP1 mitigates cytosine methylation, preferentially at TFBSs through a mechanism that depends on its N-terminal DBD as well as its interaction with DNA.

**Figure 5.**
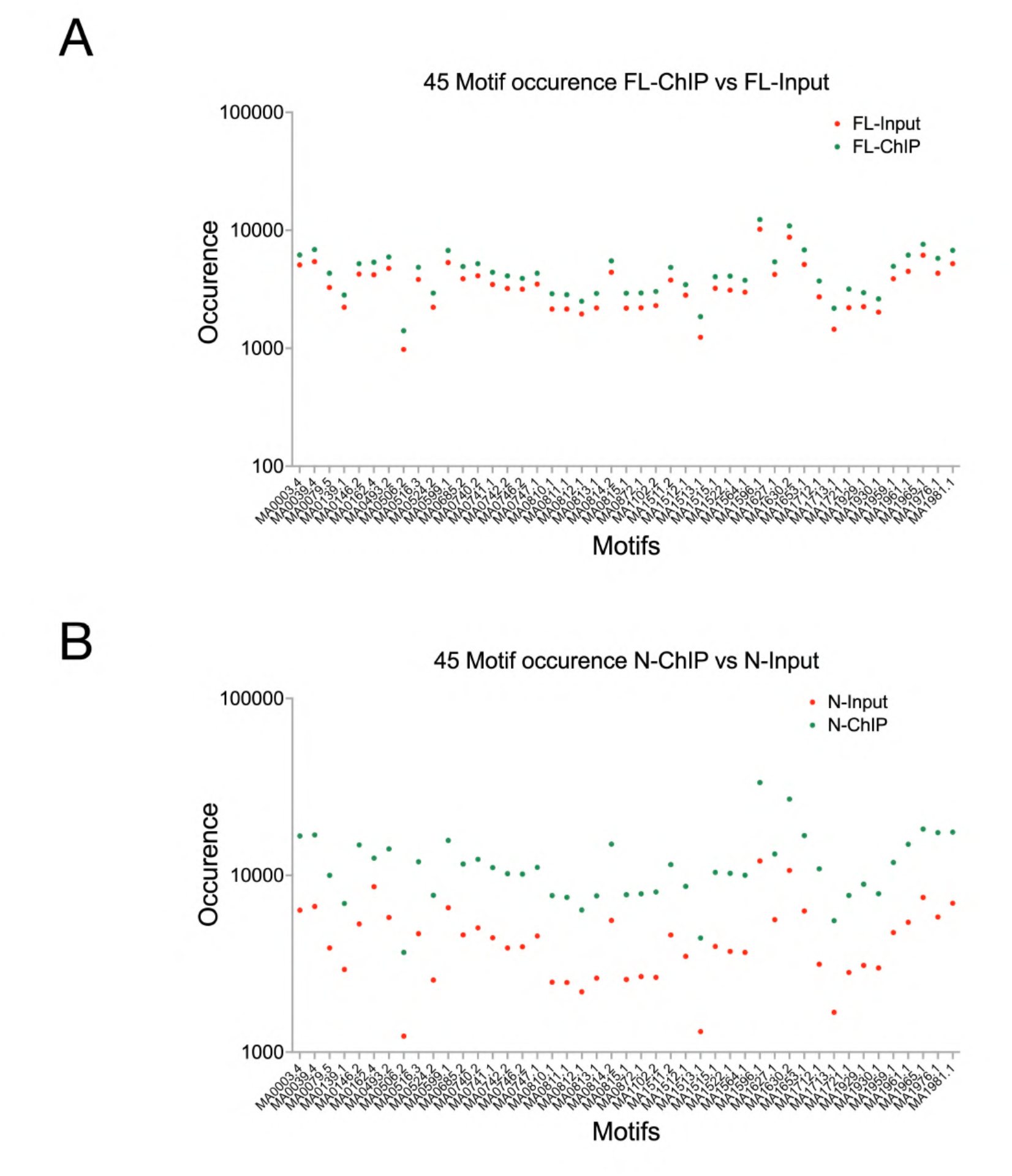
CGGBP1 occupancy determines methylation restriction at target GC-rich TFBSs. A. The abundance of the 45 motifs at which CGGBP1 restricts cytosine methylation were fetched from ChIP-seq data for full-length CGGBP1 (GSE187851) as well as the corresponding input. The occurrences were normalised to sequencing depths of input (red) and ChIP (green). The comparisons clearly showed a consistent enrichment of the motifs in CGGBP1 ChIP-seq (all motifs combined test p value <0.0001). B. To test if the N-term is capable of binding to these motifs, we compared the abundance of these motifs in input (red) and ChIP (green) for N-term. The comparisons show a strong enrichment in N-term ChIP-seq (all motifs combined test p value <0.0001) showing that N-term is capable of and sufficient for occupancy at these motifs resulting in mitigation of cytosine methylation through steric hindrance.

## Discussion

CGGBP1 has been hitherto understood to be a GC-rich DNA-binding protein with gene repressive functions (Singh and Westermark, 2015). Some of the pioneering works on CGGBP1 involved in vitro DNA-protein interactions between GFP-tagged CGGBP1 expressed in HEK293T cells and oligonucleotides of varying GC and methylcytosine contents (Deissler *et al*., 1997; Müller-Hartmann *et al*., 2000). Interactions of CGGBP1 with unmethylated DNA, its preference for ribosomal repeats (rich in CGG tandem repeat sequences) *in situ* and its ability to repress FMR1 gene, likely through regulation of cytosine methylation in its 5’-UTR, framed our initial understanding about this protein (Deissler *et al*., 1997). Increasingly, more recent works have added complexity to the functions of CGGBP1. Mitigation of cytosine methylation (Agarwal *et al*., 2015), suppression of endogenous DNA damage (Singh *et al*., 2014; Datta *et al*., 2022), regulation of global transcription and stress response through regulation of Alu repeat transcription (Agarwal *et al*., 2016) are some of the functions which have put CGGBP1 at the intersection of many seemingly disparate sets of functions, most of which are very well conserved. More recent works have shed light on molecular mechanisms underlying the multiple functions of CGGBP1. It seems that CGGBP1 is required for maintaining a genome-wide cytosine methylation pattern (Patel *et al*., 2018). CTCF, a master regulator of the epigenome with key functions in chromatin barrier function, insulator activity and transcription is one of some transcription factors which are regulated by CGGBP1 (Patel *et al*., 2019, 2020). We had recently described two types of CTCF-binding sites; motif-free repeats and repeat-free motifs. Depletion of CGGBP1 leads to a gain of cytosine methylation at motif-free repeats and a loss of cytosine methylation at repeat-free motifs (Patel *et al*., 2020). A combined interpretation of the work on CGGBP1, CTCF and cytosine methylation suggests that regulation of cytosine methylation is the cornerstone of CGGBP1 function. Its role in histone modification landscape and CTCF occupancy genome-wide can be explained through its regulation of cytosine methylation. Anecdotal evidence also suggests that these functions of CGGBP1 depend on its interaction with the DNA. Inter-strand G/C-skew is an additional feature of the regions at which cytosine methylation is regulated by CGGBP1 (Patel *et al*., 2018). Most recently, we have reported through in vitro as well as genome-wide studies that a physical interaction between CGGBP1 and target DNA is required to prevent formation of G-quadruplexes (Datta *et al*., 2024). These regions where CGGBP1 inhibits G-quadruplex formation exhibit features like G/C-skew and presence of CTCF-binding sites. Given that interactions between CGGBP1 and target DNA seem important for its function, it becomes imperative to study the importance of CGGBP1 DNA-binding domain in its function. Studies using truncated mutant forms of CGGBP1, including deletion mutants lacking a DNA-binding domain, have been done earlier but only in vitro and with limited inferences.

The domain structures of CGGBP1 only partly explain its functions as there is only one defined domain present in CGGBP1; a Zn-finger BED DNA binding domain. This study describes and delineates the roles of different parts of CGGBP1 in regulation of cytosine methylation and gene expression. Our findings suggest that cytosine methylation regulation at TFBSs is one of the major factors behind gene expression regulation by CGGBP1. The results presented in this study highlight that there is a remarkable association between CGGBP1 occupancy and a negative regulation of cytosine methylation. We show that the only known domain in CGGBP1, that is the N-terminal DNA-binding domain, is required for binding of CGGBP1 to cognate sites on DNA, prominently some GC-rich TFBSs, for this negative regulation of cytosine methylation. We discuss how our findings link the various functions of CGGBP1 to its DNA-binding property.

We have applied deletion mutants of CGGBP1 to describe the mechanisms of action of CGGBP1 by using two major read-outs: cytosine methylation and gene expression patterns genome-wide. Through a comprehensive analysis of our experimental data we show that (i) CGGBP1 is majorly a transcription repressor, (ii) the transcription repression function of CGGBP1 rests in its C-terminal part, (iii) truncation mutations disrupt cytosine methylation patterning in the genome such that some GC-rich TFBSs, including CTCF-binding motifs, become methylated in the flanks of target gene TSSs when the DBD is deleted.

Our analysis of CTCF-binding site cytosine methylation changes by CGGBP1 truncation mutants establish that all CTCF binding sites are not affected by the loss of C- or N-terminal domain of CGGBP1. It is only the CTCF-binding sites which are regulated by CGGBP1 that undergo cytosine methylation change by expression of N- or C-terminal CGGBP1. The increase in cytosine methylation at CGGBP1-regulated CTCF-binding sites (motif-rich and repeat-poor) is observed only when CGGBP1 is expressed without its DNA-binding domain. It suggests that a proper localization to the target sites and binding to them is required for restricting cytosine methylation levels at CTCF-binding sites. As a consequence the target promoters avoid constitutive silencing by cytosine methylation and remain facultatively repressed by CGGBP1 through its C-terminus. But when the C-terminus is lost, the binding is retained but the repression is lost.

Cytosine methylation mitigation effects of CGGBP1 CTCF-binding sites could have broad epigenetic effects as CTCF-binding to DNA is cytosine methylation sensitive. Similarly, cytosine methylation regulation by CGGBP1 affects DNA-binding sites of more factors which is likely to affect transcription factor binding on DNA and consequently TFBS evolution (The FANTOM consortium, no date; Kaluscha *et al*., 2022; Krieger *et al*., 2022; Grau, Schmidt and Schulz, 2023). Three of our findings lend weight to this: the GC-rich motifs occur at high frequency in otherwise GC-poor cis-promoter regions of target genes; the signature dinucleotide composition of low CG to TG transition rate is observed only at the GC-rich motifs and not the rest of the promoter region. Also, these motifs show specific enrichment in ChIPseq data for N-term as well as full-length CGGBP1. The JASPAR (vertebrate)-wide search for motifs showed that the most GC-rich TFBSs were enriched in the promoter regions of C-repressed genes suggesting that the GC-rich motifs have been retained in these regions despite a loss of GC-content, likely due to C-T transition mutation hastened by cytosine methylation. Since cytosine methylation in the non-CpG context is unpredictable by such a sequence composition analysis, we have used only CpG as a typical sequence (Agarwal *et al*., 2015). However, many of the target motifs protected by CGGBP1 seem to be GC-rich without being CpG rich. Arguably, a protection against C-T transition and retention of these motifs can be achieved by the binding of these motifs to their cognate transcription factors. Although unverifiable, some evidence in our analysis indicates that this is not the case and much of this protection from cytosine methylation actually depends on binding of CGGBP1 to these motifs. For instance, the majority of binding factors for these motifs show no evidence of co-evolution with CGGBP1 insofar as conservation in amniotes is concerned. Nevertheless, the binding sites of a small set of amniote-conserved transcription factors show an inverse relationship between CGGBP1-favoured cytosine methylation and GC-content. It appears that these GC-rich motifs and their cytosine methylation prevention is consequential for amniotes and as such, the N-terminal part of CGGBP1 (containing its DBD) is conserved in the amniotes.

Our analysis of orthologous promoter sequences and their CpG dinucleotide content analysis shows that this protection against cytosine methylation associated C-T transition is a process well conserved in mammals and it parallels the evolution of C terminus of CGGBP1. Such a mechanism can drive evolution by generating and retaining TFBSs on DNA in regulatory regions thereby affecting the epigenome, gene expression and other functions of the DNA that are directly sterically regulated by protein binding, including G quadruplex formation. These results also explain why the deregulated genes are not identifiable as rich in any specific functional categories. It seems that CGGBP1 is primarily a regulator of cytosine methylation and through it contributes to TFBS evolution in the genome. The deregulation of genes is simply a secondary outcome of the association of TSSs with a region where cytosine methylation at TSSs is affected by changes in CGGBP1 structure and function. Previous studies have shown that when CGGBP1 depletion is combined with additional conditions such as serum stimulation or starvation of cells in culture, specific functional categories of genes do emerge as affected by CGGBP1 (Agarwal *et al*., 2015, 2016). Interestingly these CGGBP1-target genes consist of several regulators of cytosine methylation including DNA methyltransferases and TET family of oxidases involved in BER pathway. Thus, it seems that CGGBP1 has evolved to interface with cytosine methylation regulation with cellular growth stimulation and response to growth factors.

The C-terminal part of CGGBP1 however has a rather deviant evolutionary trajectory in the reptilian and avian lineages suggesting that the selection pressures for changes in the N- and C-terminal parts of CGGBP1 have been different. The N-term directs CGGBP1 to target sequences, such as CTCF-binding sites and other GC-rich motifs whereas the C-term affects the outcome of the interaction between CGGBP1 and the motifs. The C-term has a less rigid structure, can interact with a variety of proteins and hence brings about a diversity in cytosine methylation regulation by CGGBP1.

Human CGGBP1 is known to regulate GC-rich DNA with a preference for G/C-skew. The mammalian orthologs of human C-repressed gene promoters showed the most consistent high G/C-skew followed by aves. This consistency was lost in the reptilian, amphibian and coelacanth orthologs (data not shown). Thus, the evolution of tetrapods, as viewed by emergent differences in GC-content distributions in the genomes,could be subject to further diversification by site-directed cytosine methylation. The functional diversity that can be measured through just the patterns of GC content distribution in the genomes could be explained by an additional dimension of cytosine methylation. The status of regulators of cytosine methylation becomes consequential in this regard. Since DNA methyltransferases are sequence-non-specific as well as well conserved in all vertebrates, it is evident that the epigenomic diversity rests on subtle differences in the deployment of methyltransferases, which could be achieved by proteins such as CGGBP1. Similarly, the machinery of BER which would undo the C-U or meC-T mutations are well conserved and their evolutionary status does not explain how the epigenomic diversity is derived. In representative reptiles, such as Anolis, the GC content is uniformly distributed and it is purported that this is a shared property inherited from the common ancestor of all amniotes (Organ, Moreno and Edwards, 2008; Pasquesi *et al*., 2018; Beauclair *et al*., 2019). However, the rapid diversification of GC content distribution in mammalian and some avian genomes could be affected by sequence motif directed cytosine methylation protection by proteins like CGGBP1.

The findings presented in this work show that CGGBP1 is a negative regulator of cytosine methylation and that this function of human CGGBP1 partly depends on its interaction with DNA. Our findings raise interesting possibilities about how evolutionary changes in CGGBP1 could have contributed to evolution of the cytosine methylome and the epigenome in amniotes. The role of DNA-protein interactions in restriction of cytosine methylation makes case for a two-way mechanism; changes in CGGBP1 affecting the cytosine methylome and changes in DNA sequence affecting interaction with CGGBP1 and thereby cytosine methylation. The effect of this mechanism in maintenance of TFBSs suggests that changes in DNA sequence (including TFBSs), cytosine methylation patterns and steric regulators of cytosine methylation (exemplified by CGGBP1) have cooperatively given rise to the epigenomic diversity in amniotes.

## Materials and Methods

### Phylogenetic analysis

CGGBP1 DNA and protein sequences for various species were retrieved from NCBI, Ensembl and UniProt. The sequences were bifurcated into N-terminal and C-terminal parts by separating them at the NLS; sequence before and including the NLS was considered as the N-term and sequence after the NLS was considered as the C-term (Table S3). Multiple sequence alignment was done using MUSCLE at default parameters.

Phylogenetic tree construction was done using PhyML using maximum likelihood and other parameters at default settings. Tree rendering was done using FigTree version 1.4.4.

### CGGBP1 deletion constructs

N-term (1-90 aa) and FL (1-167 aa) with FLAG-tag at the N-terminal and GFP-tag at the C-terminal and C-term (79-167 aa) with HA-tag at the N-terminal and GFP-tag at the C-terminal were cloned in pEGFP-N3 expression vector (Addgene #6080-1) (Table S8) using *Xho*I and *Kpn*I sites. The clones were sequence-verified.

### Cell culture, shRNA transduction for knockdown and overexpression

HEK293T cells were cultured in DMEM (HiMedia #AL007A) supplemented with 10% FBS. Control and CGGBP1-targeting-shRNA (against 4 regions in the CGGBP1 ORF) were procured from Origene. Third-generation lenti-packaging plasmids: pRSV-Rev, pMDLg/pRRE and pMD2.G were procured from Addgene. For lentiviral production, the shRNA constructs and packaging plasmids were taken in equimolar ratios and used for transfection. Two different knockdown systems were created for overexpression of N-term and C-term: shRNA targeting the N-terminal of CGGBP1 (shA) was used to create HEK293T-shA-KD for overexpression of C-term; shRNA targeting the C-terminal of CGGBP1 (shB) was used to create HEK293T-shB-KD for overexpression of N-term; shControl was used to create HEK293T-CT for overexpression of FL-CGGBP1. Transfection was performed using JetOptimus (Polyplus #101000051). Polybrene (Sigma #TR-1003) was used for transducing cells. Transduced cells were selected using Puromycin (Himedia #CMS8861). After knockdown, a second round of transfection was done using pEGFP-N3 overexpression plasmids (5µg plasmid per 10cm dish) using JetOptimus.

### Immunofluorescence

Truncated and full-length CGGBP1 immunofluorescence was performed in HEK293T cells following standard protocol. Firstly, cells transfected with N-term, C-term and FL CGGBP1 were fixed in 4% formaldehyde solution at 37℃ for 10 min. Then cells were permeabilized using 1% Triton X-100 at room temperature for 10 min. After that, the cells were incubated in blocking solution (10% FBS and 0.05% Triton X-100 in PBS) for 1 h at room temperature. Following that, cells were incubated with primary antibodies (ab176814 antibody for N-term, pa5-57916 antibody for C-term and sc-376482 for FL) for 1 h at room temperature. Cells were then washed using washing solution (0.05% Triton X-100 in PBS). Cells were then incubated with secondary antibodies (anti-mouse Alexa fluor 594 for FL and anti-rabbit Alexa fluor 594 for N-term and C-term) for 1 h at room temperature. Cells were then stained with Fluoroshield mounting medium (Ab104135) containing DAPI. Cells were imaged using TCS SP8 (Leica) confocal microscope. Image analysis and signal quantification was done using Fiji ImageJ. Signal intensity graphs were plotted using GraphPad Prism (Mean with SEM).

### Nuclear cytoplasmic fractionation assay

Cells transfected with truncated and full-length forms were pelleted using centrifugation. The pellets were gently resuspended in 1x Cytoplasmic extraction buffer (10 mM HEPES, 60 mM KCl, 1 mM EDTA, 0.075% v/v NP-40, 1mM DTT, 1 mM PMSF) containing 1x Halt Protease inhibitor cocktail - EDTA-free (Thermo Fisher Scientific #87785) using cut tips and was incubated on ice for 2 mins. The slurry was spun at 1200 rpm and the supernatant was collected as cytoplasmic fraction in a fresh tube. The pellet was then washed with 1x washing buffer (10 mM HEPES, 60 mM KCl, 1 mM EDTA, 1mM DTT, 1 mM PMSF) to remove any contamination of the cytoplasm. The pellet was then subjected to 1x nuclear extraction buffer (20 mM Tris Cl, 420 mM NaCl, 1.5 mM MgCl2, 0.2 mM EDTA, 1 mM PMSF, 25% (v/v) glycerol) and salt concentration was adjusted to 400mM NaCl. The nuclei were resuspended using a vortex and incubated on ice for 10 mins with intermittent vortexing and the lysate was called the nuclear fraction. Both the fractions were cleared using centrifugation and were taken forward for SDS PAGE followed by western blot as described below. Antibody used for probing was ProteinTech-10716-1-AP.

### Co-immunoprecipitation

HEK293T cells co-transfected with N-term & C-term, N-term & FL and C-term and FL were lysed using a lysis buffer of composition: 50mM Tris (pH 8.0), 150mM NaCl, 1% NP40, 2mM EDTA, 1mM PMSF and containing 1x Halt Protease inhibitor cocktail - EDTA-free (Thermo Fisher Scientific #87785). This was followed by clearing cell lysates by centrifugation. Lysates were pre-cleared by Protein G Plus/Protein A Agarose Suspension (Merck Millipore #IP05). The lysates were then separately incubated with truncated form antibodies (i. For N&C CoIP, ab176814 antibody for N-term and pa5-57916 antibody for C-term; ii. For N&FL CoIP, ab176814 antibody for N-term and pa5-57916 antibody for FL; iii. For C&FL CoIP, pa5-57916 antibody for C-term and ab176814 antibody for FL) overnight at 4℃. This was followed by incubating lysates with Protein G Plus/Protein A Agarose Suspension for 2 h at 4 °C. The protein-bound Agarose beads were then washed with PBS four times. The protein was eluted from the beads by boiling in SDS-Laemmli buffer. This was followed by SDS PAGE and western blot as described below with the aforementioned antibodies.

### Western blot

4% stacking gel and 10% resolving gel were used to resolve the samples; transfer was done on to PVDF membrane; blocking for 1 h was done in blocking buffer (5% dry milk w/v in 1x TBST buffer) followed by incubation with primary antibody overnight at room temperature for 2 hrs (1:100 dilution in blocking buffer). Membranes were washed in 1x TBST, incubated with HRP conjugated anti-rabbit secondary antibody (1:5000 dilution in blocking buffer) for 1 hr at room temperature followed by washing with 1x TBST. Signals were developed using ECL substrate. Blots were imaged in the chemiluminescence mode using BioRad ChemiDoc MP Imaging System (BioRad #12003154).

### Microarray-based gene expression analysis

HEK293T-shB-KD, HEK293T-shA-KD and HEK293T-CT cells were used to overexpress N-term, C-term and FL-CGGBP1 respectively. Cells were collected in triplicates and pellets were frozen at -80℃ for RNA extraction using Trizol and purification by Qiagen’s RNeasy mini kit (Qiagen #74106). Agilent’s Quick-Amp labelling Kit (Agilent #5190-0424) was used for T7 promoter based-linear amplification to generate labelled complementary RNA (Cy3) for one-colour microarray-based gene expression analysis. Agilent’s In situ Hybridsation kit (Agilent #5188-5242) was used for hybridisation on Human GXP 8X60k (AMADID: 072363) chip. Intra-array signal normalisation was done using GeneSpring GX 14.5 Software. Intra-group quantile normalisation for triplicates for all three samples - N-term, C-term and F-len was performed before analysing the data.

### Methylcytosine DNA immunoprecipitation (MeDIP)

HEK293T-shB-KD, HEK293T-shA-KD and HEK293T-CT cells were used to overexpress N-term, C-term and FL-CGGBP1 respectively. After 72 h of transfection, cells were harvested for genomic DNA isolation using phenol-chloroform-isoamyl alcohol extraction method. Genomic DNA was sonicated to obtain 700-1000 bp fragments. 1µg fragment DNA was end repaired using NEBNext Ultra II End Repair/dA-Tailing Module (NEB #E7546). PCR adapters from Oxford Nanopore Technologies (ONT #SQK-PSK004) were ligated to end repaired DNA using NEB Blunt/TA Ligase Master Mix (NEB #M0367). 1x MeDIP master mix (10 mM Sodium phosphate buffer, 0.14 M NaCl and 0.05% TritonX-100) was added to adapter ligated DNA which was then denatured at 95℃ for 5 mins followed by snap chilling on ice. An antibody cocktail against 5-methylcytosine (EMD Millipore #MABE146, Sigma #SAB2702243 and Novus Biologicals #NBP2–42813) was added to snap-chilled DNA and was incubated at 4℃ overnight. Protein G Plus/Protein A Agarose beads (Merck Millipore #IP05) were added to the mix followed by incubation at room temperature for 2 h. Then beads were collected by centrifugation in a separate tube and were washed thrice with 1x MeDIP master mix and were subjected to Proteinase K (Sigma #P2308) digestion at 56℃ for 2 h. The slurry was centrifuged and immunoprecipitated DNA in the supernatant was collected into new tubes. This DNA was subjected to 18 cycles of PCR using whole genome primers (ONT #SQK-PSK004) and LongAmp Hot Start Taq 2X Master Mix (NEB #M0533). Library preparation was done using NEBNext® UltraTM II DNA Library Prep Kit for Illumina® (NEB #E7645, #E7103). DNA was sequenced on Illumina HiSeq 2500 platform.

### Chromatin immunoprecipitation (ChIP)

HEK293T-shB-KD, HEK293T-shA-KD and HEK293T-CT cells were used to overexpress N-term, C-term and FL-CGGBP1 respectively. After 72 h of transfection, cells were harvested for ChIP. Cells were crosslinked with 4% formaldehyde at 37°C for 10 min followed by quenching with glycine (125 mM) and were then harvested using a scrapper. Lysis was done using SDS lysis buffer (50mM Tris pH 8.0, 10mM EDTA, 1% SDS) containing 1x Halt Protease inhibitor cocktail - EDTA-free (Thermo Fisher Scientific #87785) for 30 minutes on ice with intermittent flick-mixing. DNA was then sonicated for 20 cycles of 30 seconds ON/30 seconds OFF to obtain DNA fragments of 0.15 kb – 0.35 kb length. 10% volume of fragmented DNA was taken separately as input and the remaining was diluted in ChIP-dilution buffer (0.01% SDS, 1.1% Triton X-100, 1.2 mM EDTA, 16.7 mM Tris-HCl pH 8.1, 167 mM NaCl). Chromatin pre-clearing was done by incubating with Protein G Plus/Protein A Agarose beads (Merck Millipore #IP05) for 4 hrs at 4°C. The DNA was then incubated with an antibody cocktail (Invitrogen #PA5-57317, Abcam #ab176814, Abclonal #AE005 against the FLAG-tag attached to the N-term) targeting CGGBP1-N-term with mild tumbling at 4°C overnight. Subsequently, Protein G Plus/Protein A Agarose beads were added to capture antibodies and incubated for 1 hour at room temperature in tumbling condition. Beads were then pelleted and washed with four buffers in the order: low-salt (0.1% SDS, 1% Triton X-100, 2 mM EDTA, 20 mM Tris–HCl and 150 mM NaCl), high-salt (0.1% SDS, 1% Triton X-100, 2 mM EDTA, 20 mM Tris–HCl and 500 mM NaCl), LiCl (0.25 M LiCl, 1% IGEPAL (NP40), 1% sodium deoxycholate, 1 mM EDTA and 10 mM Tris–HCl) and two washes of TE buffer (10 mM Tris–HCl and 1 mM EDTA). Elution of immunoprecipitated DNA was done using an elution buffer (1% SDS and 0.1 M NaHCO3) in a total volume of 500ul. Reverse-crosslinking was done by adding 200mM NaCl and incubating at 65°C for 8 hrs. The DNA was then subjected to Proteinase K (Sigma #P2308) digestion for 1 hour at 45°C by adding 10mM EDTA, 0.1 M Tris-HCl pH 6.8 and 0.1 mg/ml Proteinase K). ChIP DNA was then purified by adding 10% v/v 3M sodium acetate pH 5.2 followed by ethanol precipitation. DNA was quantified using Qubit® dsDNA HS Assay Kit (Invitrogen, #Q32854). Library preparation was done using NEXTFlex Rapid DNA Sequencing Bundle (NEXTflex, #5188-12). DNA was sequenced on Illumina NovaSeq 6000 platform.

### Sequence data analysis

Adapter trimming and quality filtering were done using fastp version 0.23.4. Alignment at default parameters was done using bowtie2 version 2.5.1. Broad peaks were called using MACS2 version 2.2.7.1 without using any background. deeptools version 3.5.1 was used to plot signal profiles, and to compare multiple bigwigs. RepeatMasker version 4.1.0 was used to identify repeats. Motif search was performed using fimo version 5.5.4 (MEME Suite) at default parameters. RStudio was used to make volcano plots, bubble plots and heatmaps. Venny was used to calculate intersections between datasets. GraphPad Prism was used to perform t-tests and plot frequency distributions.

## Acknowledgement

The authors acknowledge Mr Aditya Teja and Ms Hema Naveena A for help with confocal microscopy, Dr Sweta Patel and Dr Vivek Tanavde for flow cytometry (unpublished data) and, Dr Dhiraj Bhatia and Dr Noopur Thakur for inputs in various experiments.

## Declaration

The authors declare no conflict of interest.

---

**Table S1.**
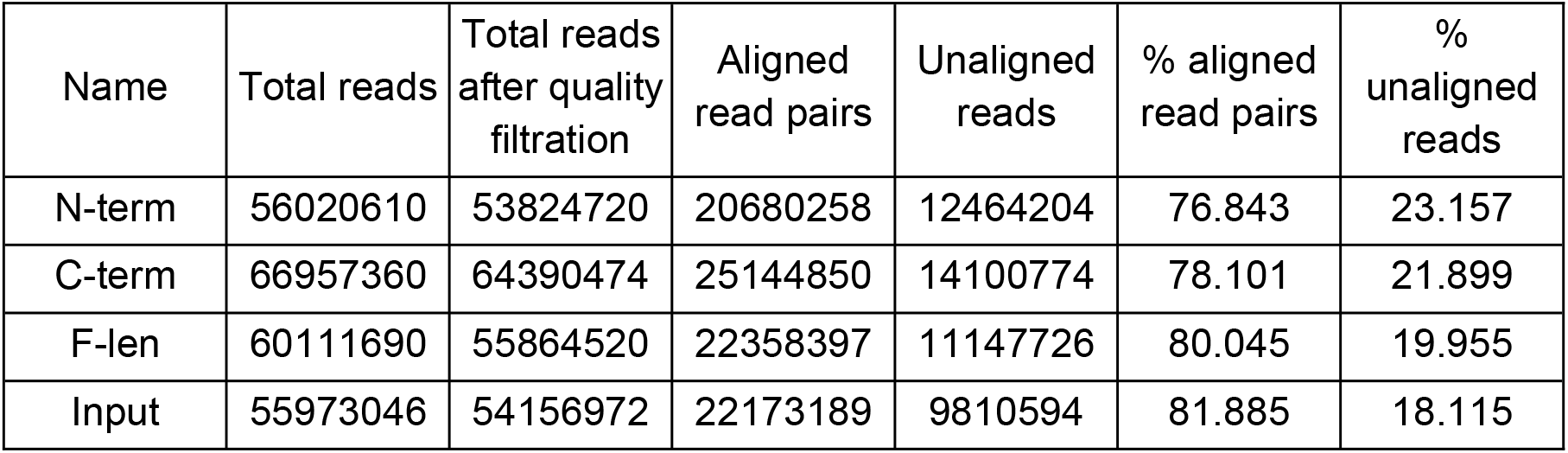
The table shows the number of raw MeDIP-sequencing reads, quality-filtered reads (using fastp at default parameters) and aligned or unaligned reads (after alignment with unmasked hg38 using Bowtie2; read length range constraint of 100bp-1000bp).

**Table S2.**
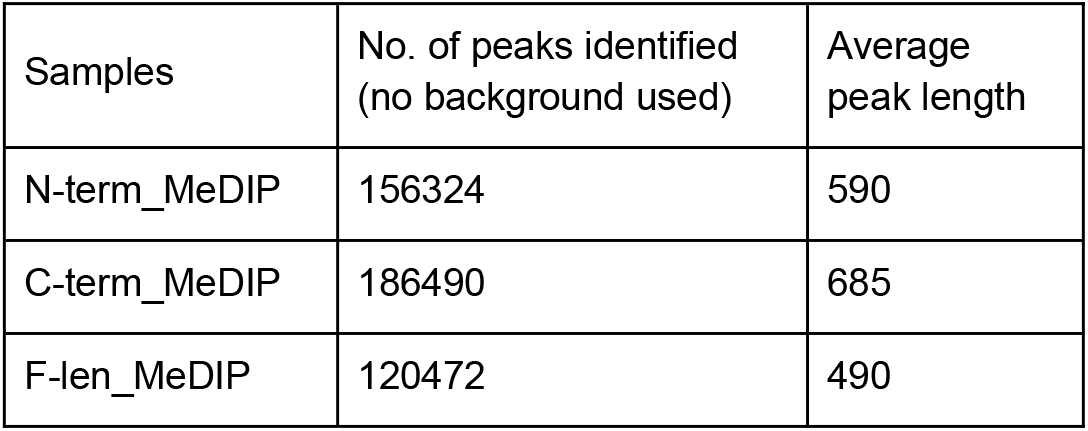
A summary of peaks identified in MeDIP reads by MACS2 (no background, no model) and average peak length.

**Table S3.**
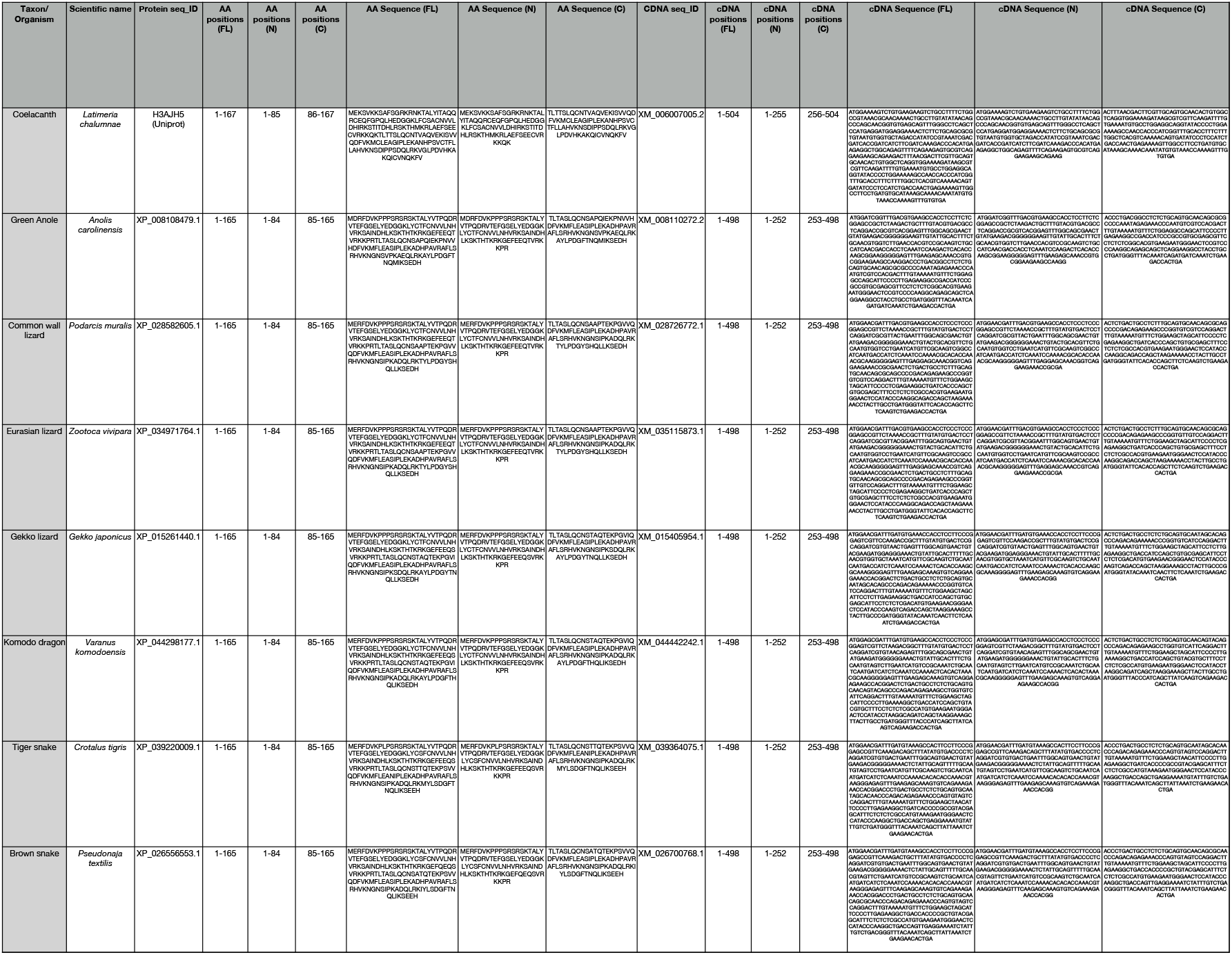
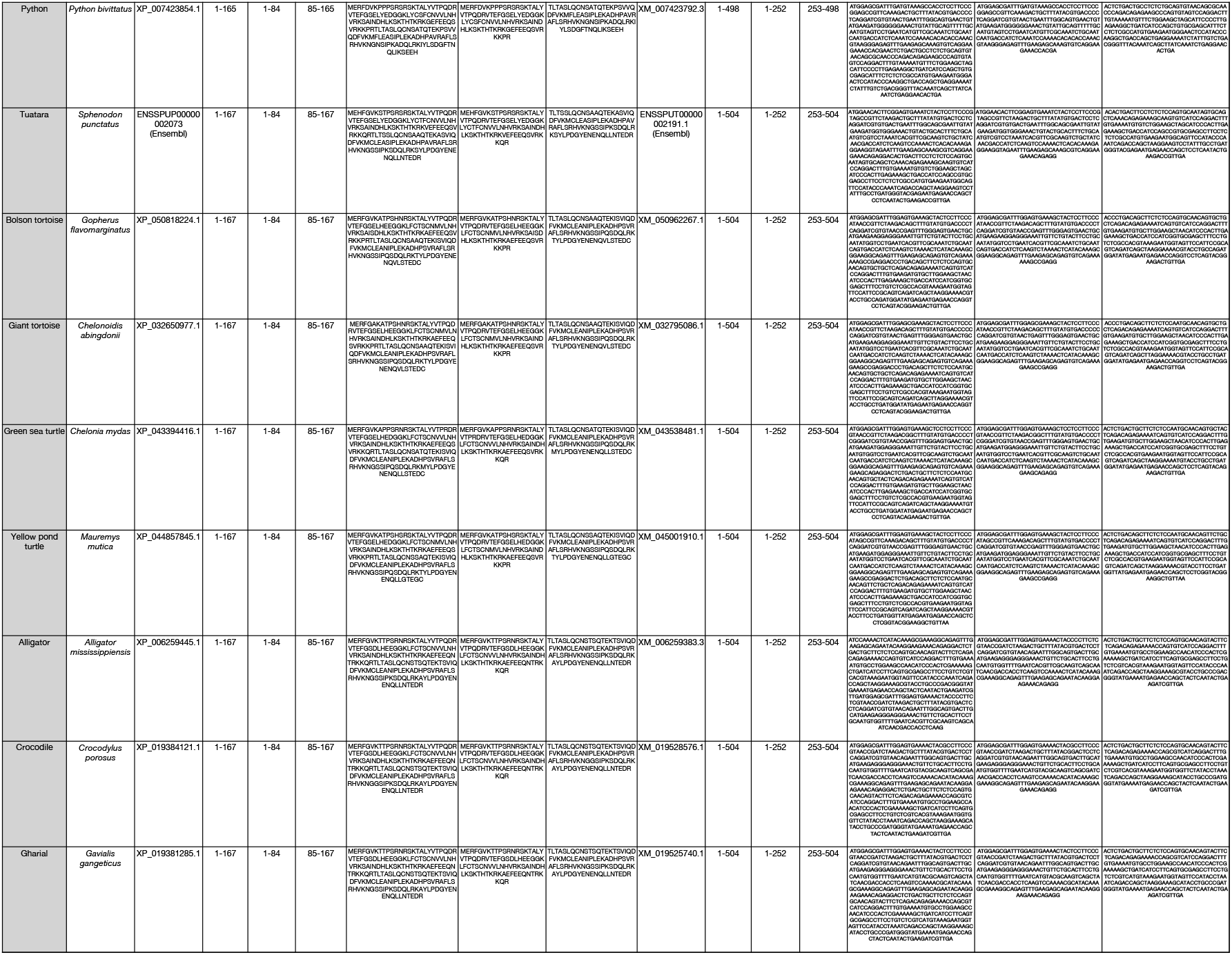
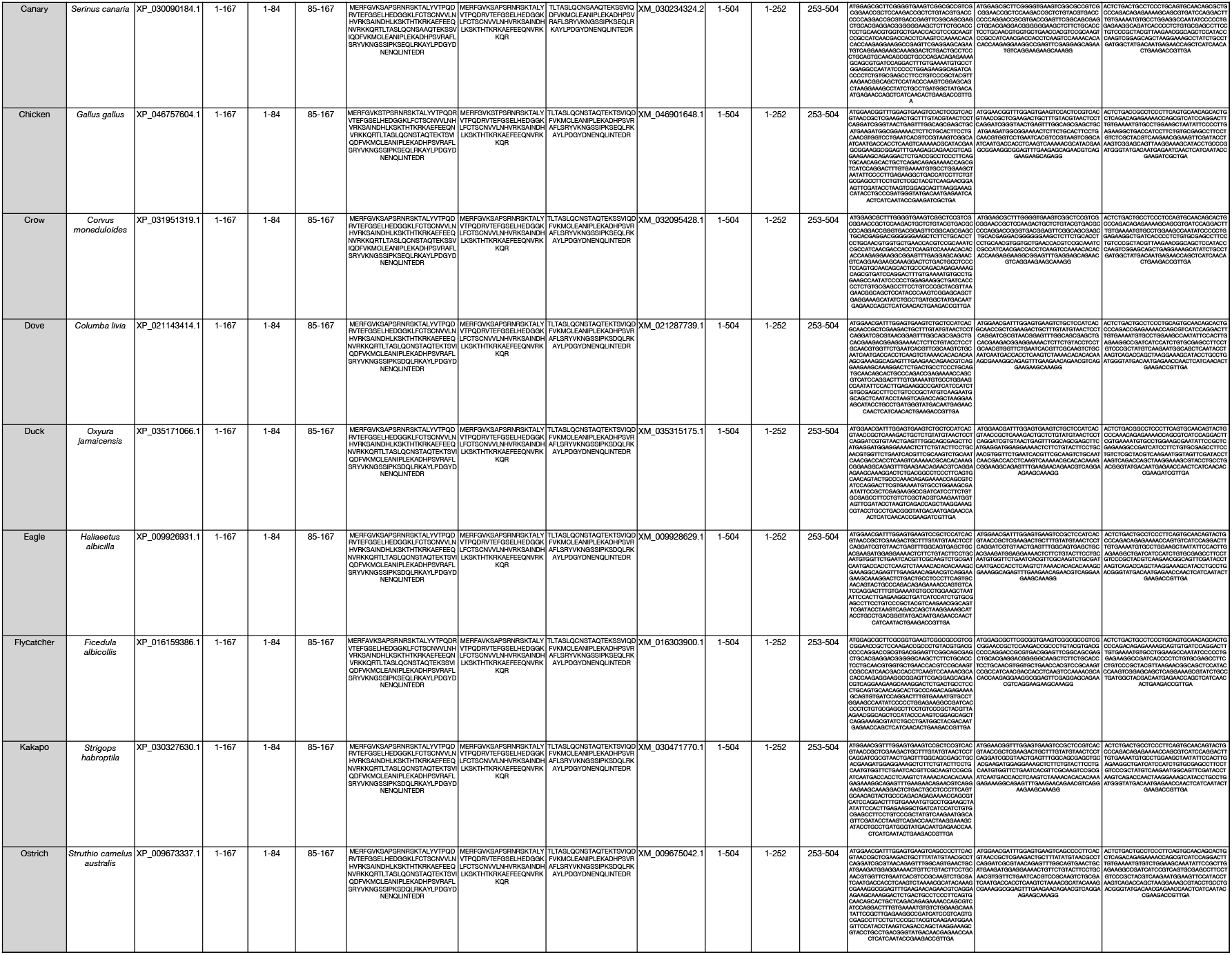
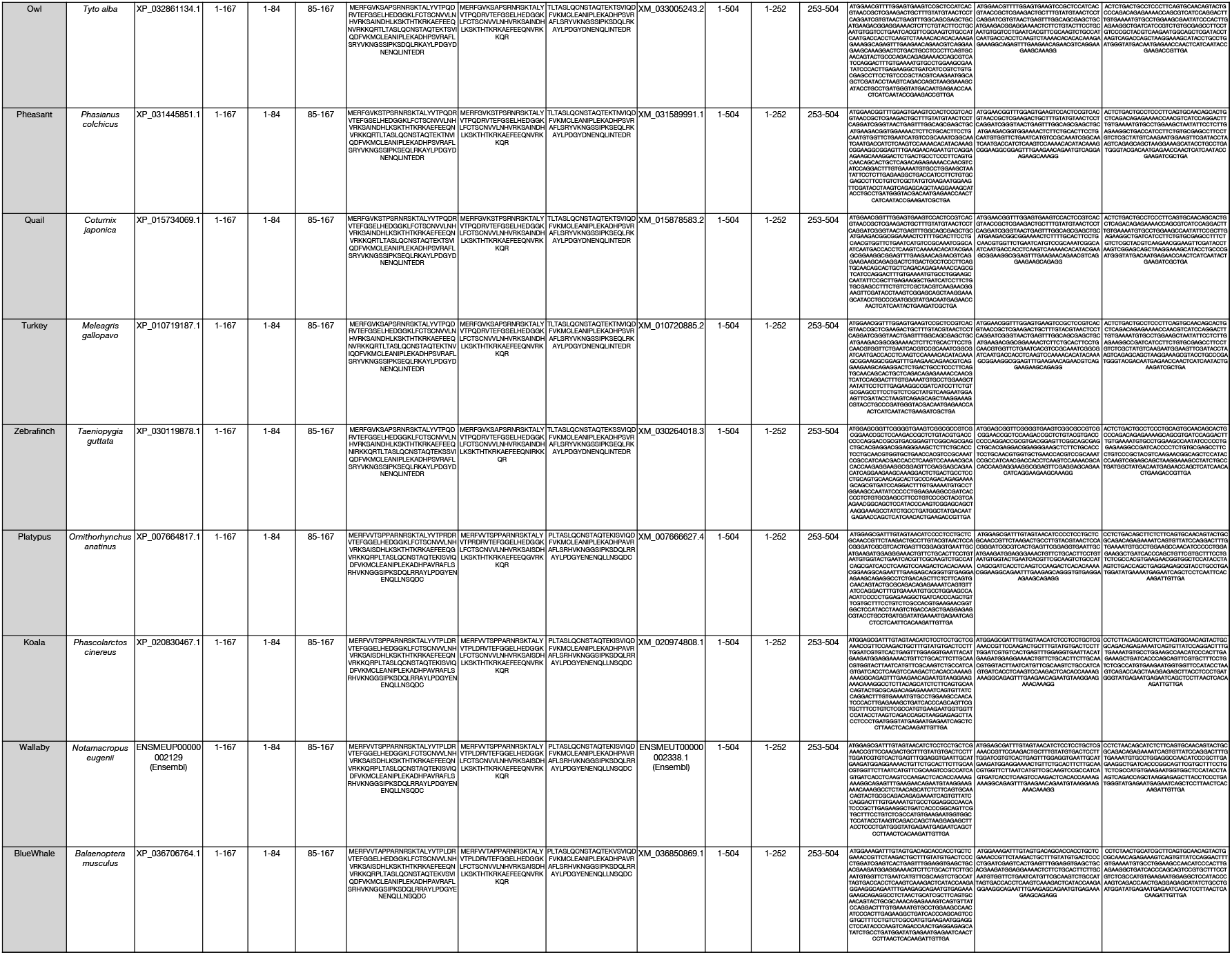
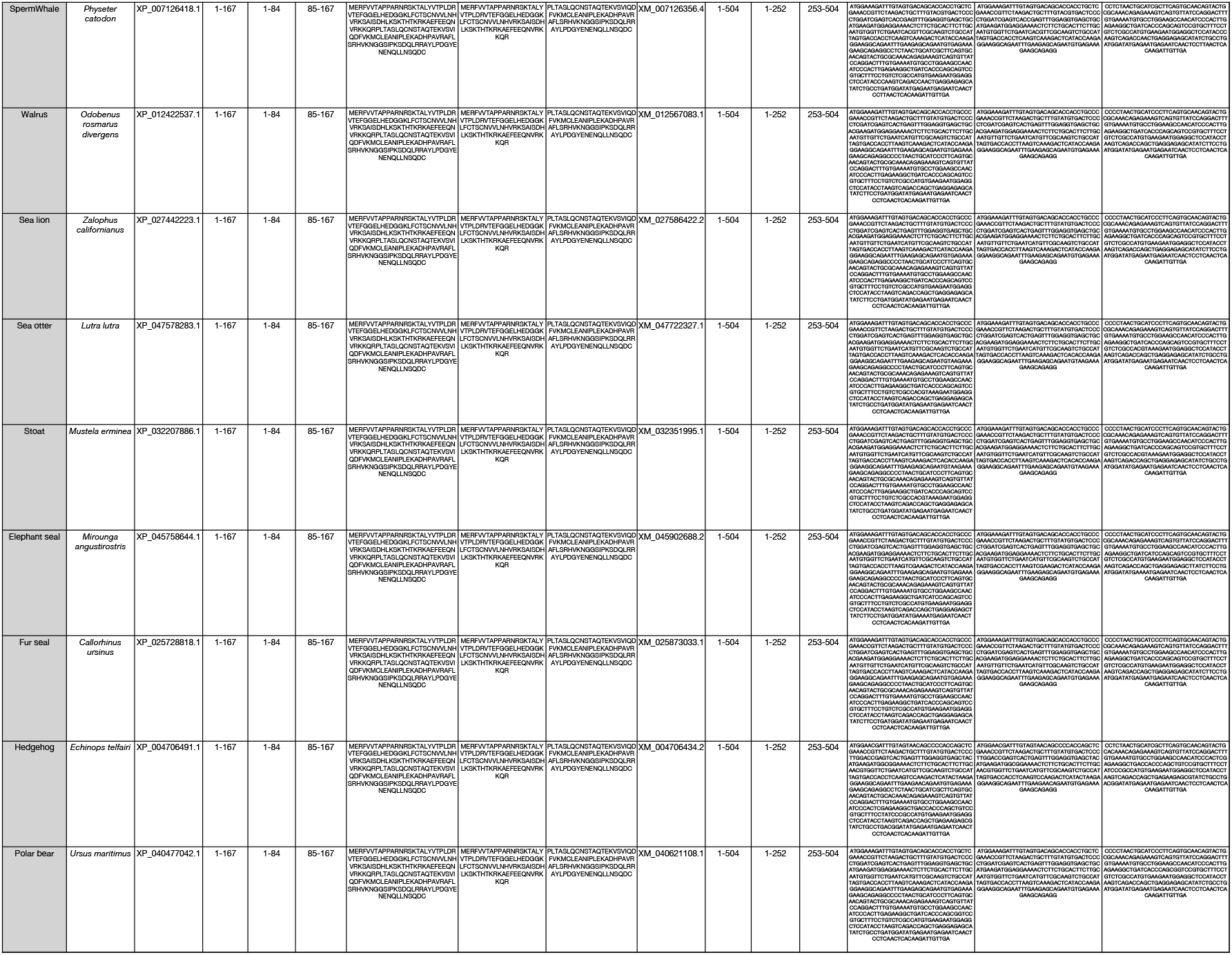
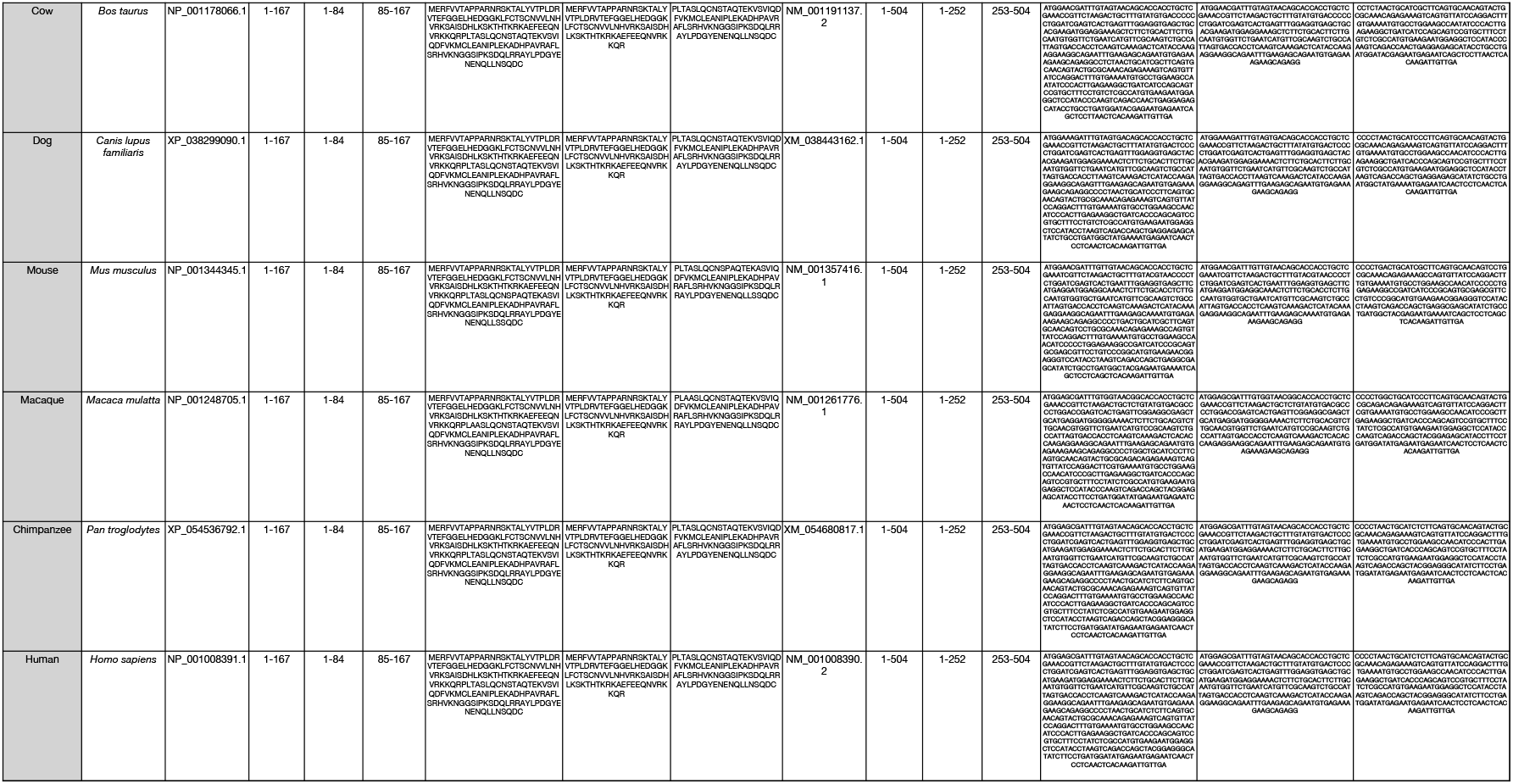
Details of CGGBP1 cDNA and protein sequences from different species available from NCBI, Ensembl or Uniprot that have been used to make phylogenetic trees have been compiled.

**Table S4.**
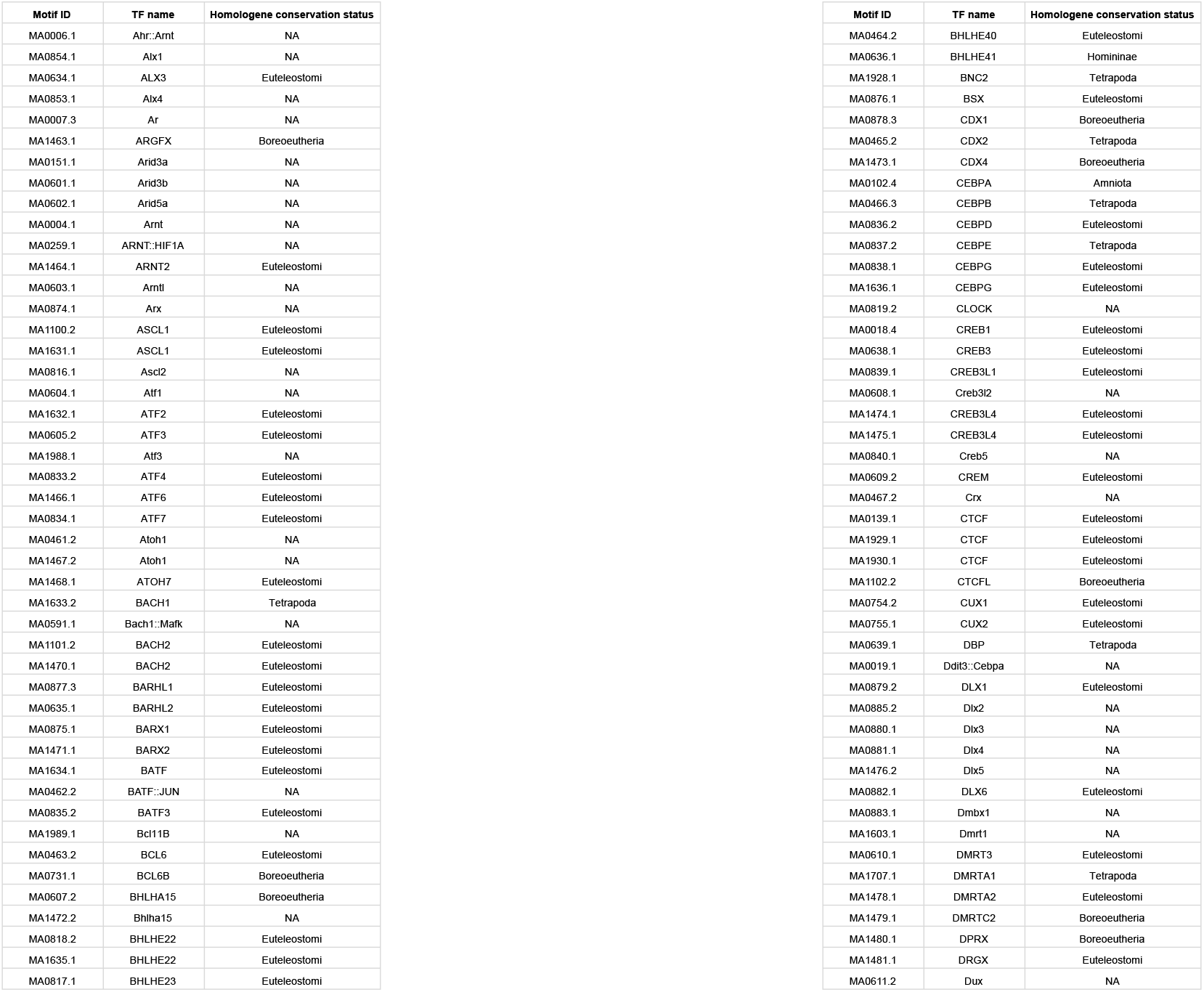
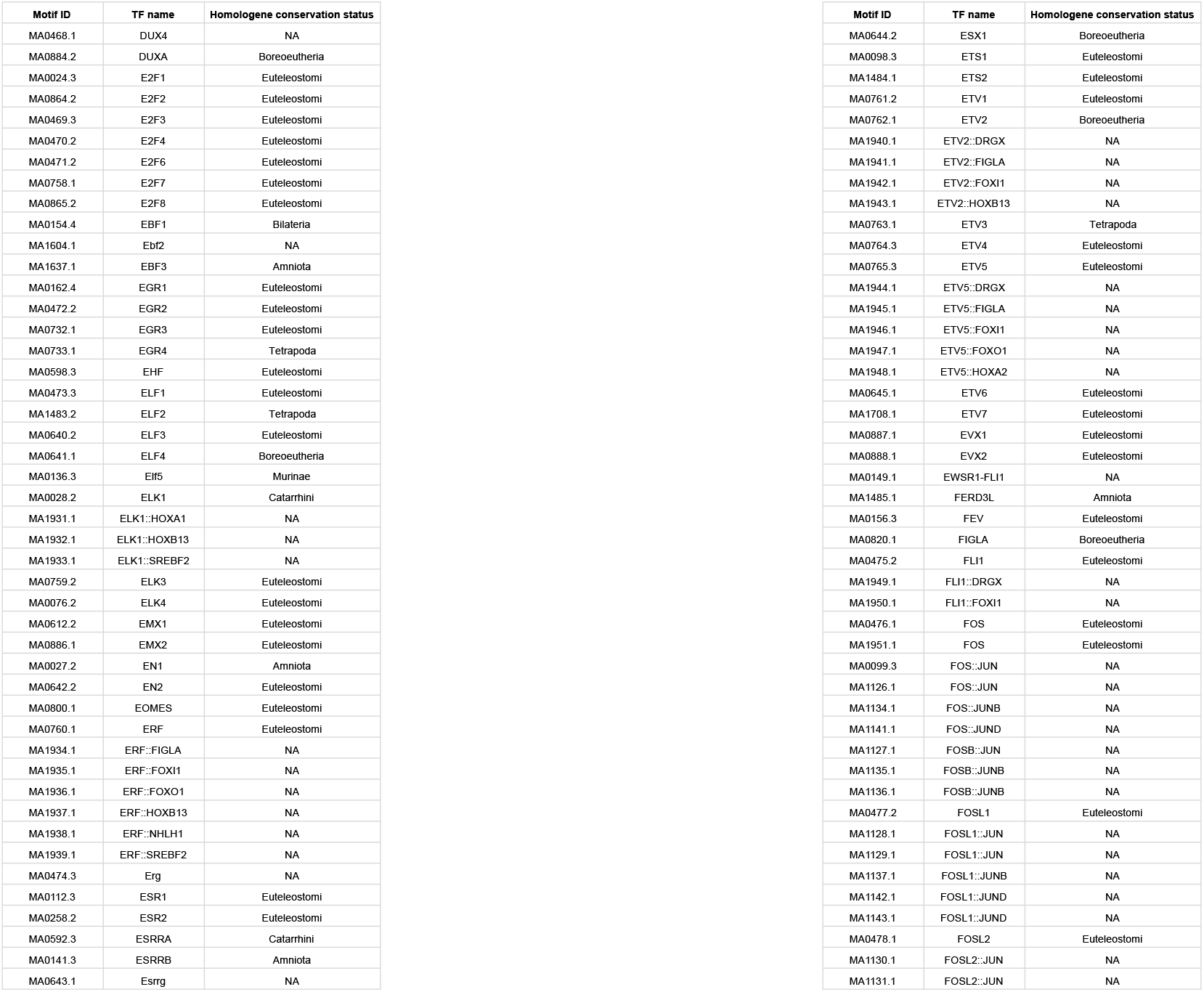
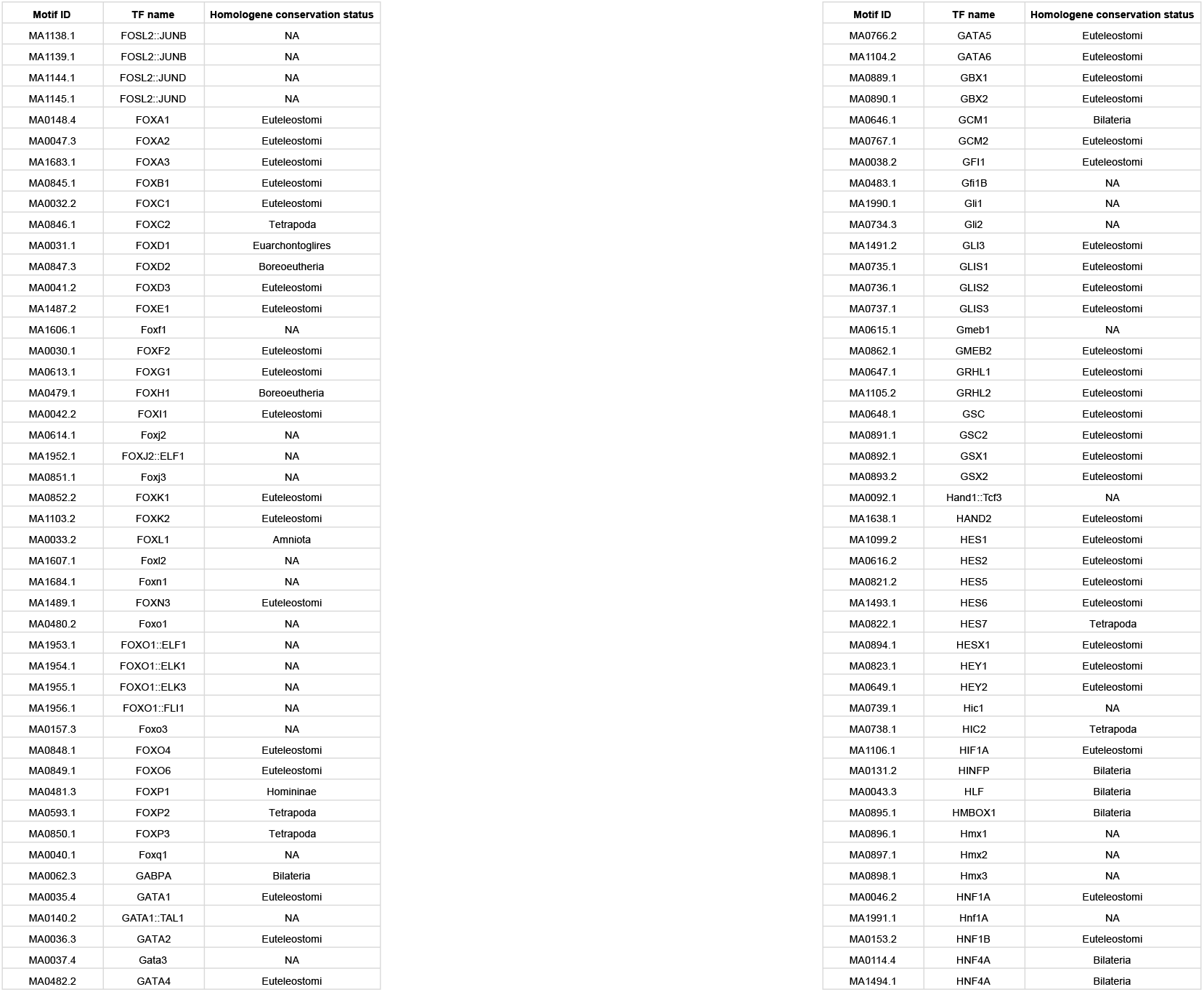
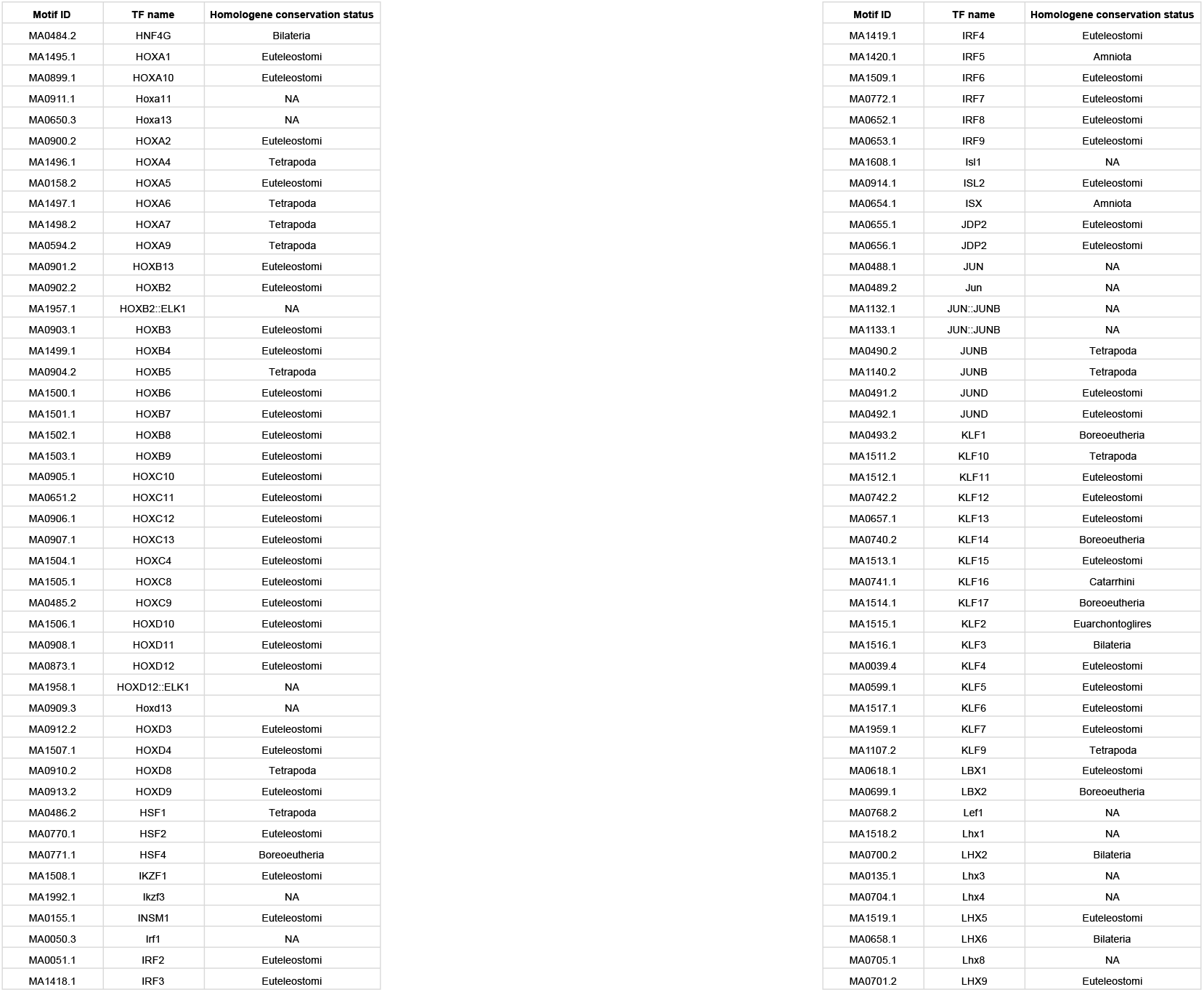
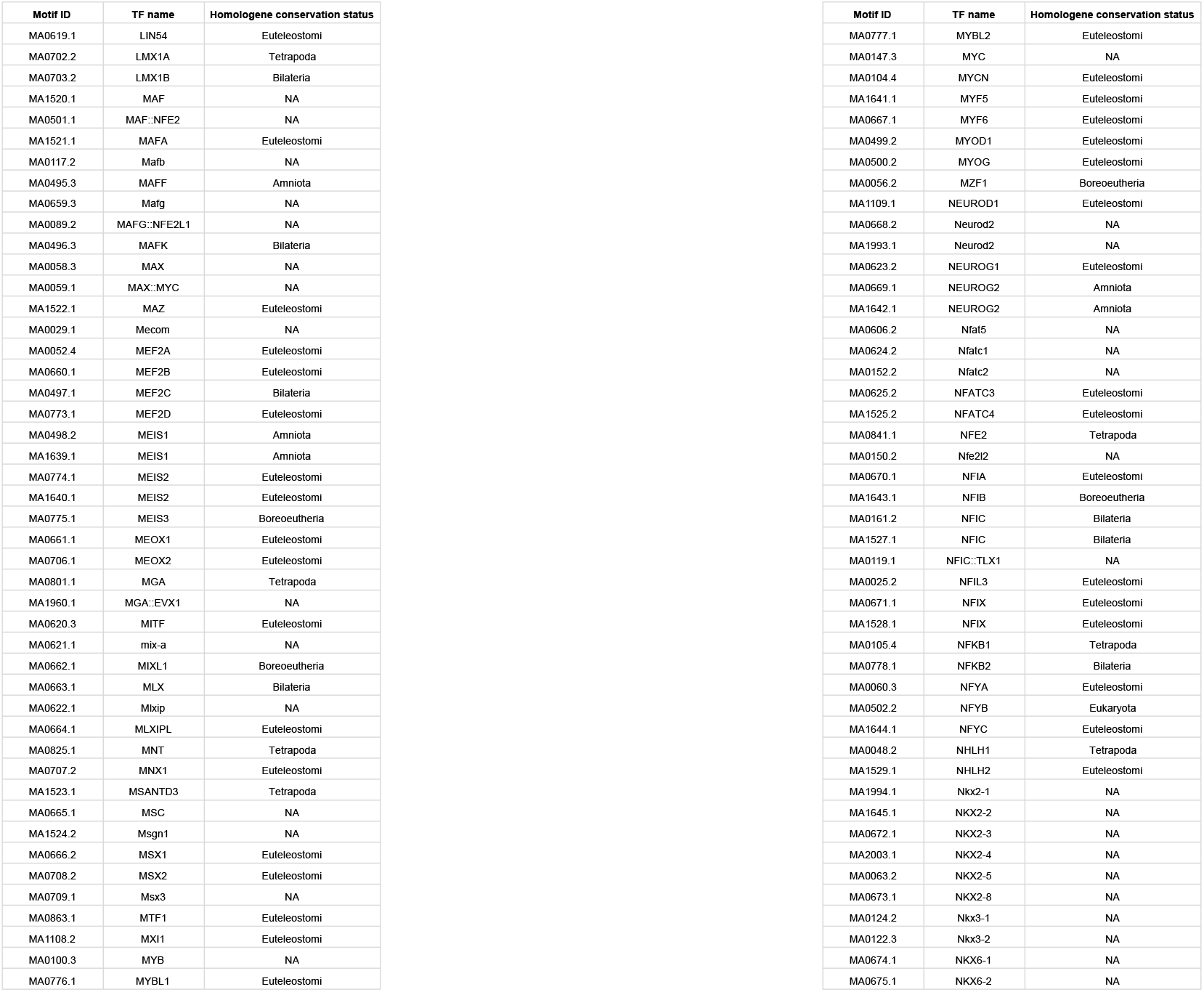
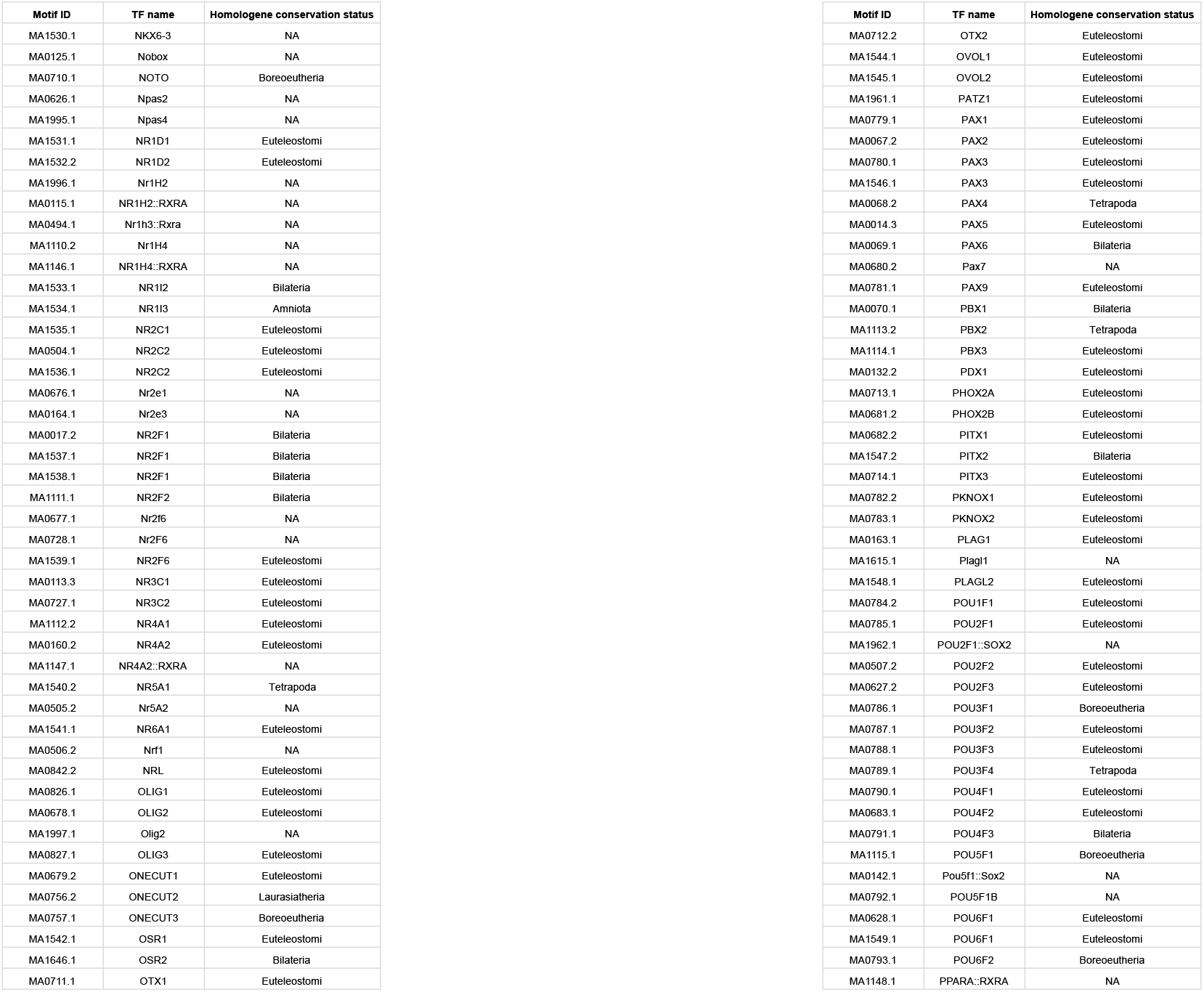
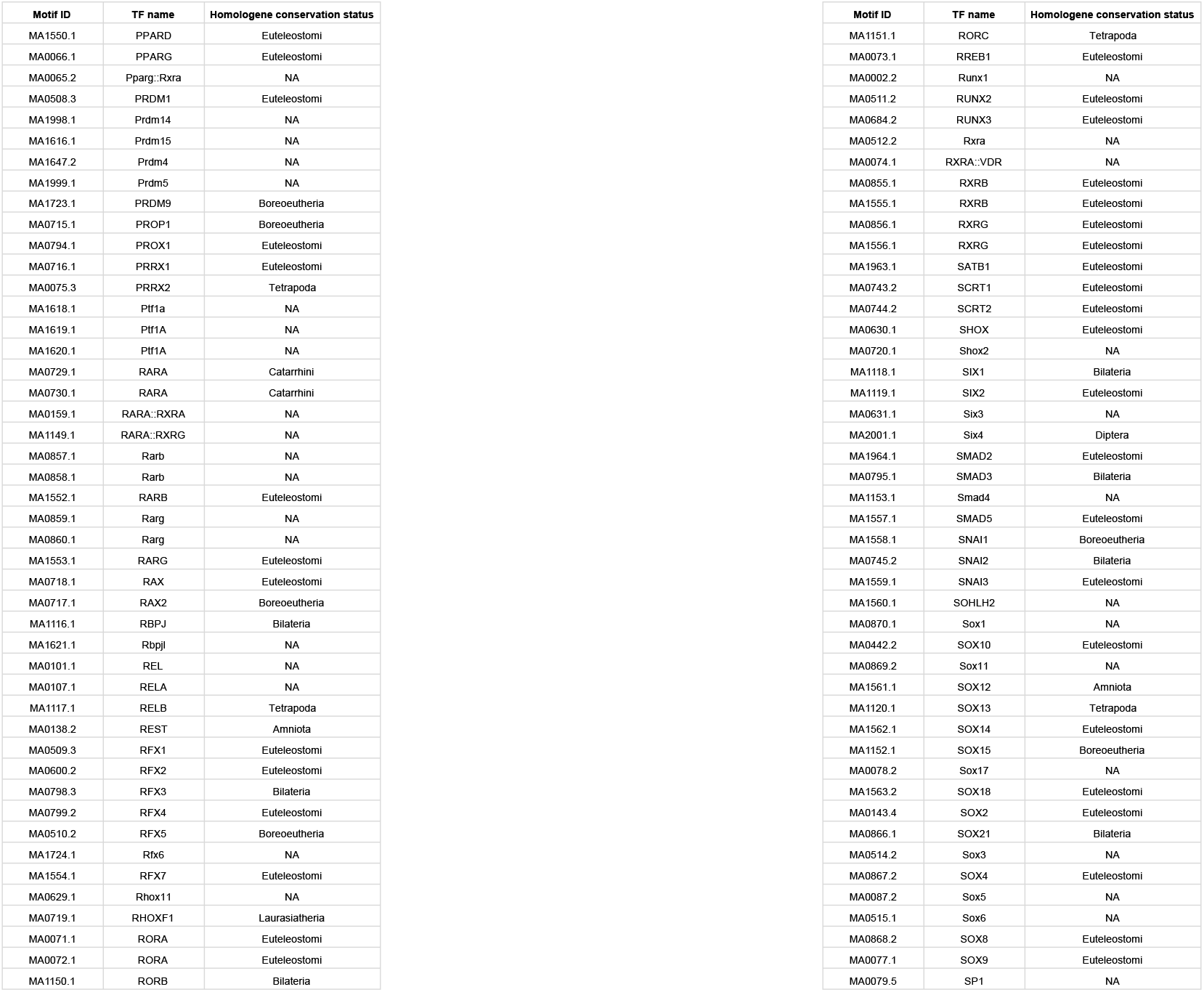
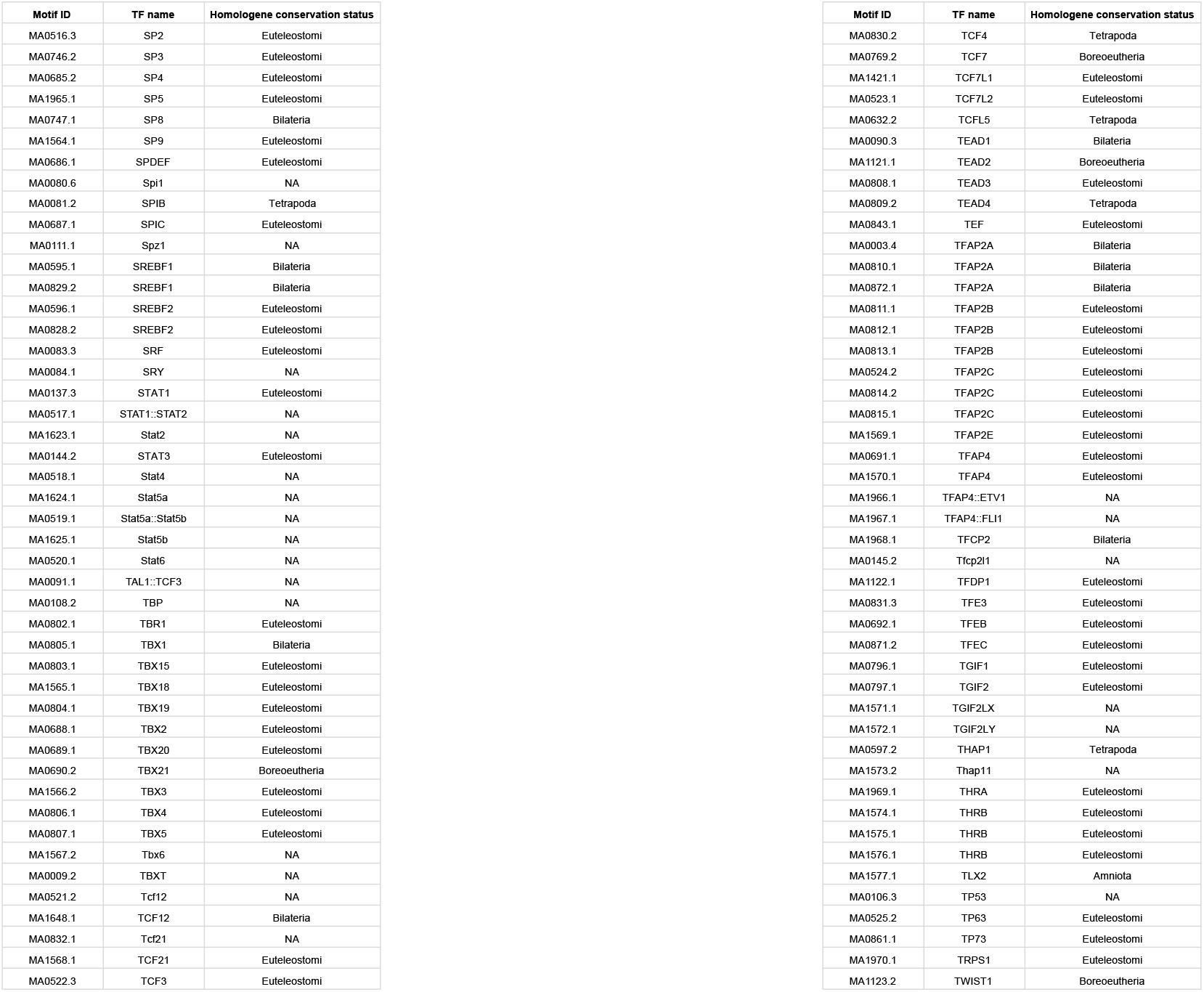
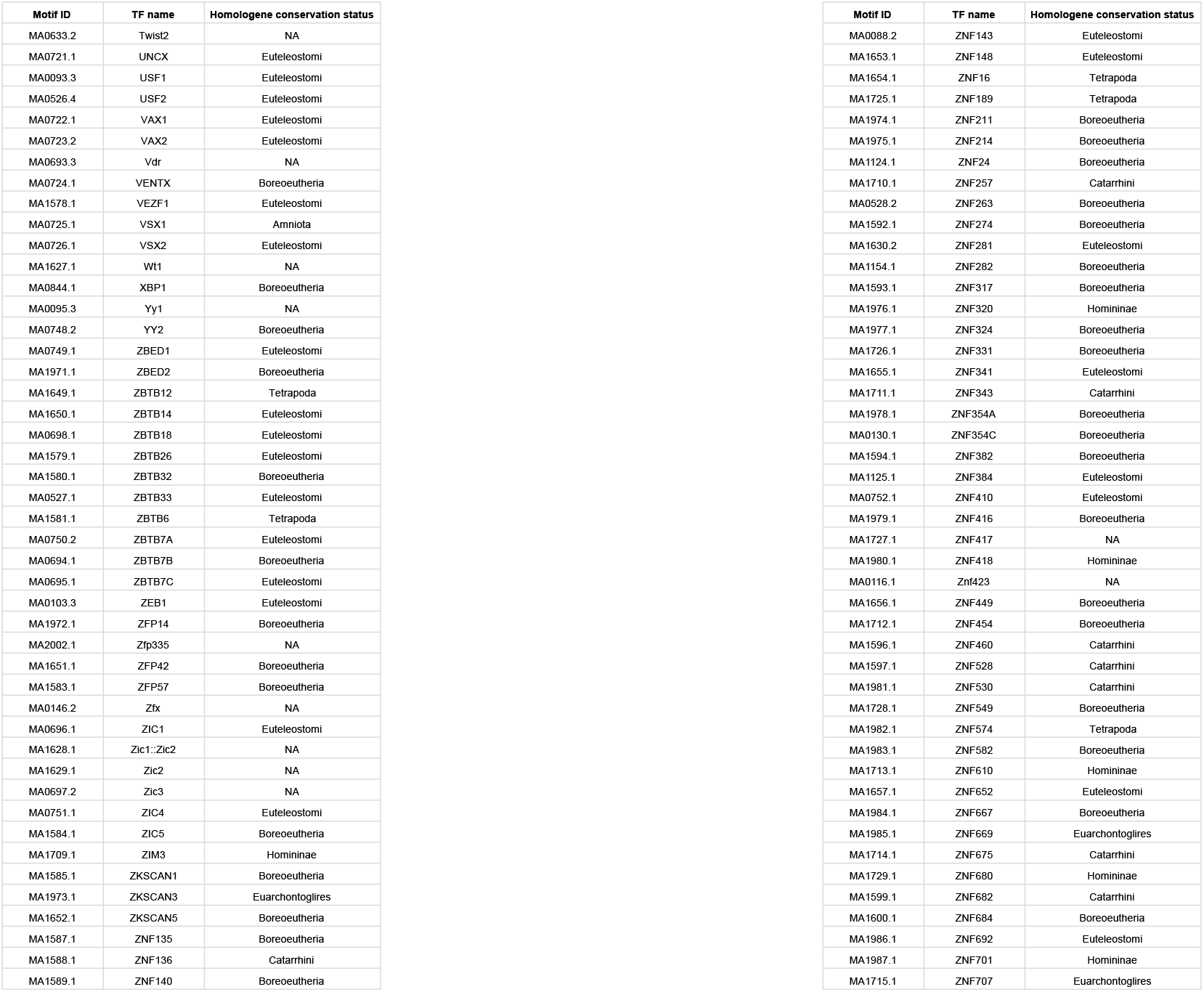
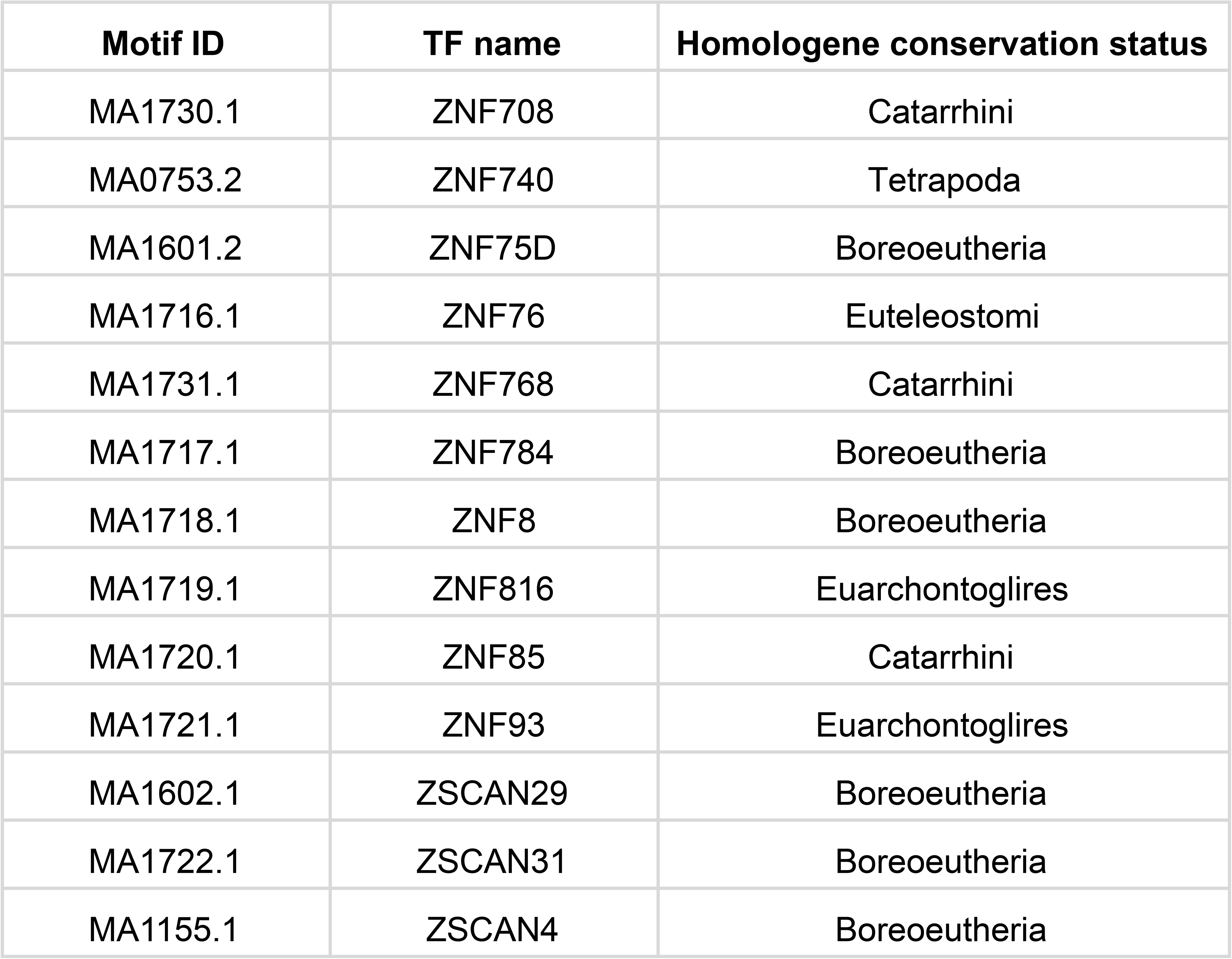
The conservation status of all the transcription factors in JASPAR taken from legacy Homologene has been enlisted.

**Table S5.** List of probe identifiers of differentially expressed genes in N-term vs F-len, C-term vs F-len and N-term vs C-term comparisons with fold change and associated p-values.

**Table S6.** The table contains genomic coordinates (hg38) of C-repressed genes along with their Ensembl gene and transcript identities.

**Table S7.**
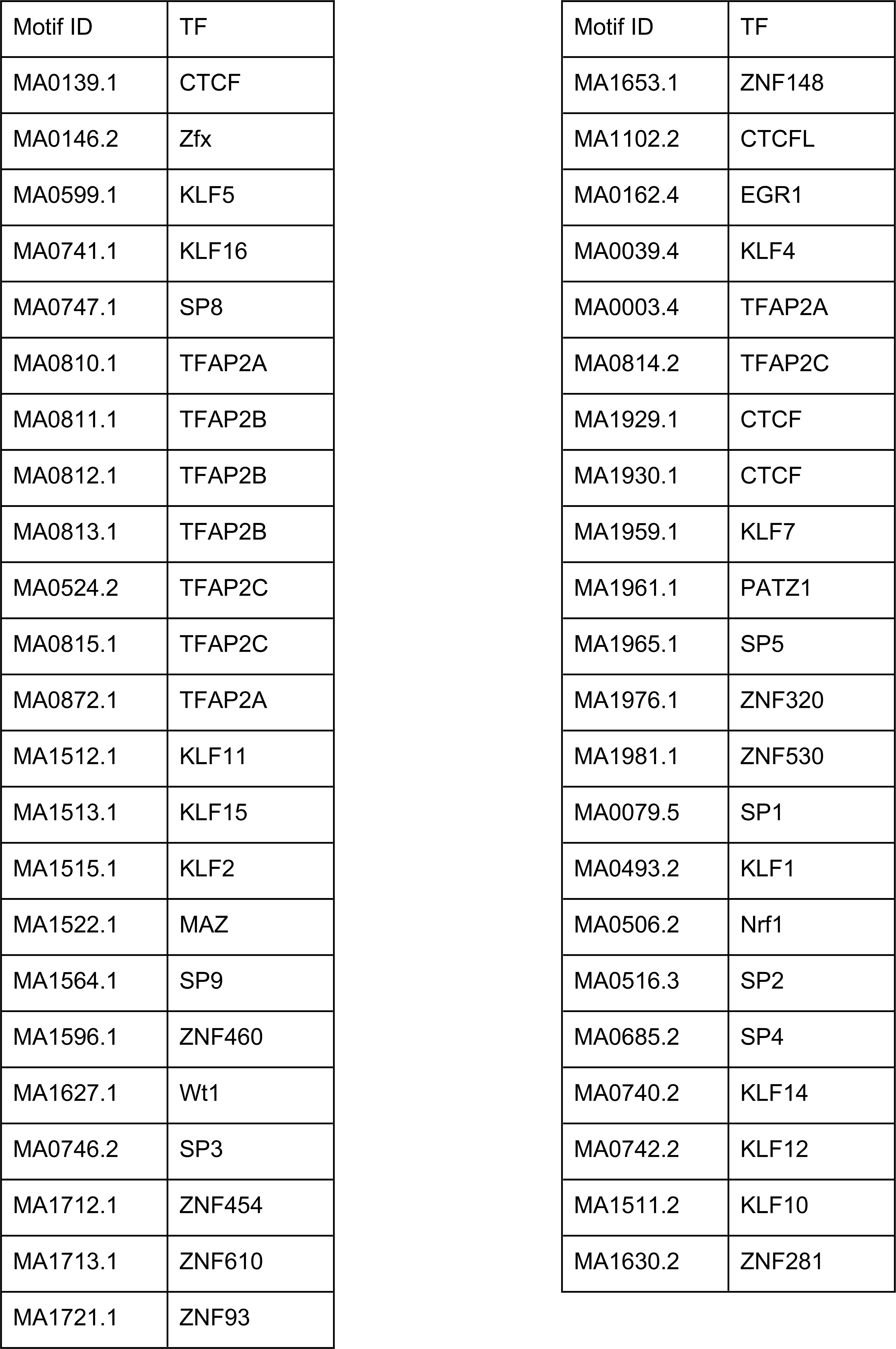
The JASPAR identities of 45 motifs with low obs-exp occurrence differentials in mammals have been enlisted.

**Table S8.**
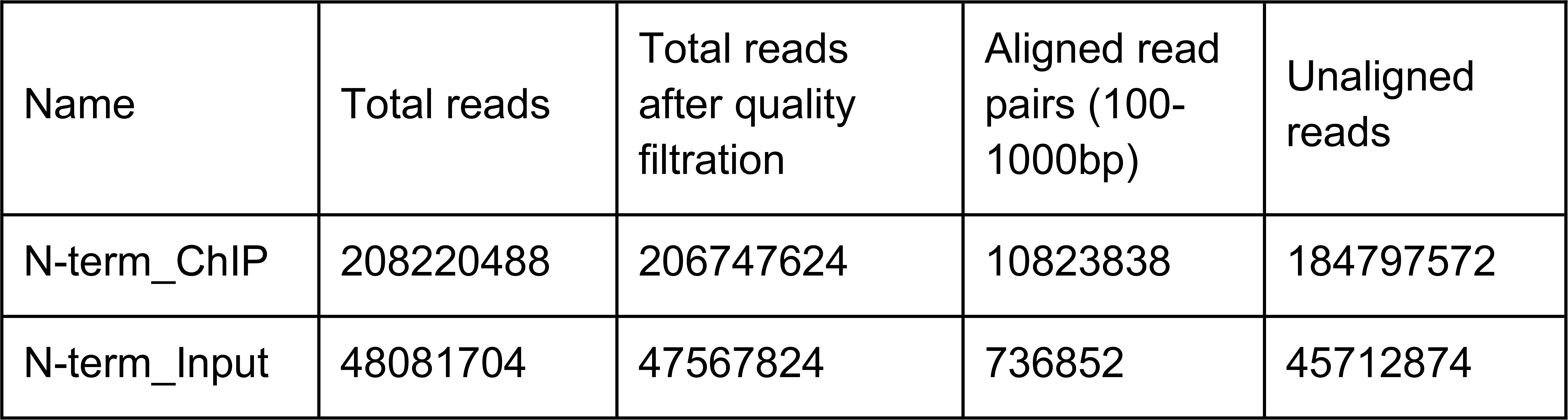
The table shows the number of raw ChIP-sequencing reads, quality-filtered reads (using fastp at default parameters) and aligned or unaligned reads (after alignment with unmasked hg38 using Bowtie2; read length range constraint of 100bp-1000bp).

**Table S9.**
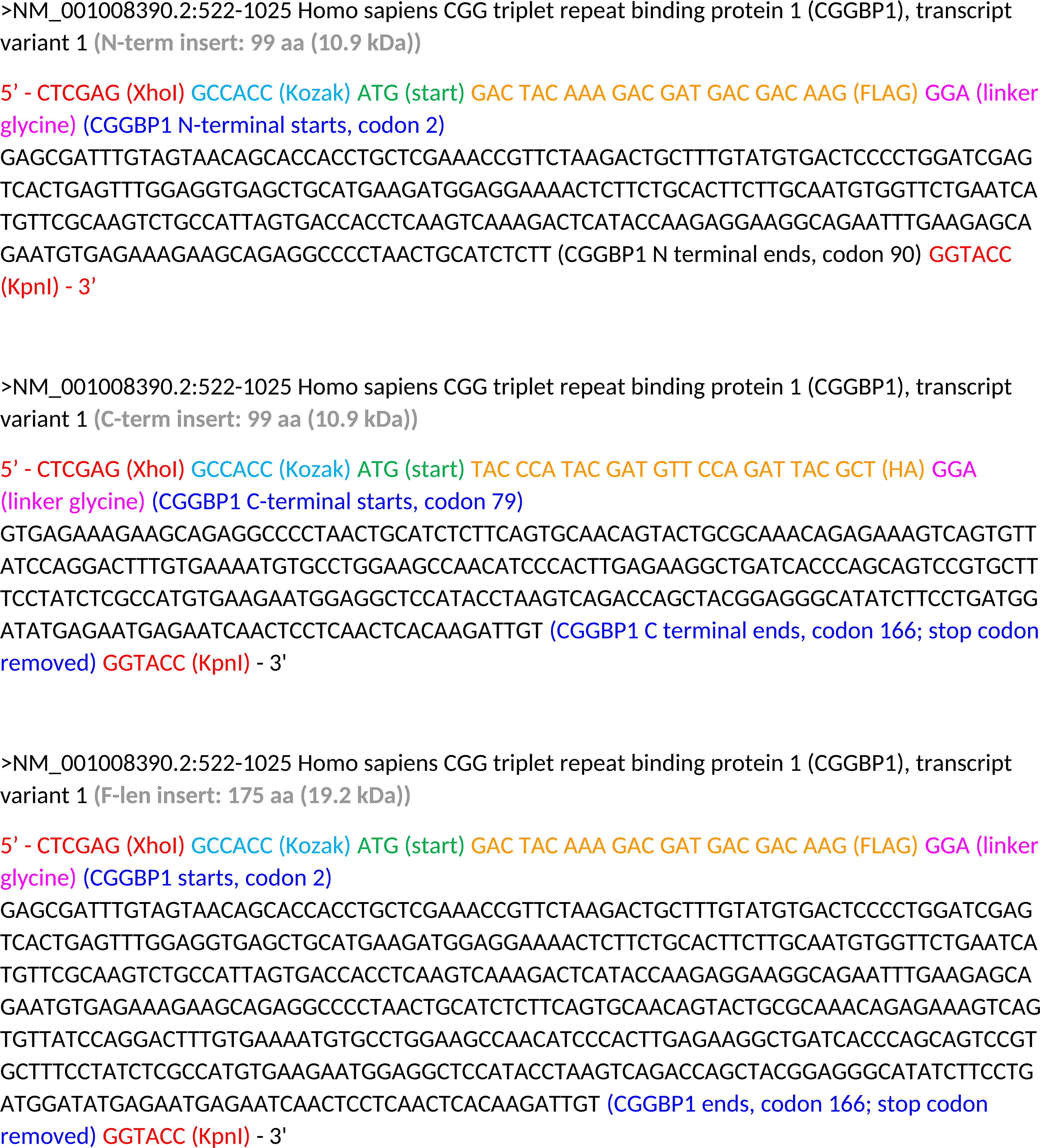
Details of CGGBP1 truncated forms and full-length constructs inserted in pEGFP-N3 over-expression plasmid.

**Fig S1.**
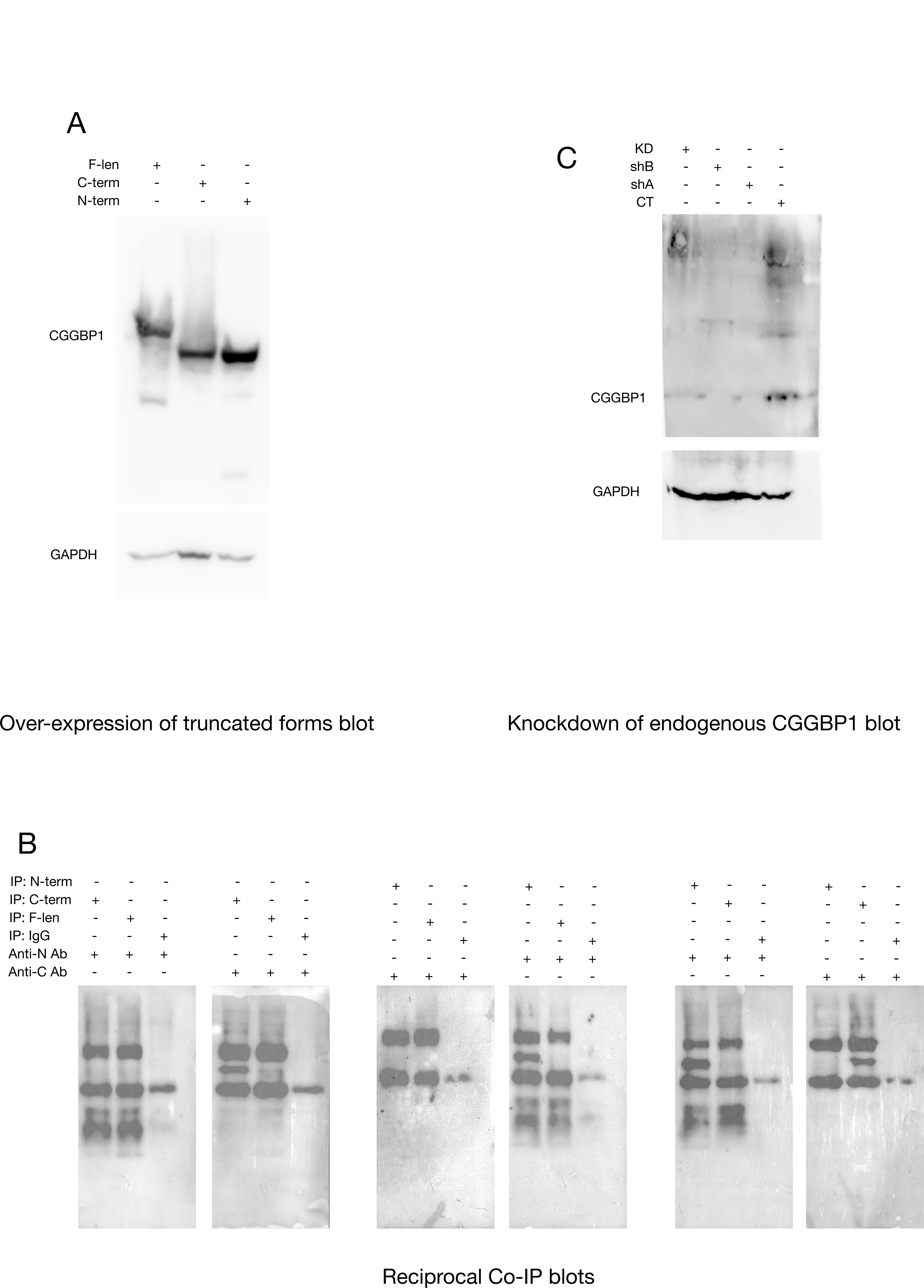
(A) Immuno-blot for N-term, C-term and F-len over-expression and corresponding control GAPDH; (B) Immuno-blots for reciprocal Co-IPs between N-term–C-term, C-term–F-len and N-term–F-len along with IgG control showing no physical interaction among the truncated forms or the truncated forms and the full-length protein; (C) Immuno-blot demonstrating knockdown of endogenous CGGBP1 with shA and shB shRNA as compared to the control shRNA (CT) along with the corresponding control GAPDH.

**Fig S2.**
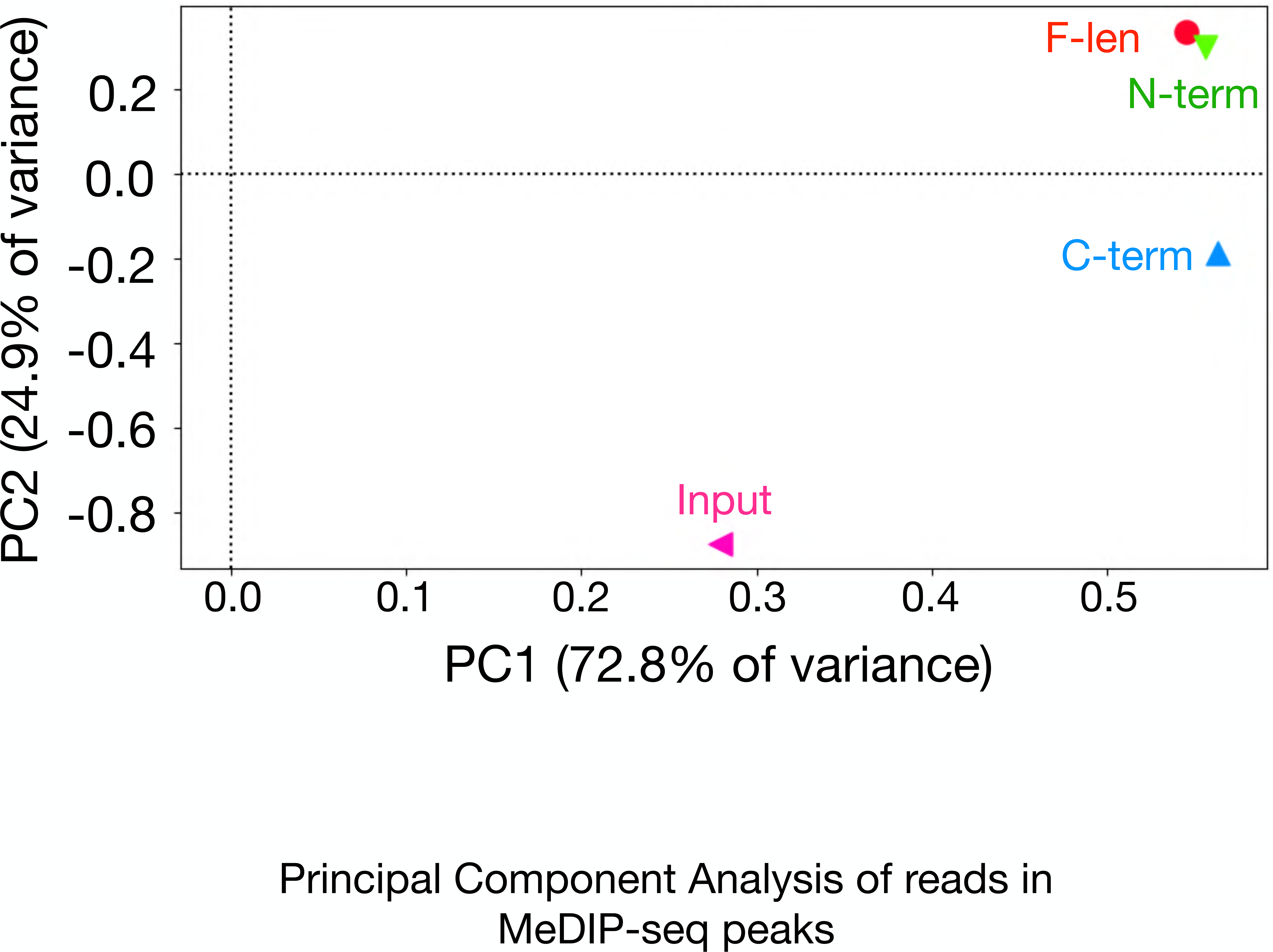
Principal Component Analysis of reads in MeDIP-seq peaks: MeDIP signals show enrichment in N-term, C-term and F-len as compared to input accounting for 72.8% variance (PC1) which does not differentiate the MeDIP samples. The second principal component PC2 accounting for 24.9% variance accounts for a smaller fraction of MeDIP enrichment as well as differentiates C-term MeDIP from F-len and N-term MeDIP. The combined variance of PC1 and PC2 (97.7% of total variance) does not differentiate F-len and N-term showing that these two samples are highly similar overall.

**Fig S3.**
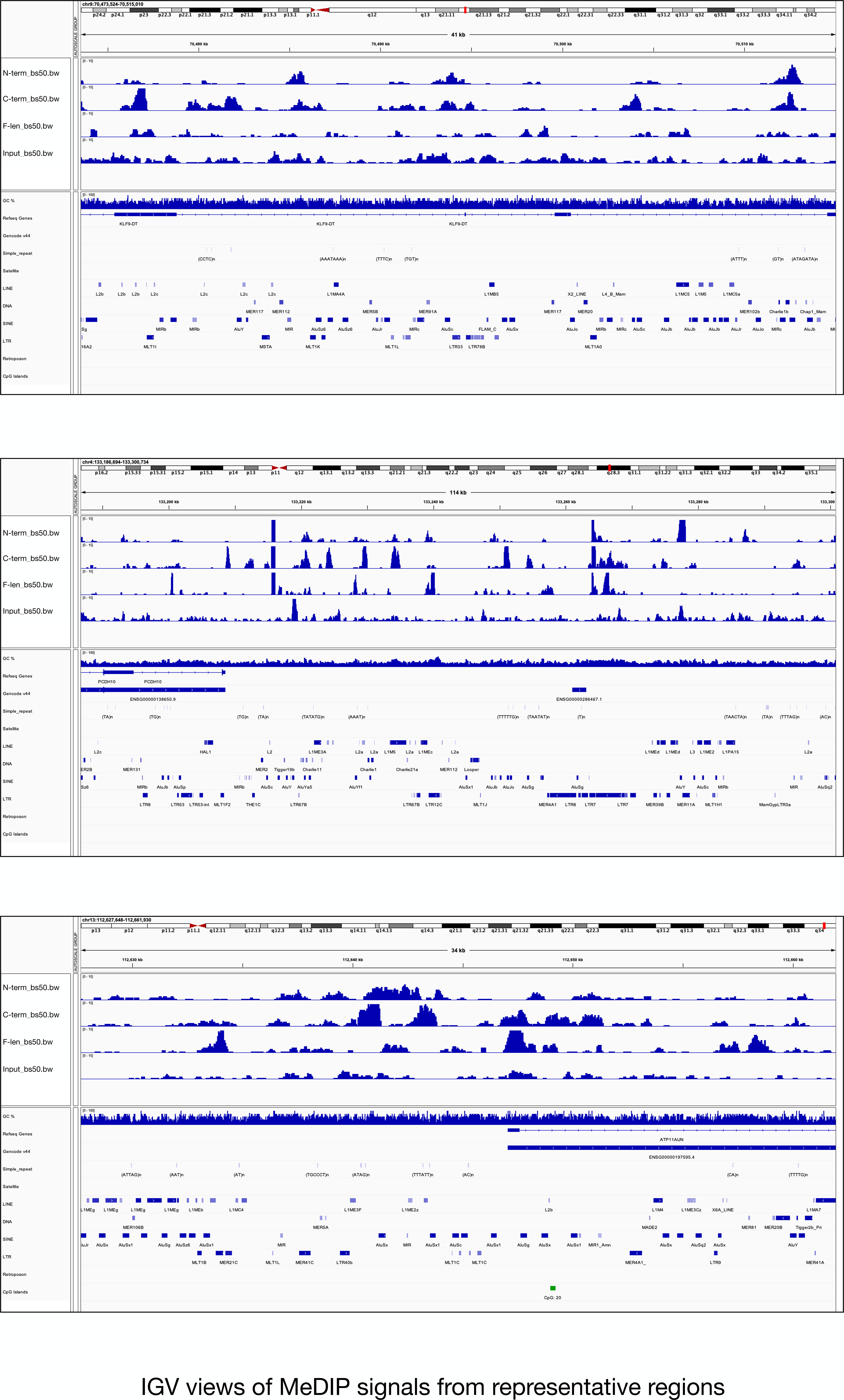
IGV views of MeDIP signals from representative regions: MeDIP signals from three selected representative regions allow a visual overview of the cytosine methylation patterns in the different samples C-term, N-term, F-len, input and some relevant annotation tracks. The regions shown are mentioned in the top left corner of each image.

**Fig S4.**
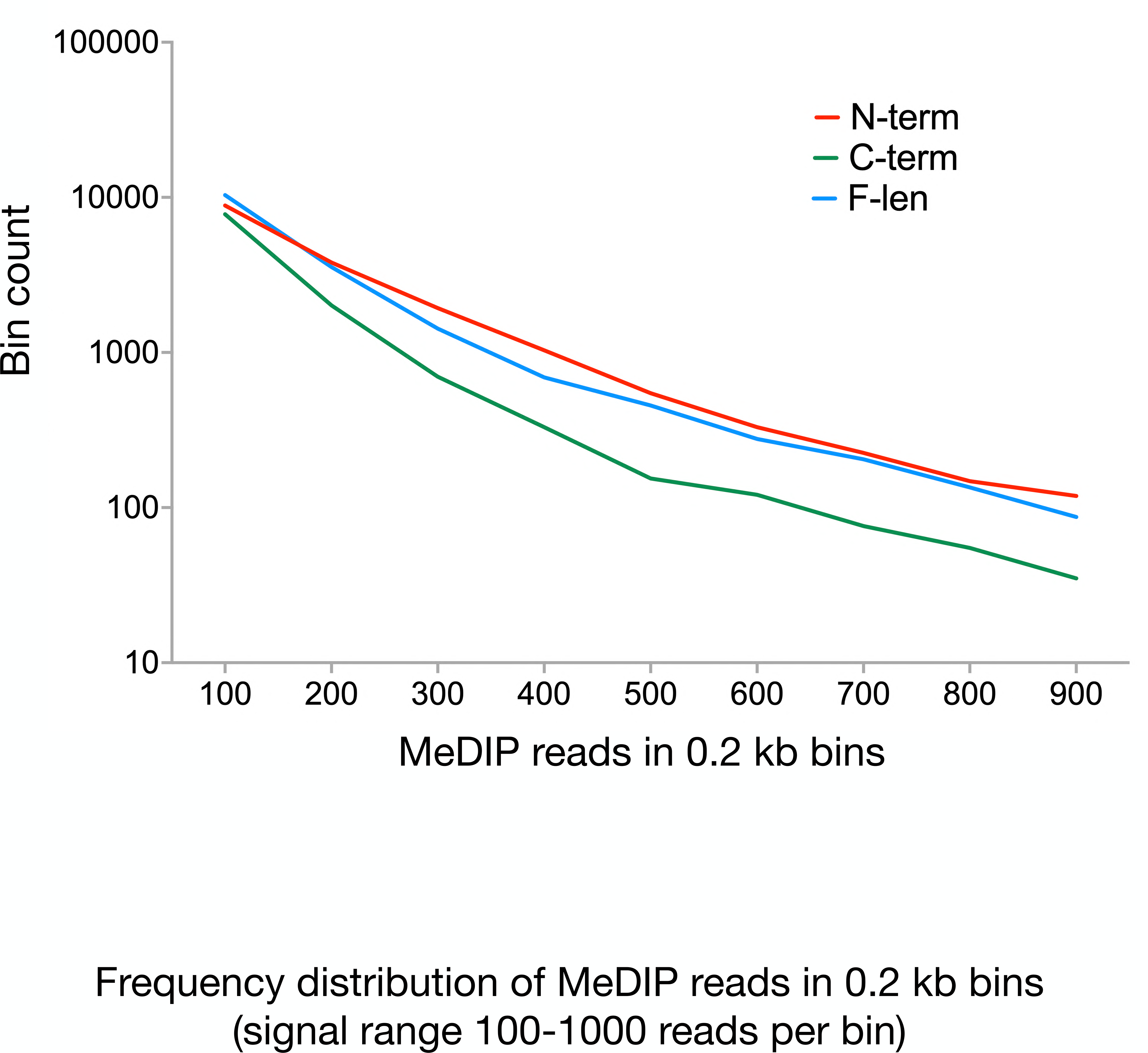
Frequency distribution of MeDIP reads in 0.2 kb bins (signal range 100-1000 reads per bin): MeDIP signal distribution genome-wide shows that the concentrated occurrence of higher levels of cytosine methylation is lower in C-term as compared to F-len and N-term. For a large range of MeDIP signals the F-len and N-term show no appreciable difference in cytosine methylation distribution. The increment of cytosine methylation levels in C-term hence occurs at regions with low cytosine methylation levels.

**Fig S5.**
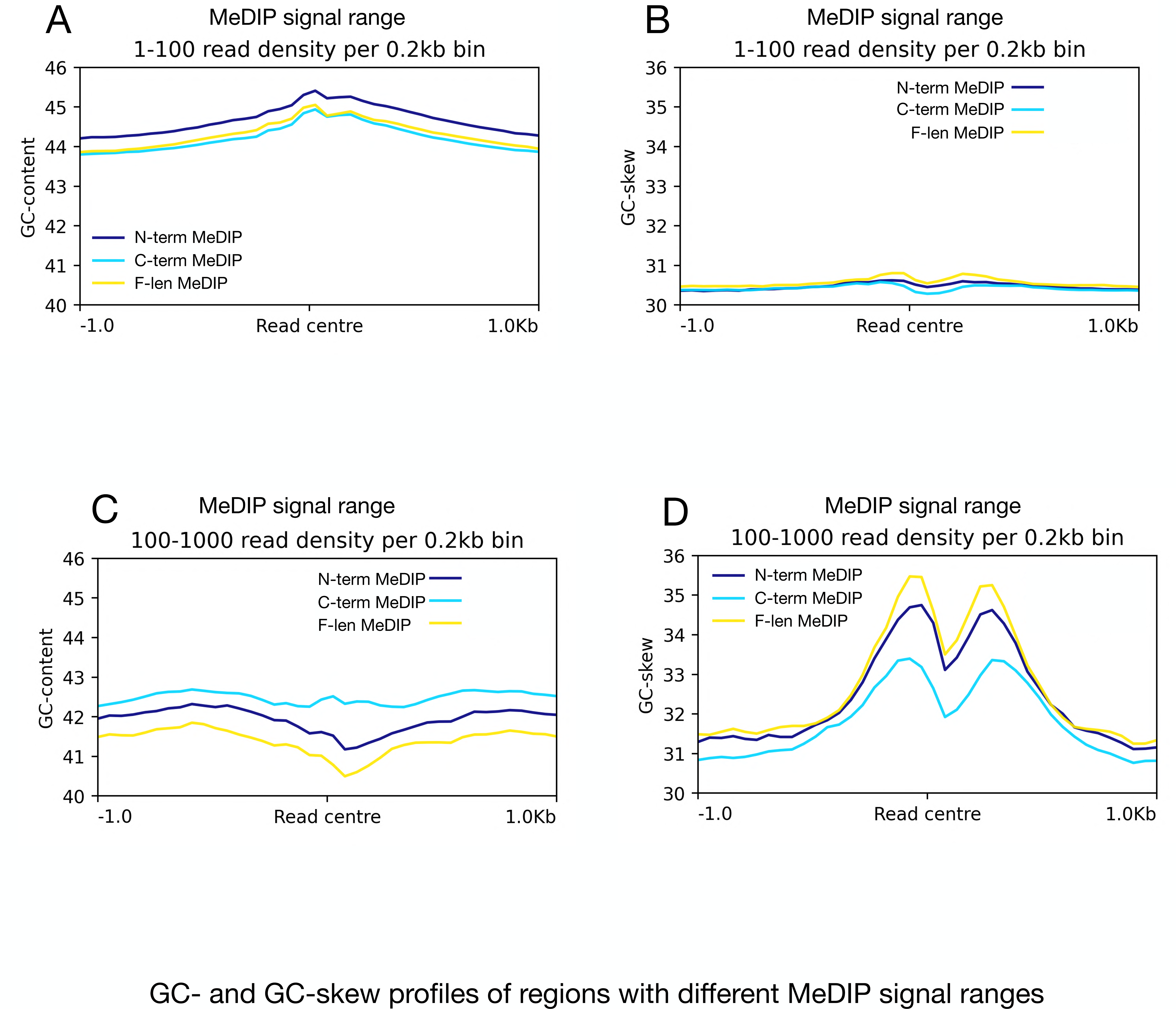
GC-content and G/C-skew of MeDIP reads mapping to regions with different cytosine methylation levels: GC-content at low cytosine methylation level regions with MeDIP signals 1-100 reads per 0.2 kb (A), G/C-skew at low cytosine methylation level regions with MeDIP signals 1-100 reads per 0.2 kb (B), GC-content at high cytosine methylation level regions with MeDIP signals 100-1000 reads per 0.2 kb (C), G/C-skew at low cytosine methylation level regions with MeDIP signals 100-1000 reads per 0.2 kb (D). The comparison of profiles presented in A-D show that highly methylated regions in C-term have high GC-content and low G/C-skew than those of N-term and F-len.

**Fig S6.**
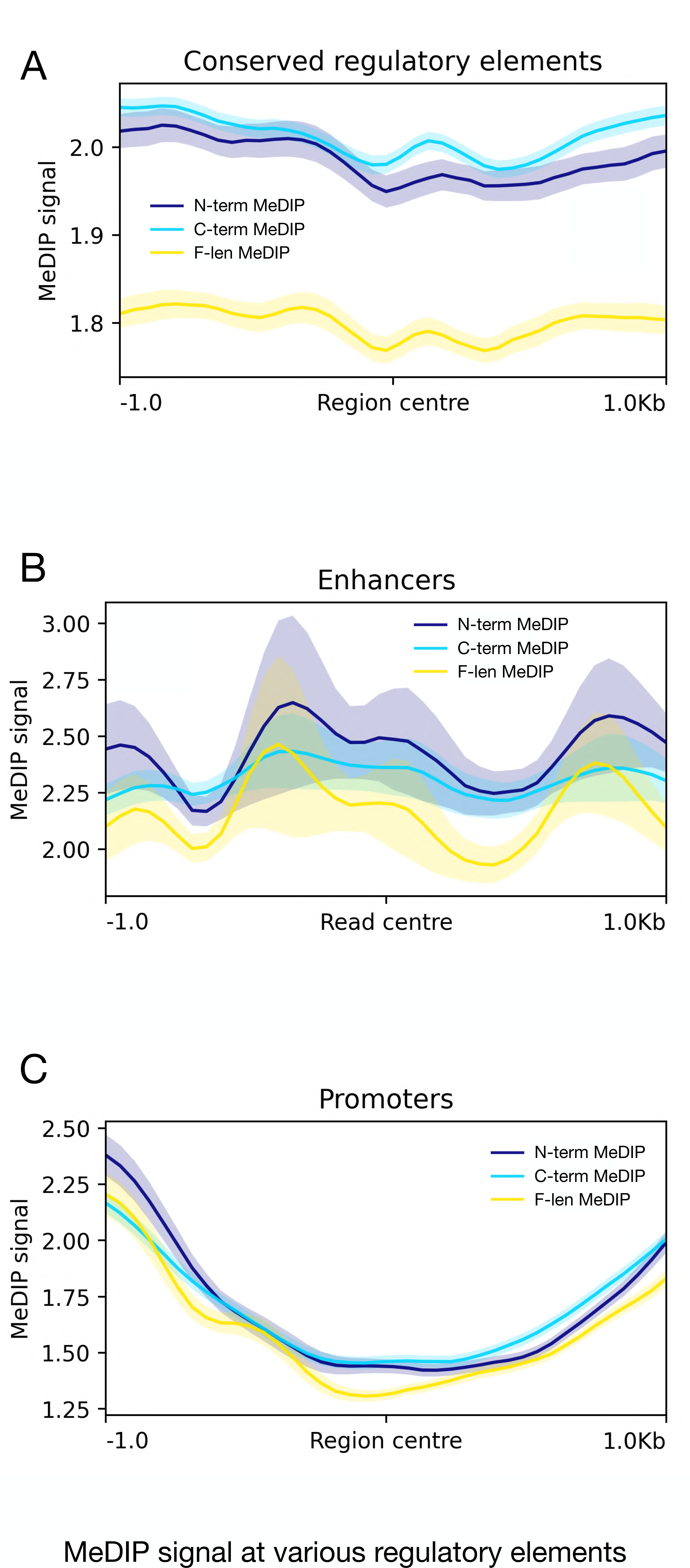
MeDIP signals at conserved regulatory elements, enhancers and promoters show mild stochastic differences between N-term, C-term and F-len. The most consistent effect of CGGBP1 loss of function, both C-terminal loss or N-terminal loss similarly affect cytosine methylation at conserved elements and increase it consistently. These findings are in agreement with the proposition that these conserved regulatory elements are maintained at low cytosine methylation by CGGBP1 through a mechanism that requires N-term as well as C-term.

**Fig S7.**
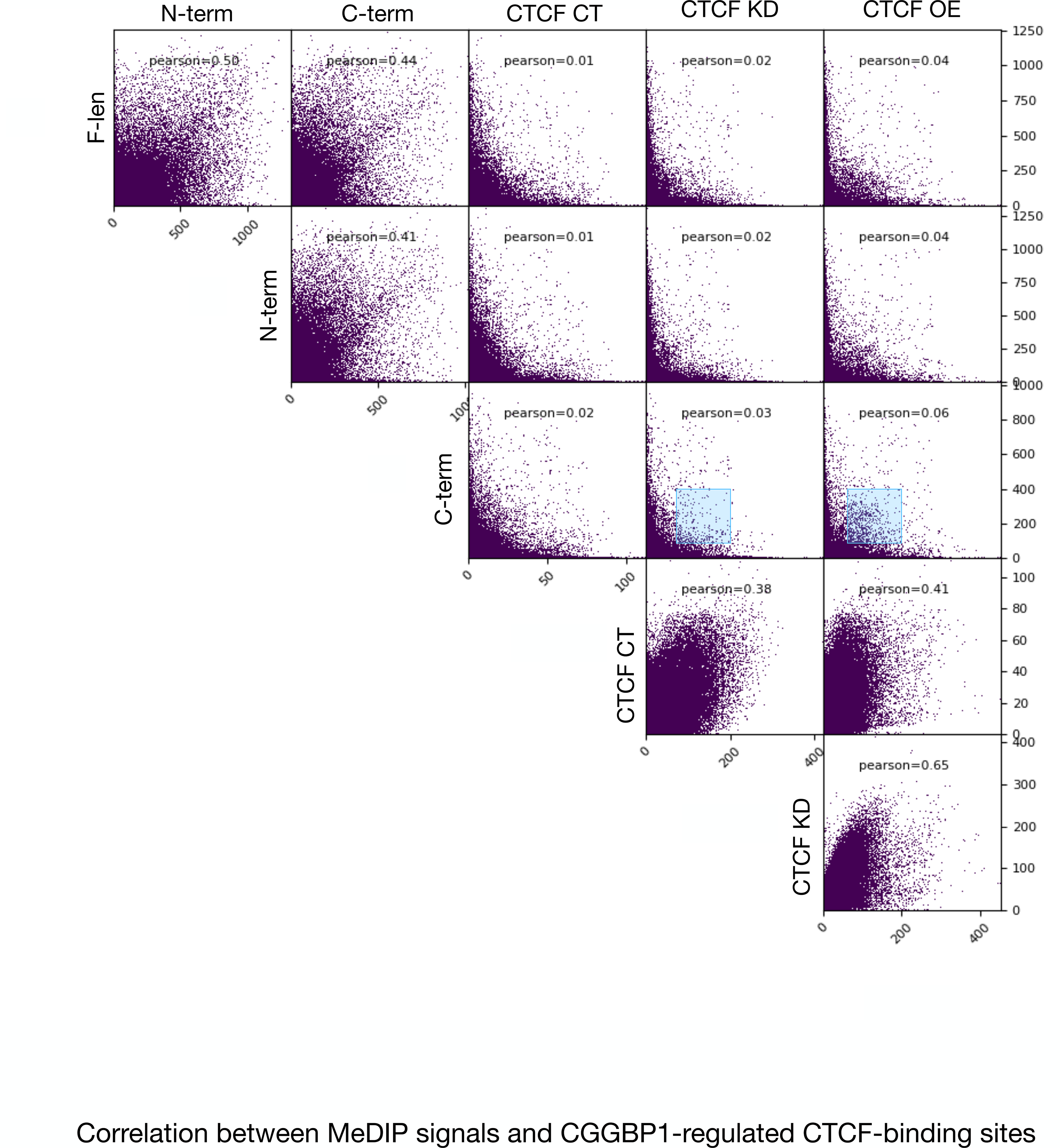
Correlation between MeDIP signals and CGGBP1-regulated CTCF-binding sites: A scatter plot showing a comparison of MeDP signals genome-wide in 50 bp genomic bins shows the effects of different truncations of CGGBP1 on cytosine methylation genome-wide as well as at regions where CTCF-occupancy is regulated by CGGBP1. The inset in each scatter plot shows the Pearson coefficients. The X-axis sample identities are mentioned at the top of the columns and the Y-axis sample identities are mentioned at the left of each row.

**Fig S8.**
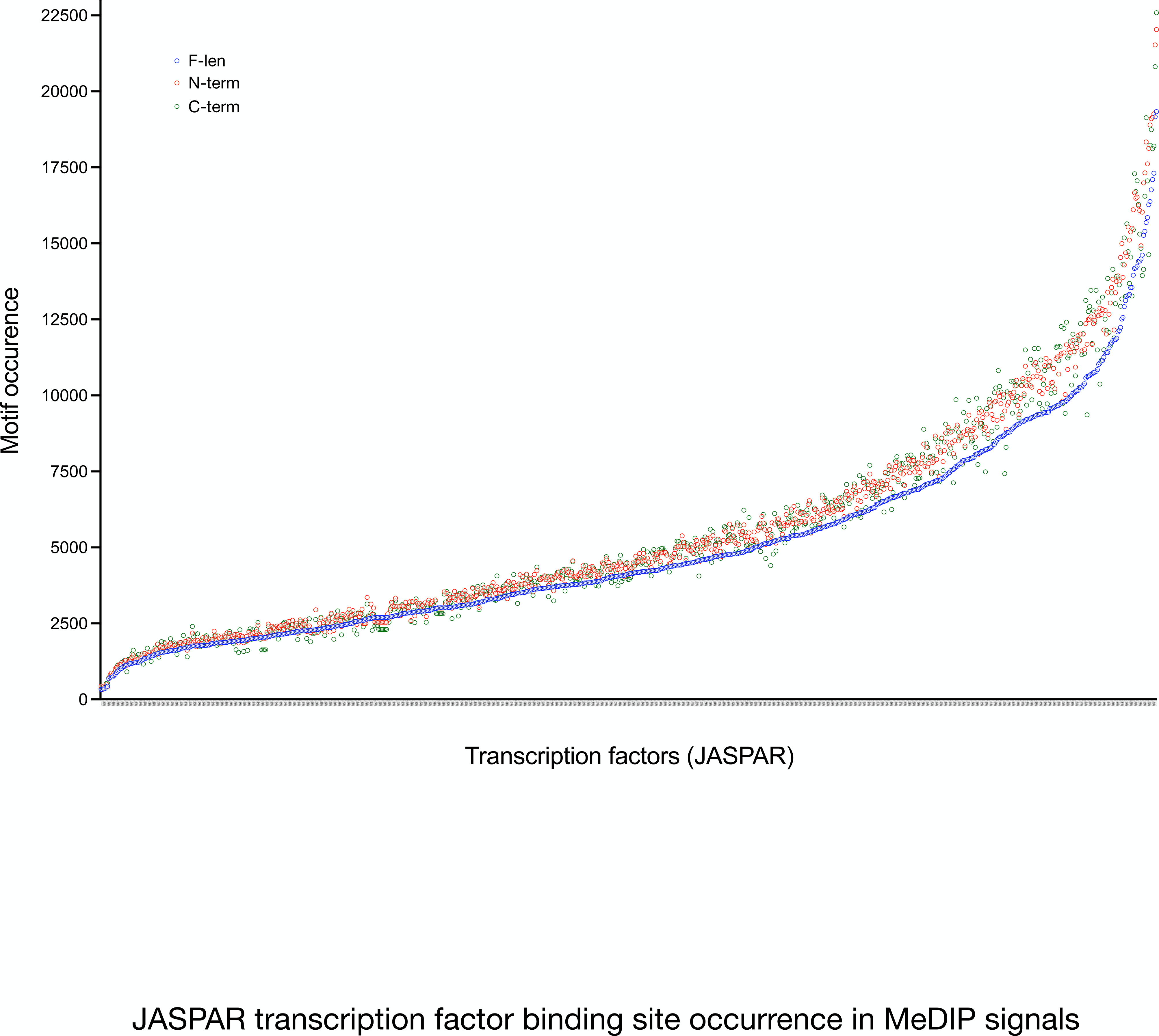
JASPAR transcription factor binding site occurrence in MeDIP signals: A motif search for JASPAR motifs in N-term, C-term and F-len MeDIP reads shows that for most transcription factor binding sites the occurrence in MeDIP reads (and hence their cytosine methylation) is lowest in F-len. With some exceptions for which C-term has lower cytosine methylation, the majority of transcription factor binding sites showed low cytosine methylation in F-len suggesting that F-len CGGBP1 maintains low level of cytosine methylation at transcription factor binding sites. The data points are sorted according to increasing order of binding site occurrence in F-len. The corresponding transcription factor binding site motif identifier is indicated along the X-axis.

**Fig S9.**
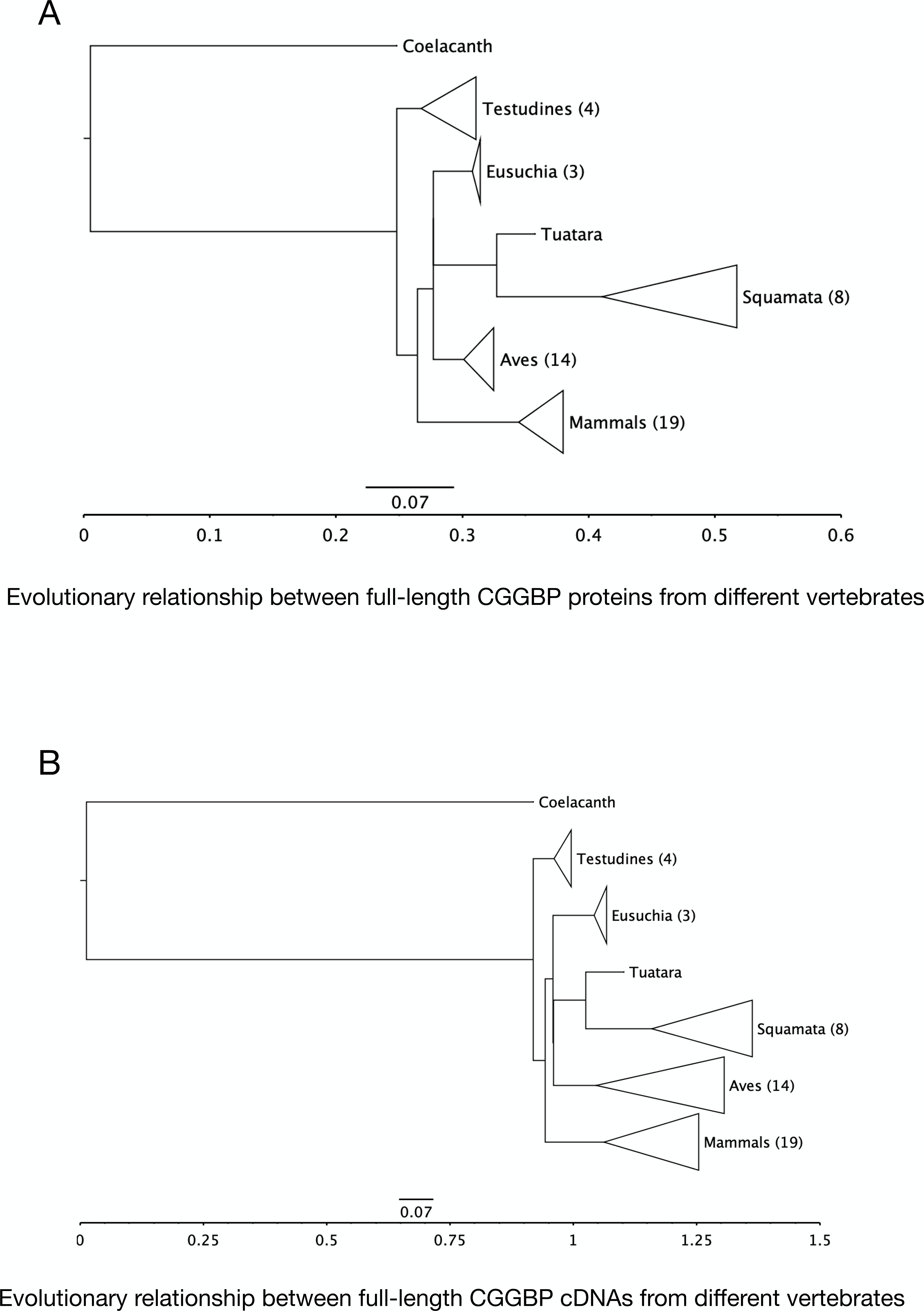
Evolutionary relationship between full-length CGGBP proteins and corresponding cDNA sequences from different representative vertebrates: Evolutionary trees of CGGBP proteins (A) and cDNA (B) sequences shows that the mammalian CGGBP1 is likely derived from the common amniote ancestor (except testudines). The trees show that reptilian CGGBP forms are highly divergent. It is noteworthy that in testudines and squamates the proteins are more diverse even if their cDNA sequences are less diverse. In aves and mammals the cDNA sequences are more diverse but the protein forms are less diverse suggesting a purifying selection in homeotherms. All the trees are rooted at coelacanth.

**Fig S10.**
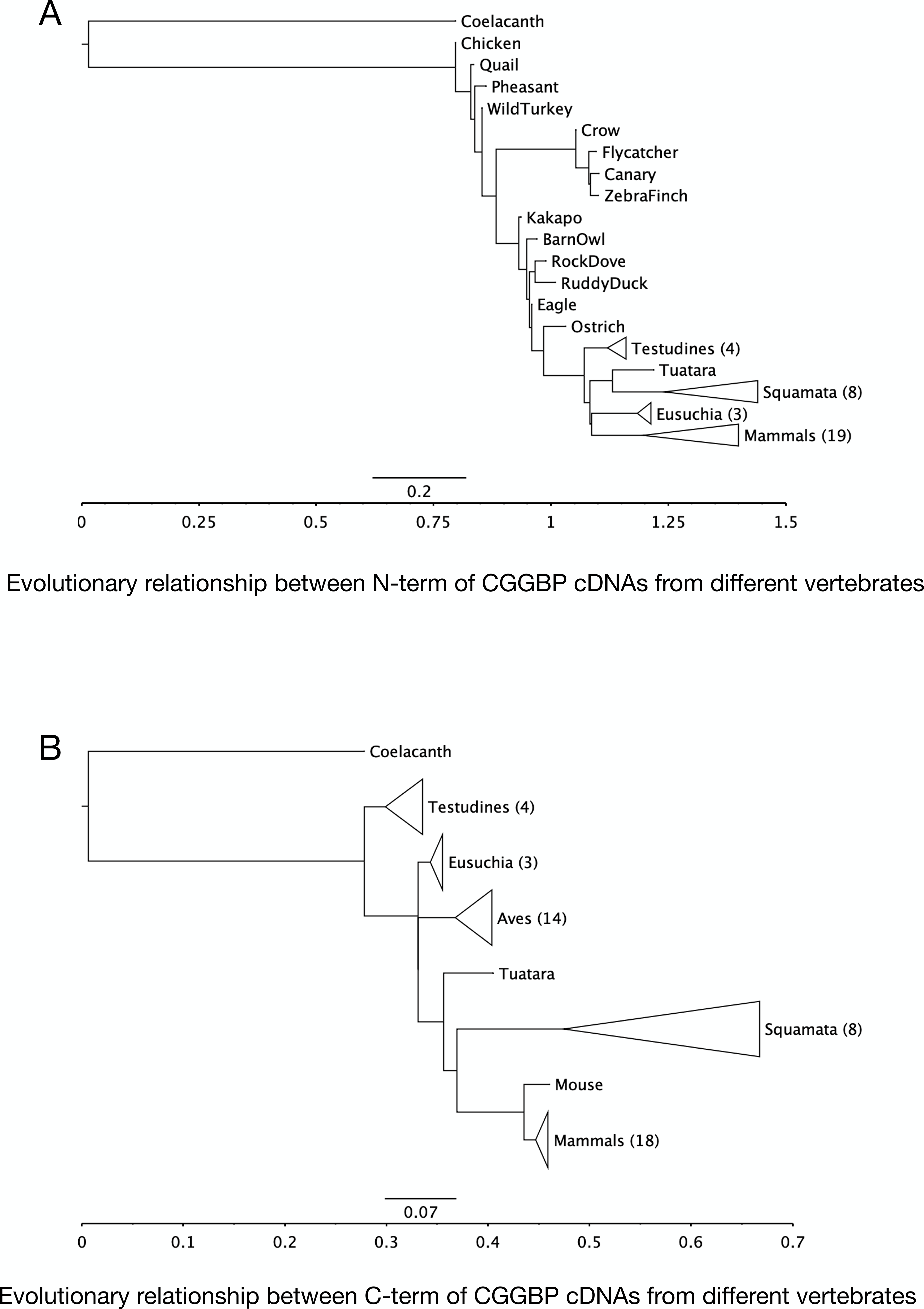
The phylogenetic relationship between the N-term and C-term cDNA sequences: The N-term cDNA (A) and C-term cDNA(B) sequences have accumulated changes such that there is no common amniote ancestor with distinct reptilian, avian and mammalian lineages as seen for F-len CGGBP1 cDNA (Fig S9B) from the same species. Interestingly, the divergent changes in C-term cDNA (B) affects the amino acid sequences and its phylogenetic relationship with other vertebrates (Fig 2C). However, larger divergent changes in N-term cDNA (A) do not affect its phylogenetic relationship with other vertebrates and the mammalian CGGBP1 N-term retains its origin from the common amniote ancestor (Fig 2B).

**Fig S11.**
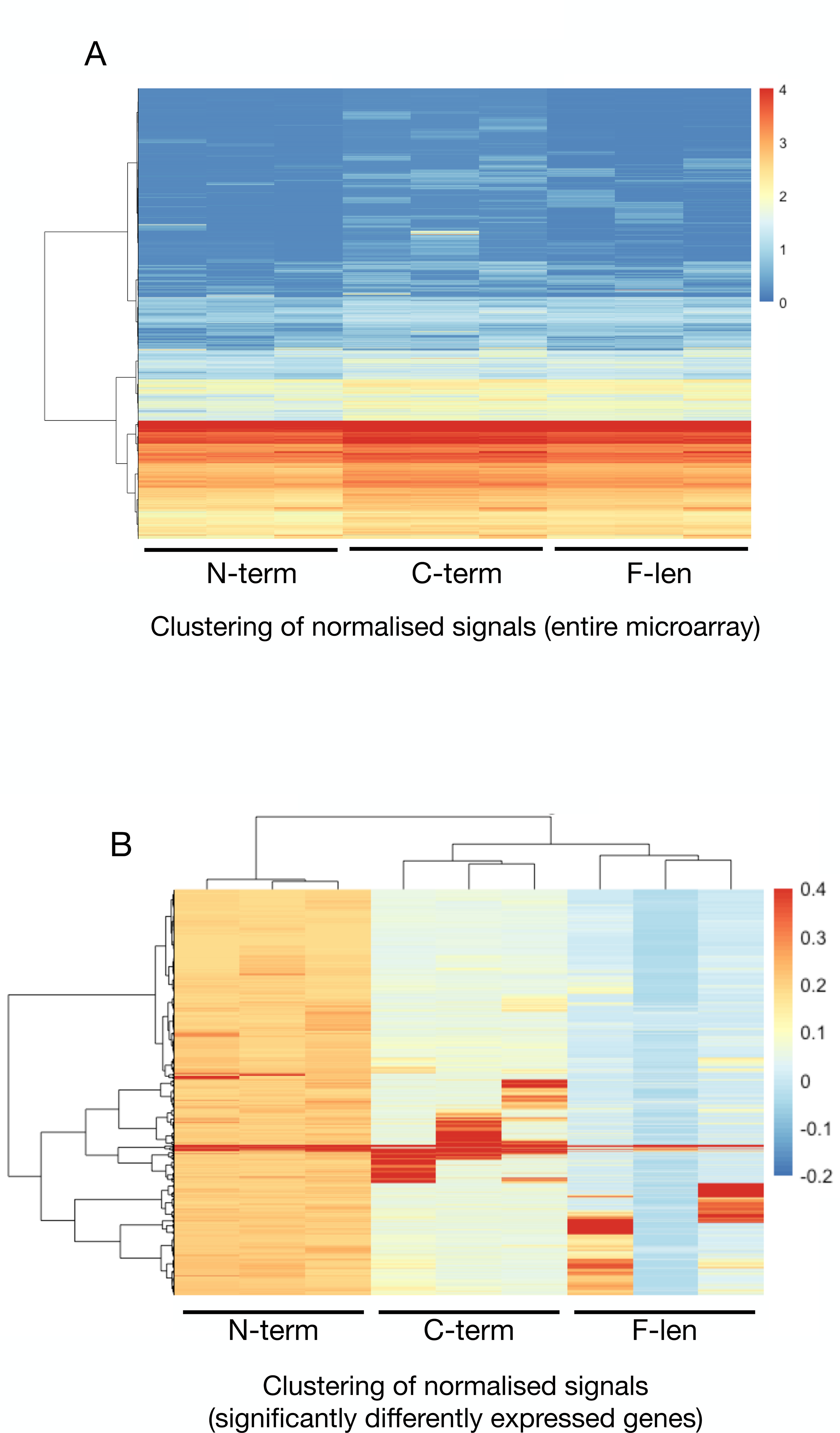
Gene expression deregulation by CGGBP1 truncation mutations: Heatmap and unsupervised clustering of normalised signals of reporters from three replicate experiments on C-term, N-term and F-len show that for >58K genes (A) the expression levels are highly consistent. For deregulated genes (p<0.01, n=3) the unsupervised clustering segregates the three sample groups distinctly. Strikingly, F-len and C-term act as repressors whereas N-term shows a derepression of gene expression.

**Fig S12.**
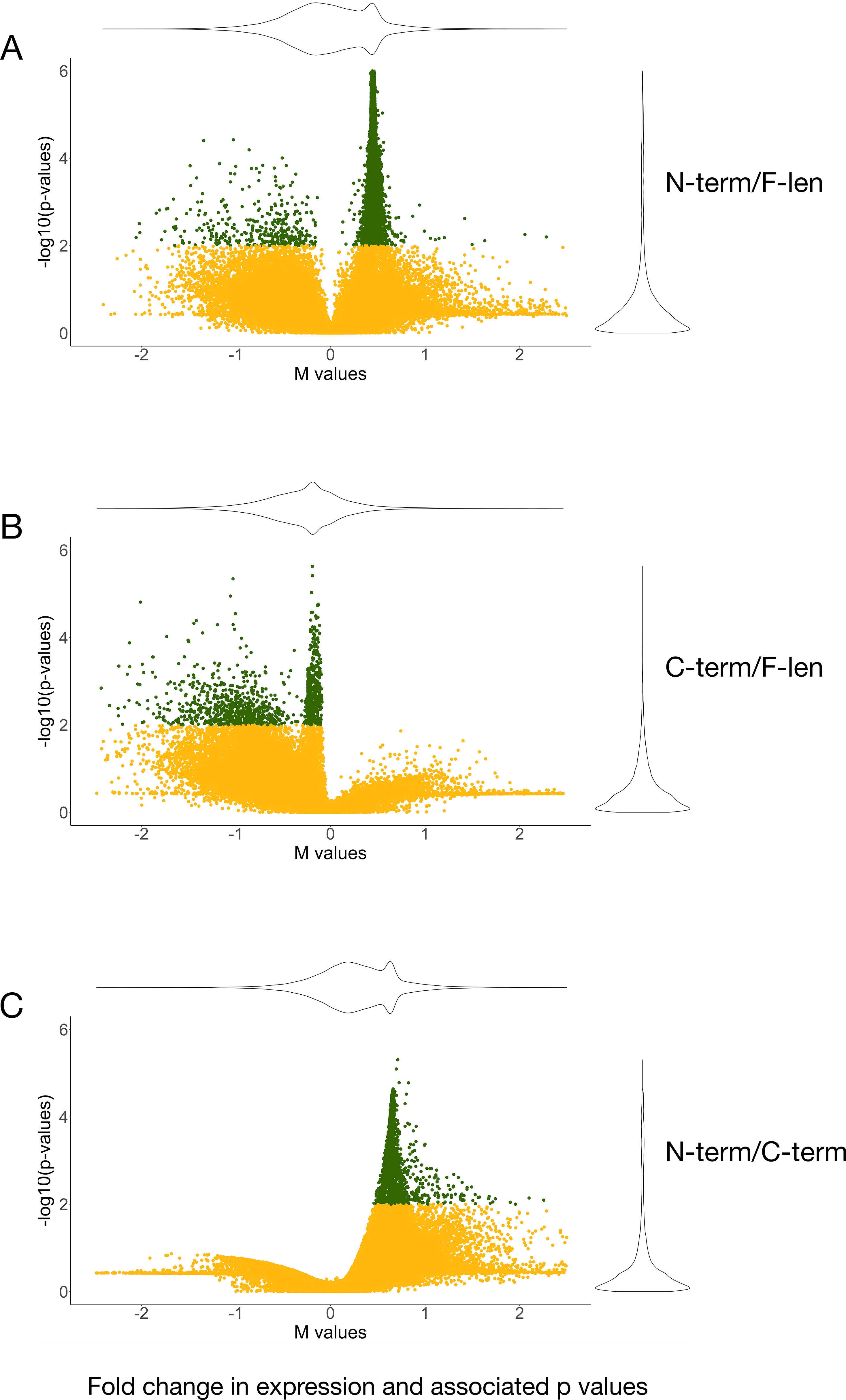
Volcano plots (significance of differential expression versus differential expression) of different sample pairs highlight the transcription repressive effect of F-len (A) which is retained in C-term (B and C). The F-len and C-term repressed genes also have high significance of differential expression (p<0.01 highlighted in green). The violin plots on the right and top of the plots show frequency distribution of the reporters.

**Fig S13.**
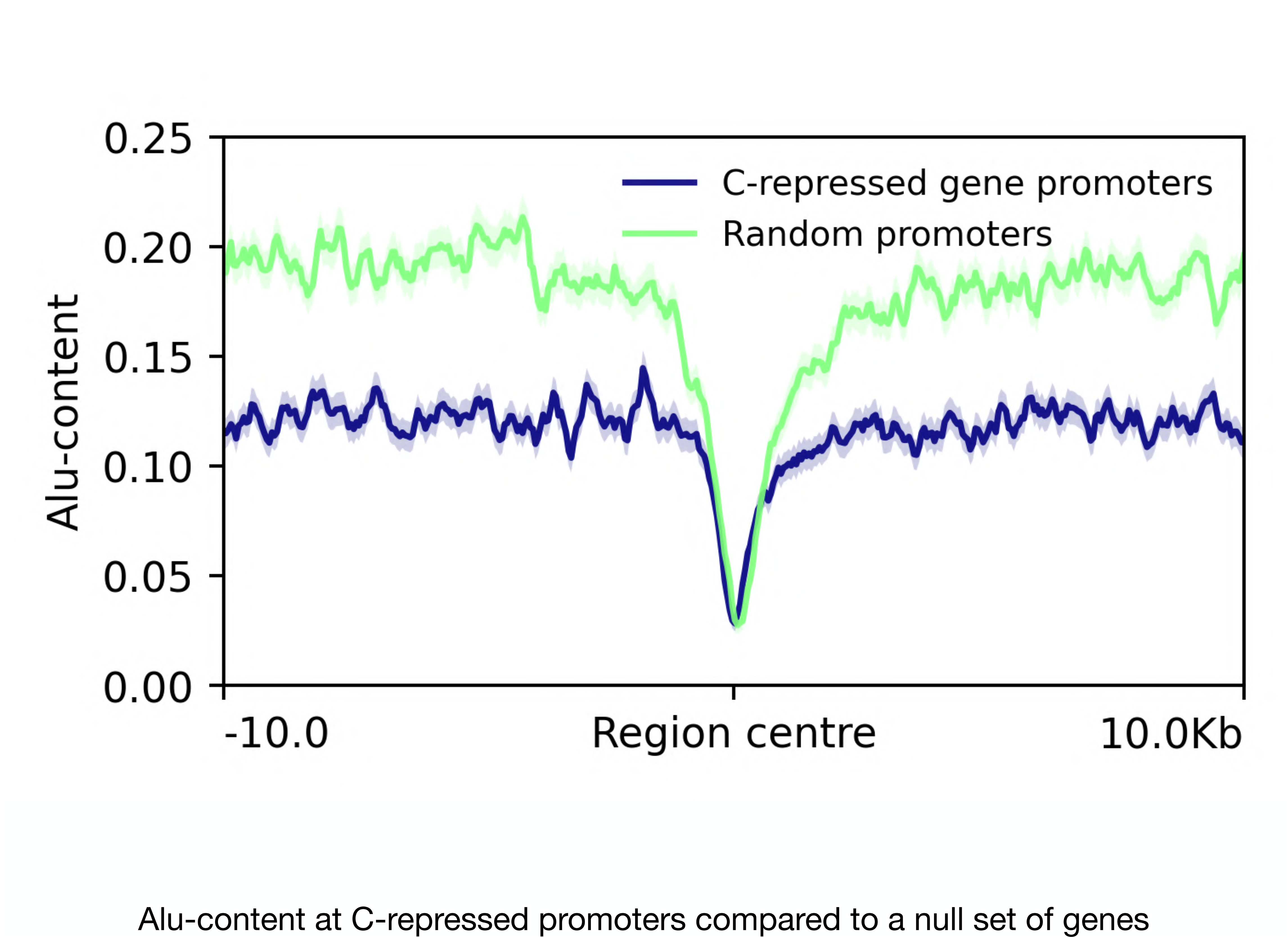
The C-repressed genes are localised in GC-poor and consequently Alu-poor neighbourhoods. However, even if the flanking regions are Alu-poor, the epicentre of MeDIP signals are at Alu-devoid regions.

**Fig S14.**
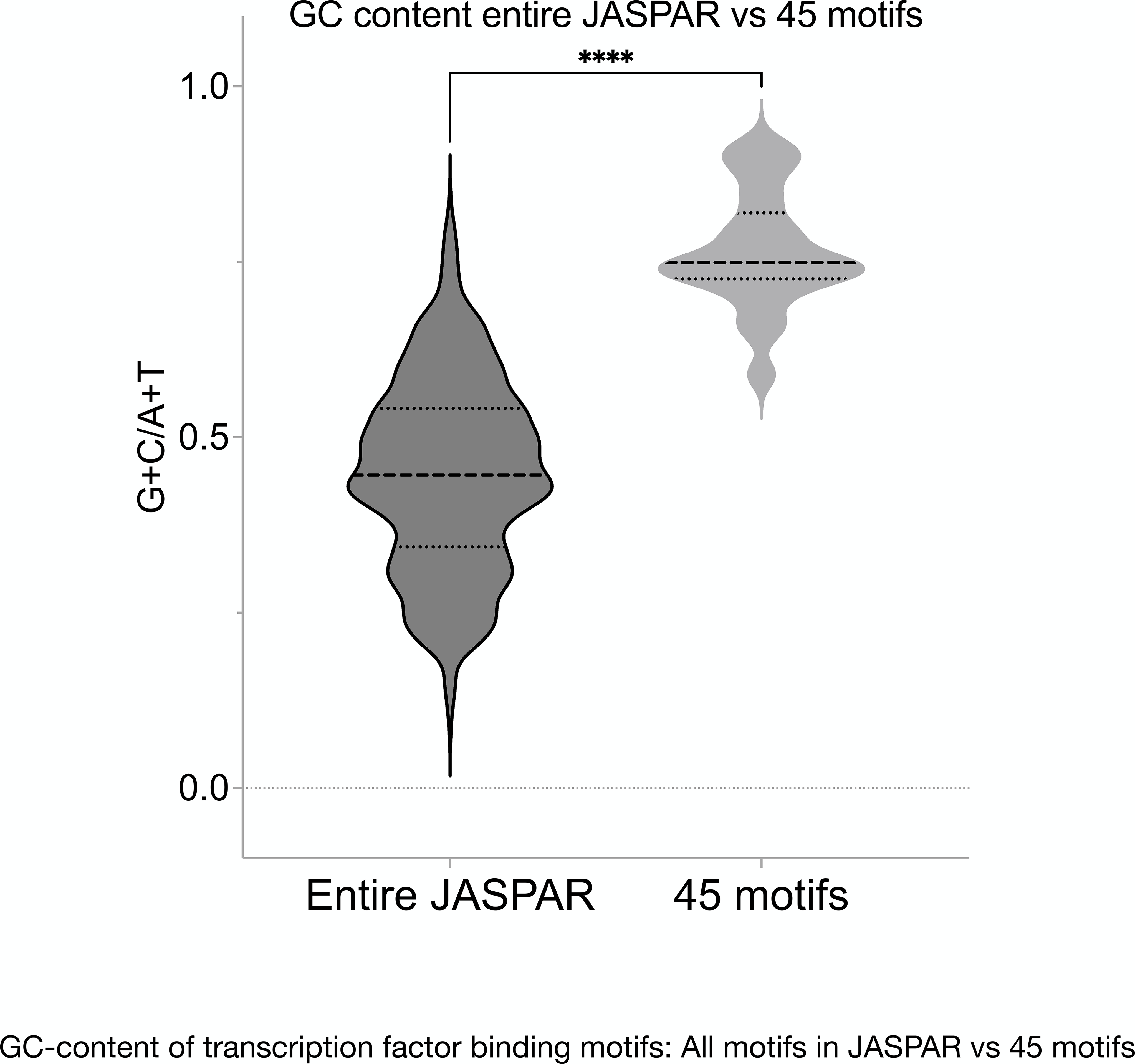
The GC-content of the 45 motifs is significantly higher (p value <0.0001) than the rest of all the JASPAR transcription factor binding motifs. The dotted lines represent the 25th and 75th percentile and the dashed line represents the median.

